# Click. Screen. Degrade. A Miniaturized D2B Workflow for rapid PROTAC Discovery

**DOI:** 10.1101/2025.07.29.667383

**Authors:** Marko Mitrović, Francesco Aleksy Greco, Yiliam Cruz García, Aleksandar Lučić, Lasse Hoffmann, Rohit Chander, Julia Schönfeld, Nick Liebisch, Martin Peter Schwalm, Markus Egner, Max Lewandowski, Daniel Merk, Viktoria Morasch, Elmar Wolf, Susanne Müller, Thomas Hanke, Ewgenij Proschak, Kerstin Hiesinger, Stefan Knapp

**Affiliations:** Institute of Pharmaceutical Chemistry, Goethe University Frankfurt, Max-von-Laue-Str. 9, 60438 Frankfurt am Main, Germany; Structural Genomics Consortium (SGC), Buchmann Institute for Molecular Life Sciences (BMLS), Max-von-Laue-Str. 15, 60438 Frankfurt am Main, Germany; German Cancer Research Center (DKFZ), Im Neuenheimer Feld 280, 69120 Heidelberg, Germany; Institute of Biochemistry, University of Kiel, Rudolf-Höber-Str. 1, 24118 Kiel, Germany; Department of Pharmacy, Ludwig-Maximilians-Universität (LMU) München, 81377 Munich, Germany; Fraunhofer Institute for Translational Medicine and Pharmacology (ITMP), Theodor-Stern-Kai 7, 60596 Frankfurt/Main, Germany

**Keywords:** PROTAC, D2B, HTS, Click Chemistry, Targeted Protein Degradation

## Abstract

Targeted protein degradation is one of the fastest developing fields in medicinal chemistry and chemical biology. Despite significant development in assay technologies and inhibitor discovery, the development of PROTACs remains a challenging endeavor since rational design approaches remain widely elusive. Our workflow eliminates the rate-limiting step of classic synthesis, namely compound purification, and pairs it with high-throughput, semi-automated plate-based synthesis, and direct cellular assay evaluation. We applied this direct-to-biology approach to four diverse targets demonstrating the general applicability of this technology. PROTAC synthesis was realized by using the highly efficient copper-catalyzed azide-alkyne cycloaddition reaction. This simplified reaction setup allowed synthesis in the nanomole scale with reaction volumes as low as 5 μL. This high throughput approach enables the synthesis and testing of hundreds of PROTACs within a few days, allowing for a comprehensive assessment of target degradability and the identification of the most suitable E3 ligase for degrader development.

## INTRODUCTION

Targeted protein degradation (TPD) has emerged as a new and promising, and potentially transformative pharmacological modality, receiving increased focus from researchers in academia and industry. Since the first reports of PROteolysis TArgeting Chimeras (PROTACs) in 2001, more than 30 degrader molecules have entered clinical development for diverse oncology indications.^1–3^ The general architecture of a heterobifunctional PROTAC molecule comprises a ligand that targets a given E3 ligase and a warhead that binds to the protein of interest (POI) connected via a linker region. A ternary complex is formed between the PROTAC, the POI and the E3 ligase, which is poised to catalyze the transfer of ubiquitin onto the POI, thereby tagging it for subsequent degradation by the ubiquitin-proteasome system (UPS).^4,5^Unlike conventional small molecule inhibitors, which rely on druggable binding pockets and require high inhibitor occupancy at the targeted binding site, degraders function catalytically via an event-driven mode of action. Additionally, the ligands used in the development of PROTACs do not need to bind to sites relevant to the disease-causing process of the POI. This has greatly expanded the druggable proteome.^6^

However, one of the main challenges for the development of PROTACs relies on the lack of rational design strategies. Furthermore, due to the increased structural complexity associated with this class of molecules, the synthesis and purification of the compounds remain the rate-limiting steps in the development of degraders (**Figure 1A**). Although critical steps for PROTAC activity have been highlighted,^7^ such as the stability of the ternary complex consisting of the PROTAC-mediated POI and the E3 complex, the design of PROTACs predominantly relies on combinatorial approaches that optimize linker properties and attachment points. This approach renders the development of PROTACs to be very time-consuming and resource-intensive. Thus, the development of PROTACs would benefit from methods that would allow for more streamlined approaches which reduce the time from synthesis to biological evaluation. Direct-to-biology (D2B) has emerged as a promising strategy that eliminates the need for purification by directly testing crude reaction mixtures in relevant assay systems.^8–11^

**Figure 1.**
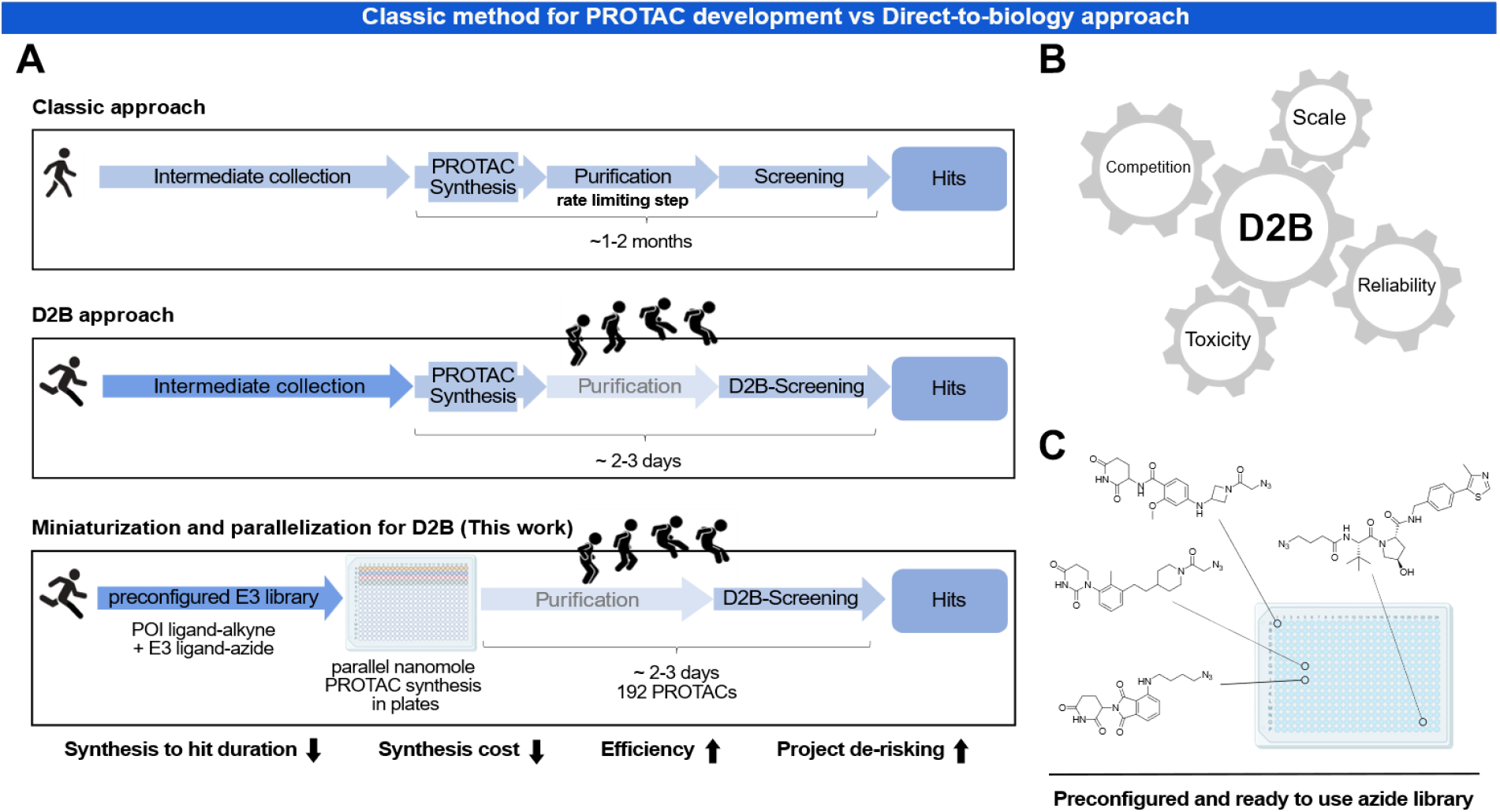
A Direct-to-biology platform for PROTAC synthesis: (A) Comparison between a conventional PROTAC development and a direct-to-biology (D2B) approach. (B) Different parameters that need to be optimized for a successful D2B PROTAC development including miniaturization. (C) A chemically diverse set of azides is transferred to 384 well plates that can be directly used in the Mini-Click synthetic approach.

In a D2B approach different factors need to be optimized in order to generate reliable cellular data. In fact, unreacted starting materials and reagents can affect the cellular readout as they can act both as competitors of PROTAC POI interaction and as cytotoxic agents, which can interfere with the assay. The reaction scale can become a key factor in achieving optimal reaction conversion, often limiting the useful volume and concentration ranges for high-throughput synthesis (**Figure 1B**).^12^

When paired with high-throughput synthesis (HTS) a D2B approach enables quick access to a large set of degrader molecules that can be characterized using appropriate sensor cell lines that monitor cellular POI concentration. Several luminescent detection systems are suitable for developing cellular assay including luciferase complementation assays such as HiBiT or assays using fluorescent tags, for instance GFP.^13^ Given the resource intensive nature of degrader synthesis, especially with varying linker moieties, a miniaturization approach would be expected to cut down synthesis costs. Herein, we report a high-throughput, nanoscale D2B synthesis approach for the rapid assembly of PROTAC molecules in a 384 well plate format using the highly efficient and robust copper-catalyzed azide-alkyne cycloaddition (CuAAC) with subsequent cell-based evaluation. This method allowed us to quickly address central TPD questions such as degradability, linkerology and the best combination of E3 ligase/POI ligand.

## RESULTS

### Synthesis of alkyne-bearing ligands for four different proteins of interest

To study the general applicability of the developed workflow, we conducted a proof-of-concept study examining a diverse set of four proteins. Target proteins from four different drug target families were selected including the well-established model system for PROTAC development Bromodomain-containing protein 4 (BRD4), soluble epoxide hydrolase (sEH), WD repeat-containing protein 5 (WDR5) and Aurora kinase A (AURKA). The targets vary in molar mass, protein family, function, structure and subcellular localization (**Table S1**). Since these proteins have been previously reported to be targeted by degraders, control PROTACs were available and have been used for benchmarking and as validation tools for our method.^14–17^ To enable the rapid assembly of PROTACs, validated high affinity inhibitors were combined with linkers harboring an alkyne moiety that was oriented towards the solvent region and acting as an exit vector. For each POI, four alkyne derivatives with varying linker composition with regard to length, polarity and rigidity were prepared. Alkyne (=A) derivatives obtained from the highly potent BET bromodomain inhibitor JQ1 targeting BRD4 (**Figure 2**, top left panel) as well as AURKA inhibitor MK-5108 (**Figure 2**, bottom right panel) were easily accessible to yield A1-A4 (BRD4) and A13-A16 (AURKA).^14,18^ For sEH, two exit vector strategies across three different chemotypes were introduced to the alkyne set: as previously reported, ligands A5 and A6 exit at the short branch and ligands A7 and A8 at the long branch of the L-shaped binding pocket of the hydrolase domain of sEH (**Figure 2**, top right panel).^19^ In the case of WDR5, ligands A9-A11, based on Dimethyl-F-OICR-9429-COOH (XF056-121) and A12, derived from the previously reported WDR5 chemical probe LH168, were chosen, which target the WIN-site pocket of WDR5 and share a similar strategy for linker attachment (**Figure 2**, bottom left panel).^15,20^

**Figure 2.**
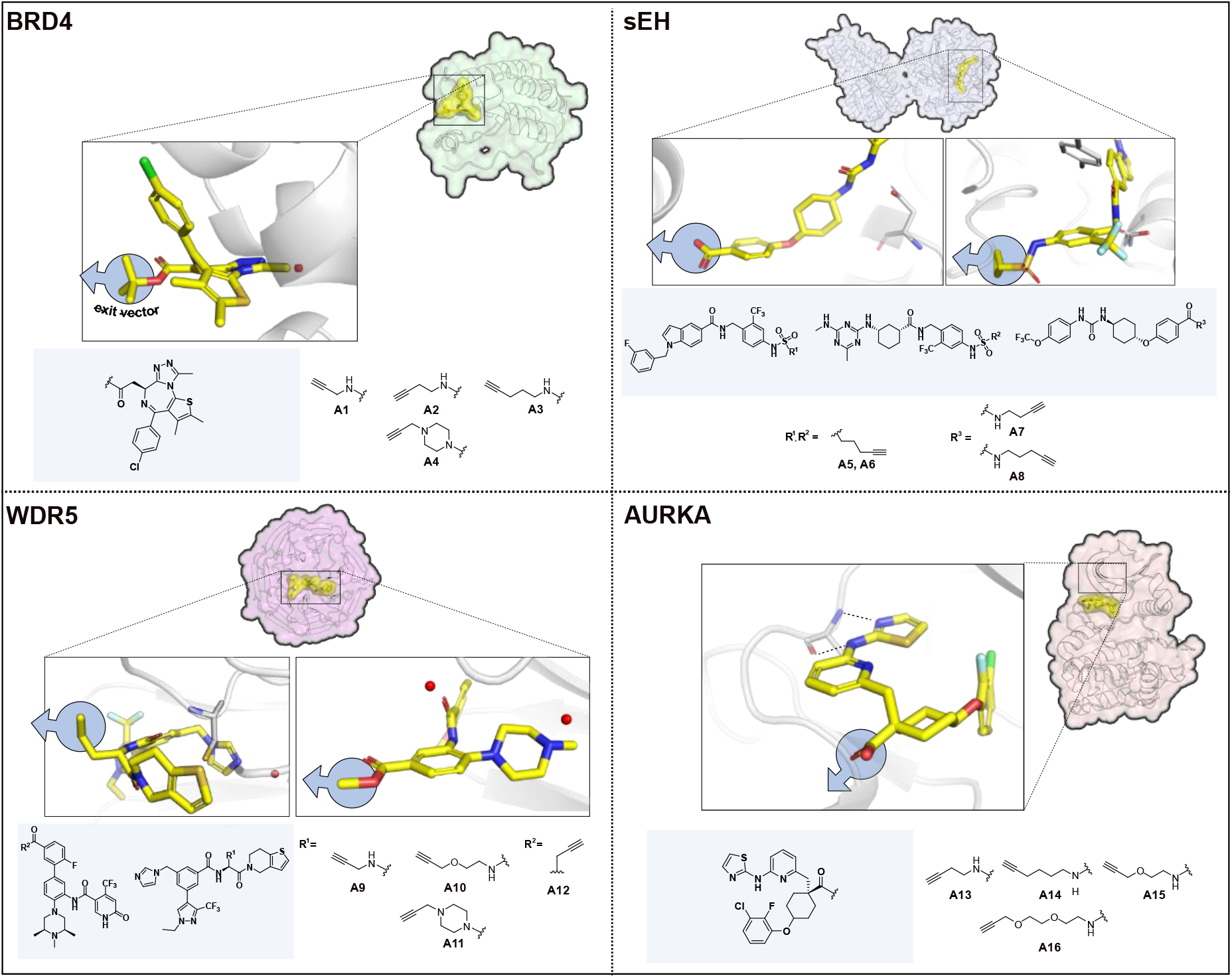
Overview of the proteins of interest used in this work: Four target proteins from different families were used in this study: Bromodomain-containing protein 4 (BRD4, top left), soluble epoxide hydrolase (sEH, top right), WD repeat-containing protein 5 (WDR5, bottom left) and Aurora kinase A (AURKA, bottom right) in complex with their corresponding ligands and exit strategies. The ligands were modified with various alkyne moieties and are shown next to the respective POI.

### Design and synthesis of the E3 ligand-linker library

Next, we synthesized a wide selection of E3 ligase ligands combined with various types and lengths of linkers bearing a terminal azide functionality, including alkyl, polyethylene glycol (PEG) and saturated heterocyclic moieties (**Figure 3**). The E3 ligase ligand core was either commercially available or prepared on a multigram scale and further modified to yield a total of 48 azides (X1-48), with 36 analogues linked to CRBN (thalidomide, phenyl dihydrouracil^21^ and benzamide^22^) and 12 ligands linked to VHL ligands. Straight forward synthetic routes including amide couplings and S_N_2 reactions were applied to rapidly assemble the azide building blocks, yielding the final azides at 50-150 mg scale. All synthetic procedures, NMR and mass spectra for active analogues and precursors are provided in the STAR Methods section. With four alkyne derivatives selected per POI ligand and 48 azides in hand, a total of 192 unique PROTACs were synthetically accessible for each target protein using CuAAC reactions.

**Figure 3.**
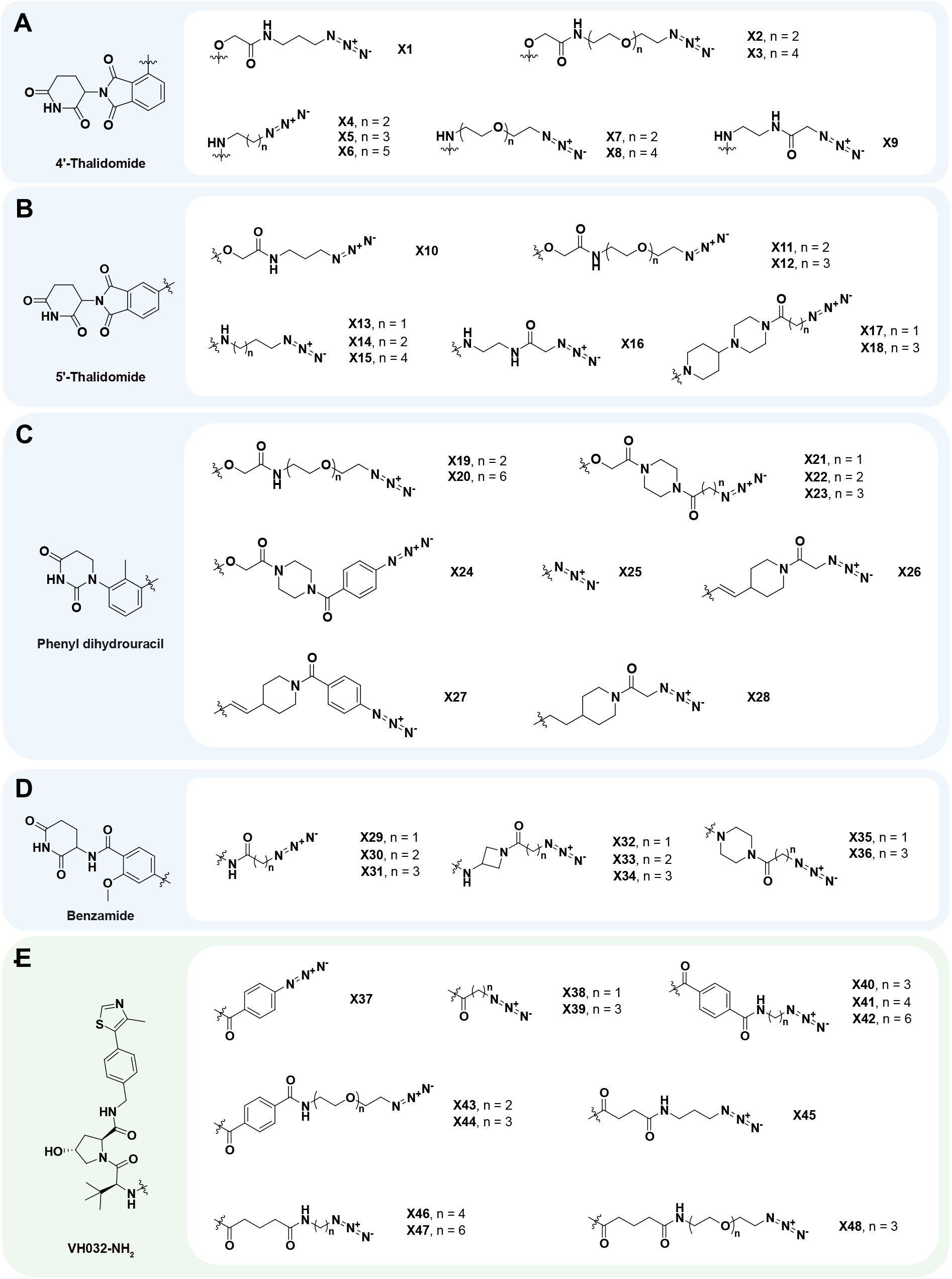
Overview of the modified E3 ligands used in this work: Chemical structures of the E3 ligase azide combinations (48 in total) targeting CRBN (blue shades) and VHL (green shade). (A) Nine derivatives based on 4’-substituted thalidomide scaffold, (B) nine derivatives based on 5’-substituted thalidomide scaffold, (C) 10 derivatives based on phenyl dihydrouracil (PDHU) scaffold, (D) eight derivatives based on benzamide scaffold and (E) 12 derivatives based on VH032-NH_2_ scaffold were synthesized.

### Optimization and Evaluation of the D2B platform on the first model system BRD4

To reliably apply our method, we sought to optimize the reaction conditions for the CuAAC by using BRD4 as a target protein. Since the biological evaluation of the PROTACs was performed in a D2B approach, the use of Cu^I^ as a catalyst and its associated effects on cell viability needed to be assessed.^23^ For this purpose, we examined different reaction conditions, modifying reactant concentration and volume, with the intent to lower the used Cu^I^ concentration (**Figure 4B and 4C**). The optimal and most reproducible reaction conditions were identified with reactant concentrations of 20 mM and a reaction volume of 5 μL. These conditions offered the best balance, providing the lowest possible scale, consistently resulting in high conversion rates, and the low Cu^I^ concentration had negligible influence on cell viability in the HiBiT assay. In addition, the specified reaction conditions enable longer use of the library, as on average only ∼ 50 μg of a single azide was utilized per reaction. Finally, we set the equivalent of alkyne and azide at 1 eq., the equivalent of CuSO_4_·5H_2_O and sodium ascorbate at 0.3 eq. and the reaction temperature at rt (**Figure 4A**). A total of 192 unique BRD4 PROTACs were synthesized after 24 h reaction time and the conversion rate of each well was determined by HPLC/UV analysis. Among the 192 reactions, 106 reached reaction conversions of more than 95% and an average conversion rate of 92% was achieved. This indicated the high reliability and broad scope of our method (**Figure 4D**). Before evaluating the degradation efficacy of the BRD4 PROTACs in a sensor cell line (HEK293T cells), a CellTiter-Glo® assay, with the components of an exemplary click reaction at different concentrations was performed to analyze cell viability. Encouragingly, no overall cell toxicity caused by the copper catalyst was observed even after 24 h of exposure with the reaction mixture, paving the way for subsequent direct POI degradation studies in cell-based assay systems (**Figure 4E**, dotted yellow lines represent the copper concentrations present in our intended assays conditions at a PROTAC concentration of 1 μM and 200 nM). We therefore proceeded to perform HiBiT lytic assay-based screening using the crude BRD4 PROTACs at 1 μM concentration. Examination of our assay data revealed that after 6 h, several PROTACs significantly decreased BRD4 levels, with 11 PROTACs achieving degradation above 50% (**Figure 4F**). Noticeably, among the different CRBN targeting chemotypes only the thalidomide based (X1-X18) PROTACs showed a significant degradation efficacy, indicating that the right choice of E3 ligase ligand scaffold has a crucial effect on the performance of the PROTAC. Next, eight PROTACs P1-8, ranging from high to low degradation efficacy in our D2B screen, were resynthesized at larger scale and purified to validate the hits generated with the developed plate based synthetic method (P1-P8, Data S1). Analysis of the re-synthesized PROTACs revealed degradation of BRD4 in cells in a dose dependent manner. Importantly, the determined PROTAC potencies of the purified degraders correlated well with degradation levels of the corresponding molecules synthesized and evaluated by our D2B approach (**Figure 4G**). Additionally, a HiBiT lytic assay comparing the purified and crude PROTACs P1-8, was carried out at 200 nM (**Figure 4H**). We observed a significant difference in degradation efficacy when comparing purified with crude PROTACs (∼ 25% difference). These data indicated that competition of PROTAC binding to the POI or E3 ligases with unreacted reactants in crude reaction mixtures, reduced sensitivity detecting degrader molecules. To confirm that the lower degradation efficacy was due to unreacted POI and E3 ligands, we designed “spiking experiments”, in which a pure PROTAC (P1) was added to a solution of the corresponding reaction components (alkyne A3, azide X15 and a mixture of A3 and X15 in 1:1 ratio), mimicking partial conversions of the CuAAC reaction (**Figure 4I**). The obtained degradation data revealed that the presence of just 10% unreacted alkyne decreased the degradation efficacy by half compared to pure PROTAC P1 (24% vs. 51% residual BRD4 levels). The effect of 10% residual azide was smaller but still noteworthy (24% vs. 30% residual BRD4 levels), probably due to weaker E3 ligand binding affinities. Nevertheless, based on the excellent conversion rates we obtained, the developed screening method was sensitive enough to identify 11 active BRD4 PROTACs with a degradation efficacy above 50%, resulting in an overall hit rate of approximately 6%.

**Figure 4.**
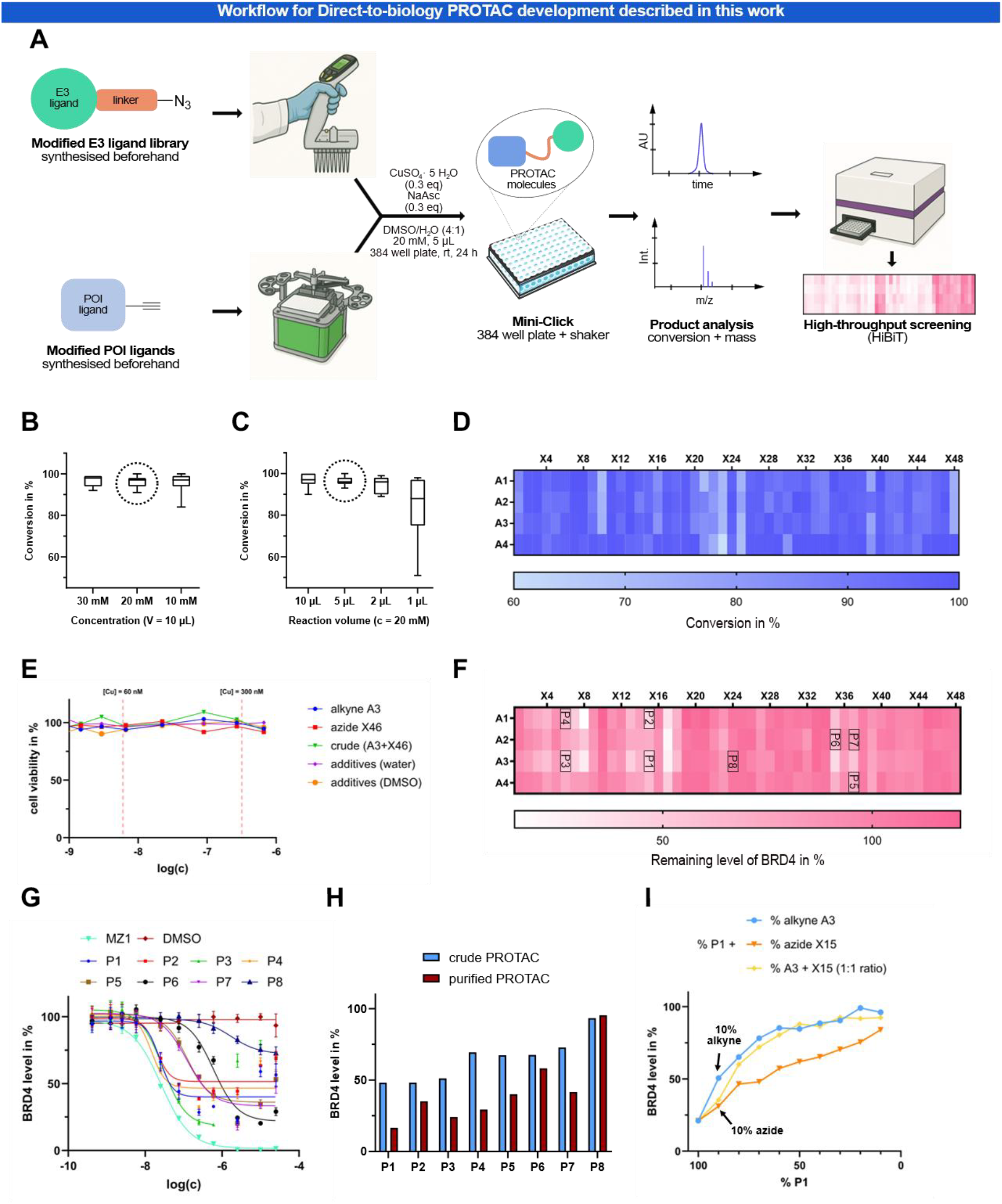
Assembly of the PROTAC library using CuAAC and identification of degraders from D2B screening: (A) General workflow of the developed, miniaturized PROTAC synthesis in 384 plates utilizing CuAAC described in this work: Library development including POI/E3-ligands derivatized with alkynes or azides, respectively. Automated and semi-automated dispensing for reaction set up. Product formation monitoring via HPLC-MS and hit identification using direct cell-based screening and automated data analysis. (B) Fluctuation of the overall conversion rate of the BRD4-targeting ligands with varying reactant concentrations (constant reaction volume = 10 μL). (C) Effects of reaction volumes on conversion rates (constant concentration= 20 mM). (D) Conversion rates of the CuAAC chemistry for BRD4 PROTACs, indicated by a color-coded heat-map based on HPLC UV data. (E) Cell viability data after 24 h with components used in an exemplary click reaction (CellTiter-Glo). (F) HiBiT lytic assay-based screening results for BRD4 at 1 μM after 6 h PROTAC exposure. (G) Dose response curves for the purified PROTACs P1-8 using the HiBiT detection system. MZ1 was used as a positive control (26035625). (H) HiBiT lytic assay data for purified BRD4 PROTACs P1-8 compared to the corresponding crude reaction mixtures after 6 h treatment, using a theoretical 200 nM PROTAC concentration assuming 100% conversion rates. (I) Spiking experiment for PROTAC P1 with alkyne A3 (blue), azide X15 (orange) and a mixture (1:1) of A3 and X15 (yellow) at 200 nM PROTAC concentration after 6 h exposure.

After successfully implementing the D2B platform with the model protein BRD4, we used the established protocol to identify active PROTACs for three additional structurally diverse proteins: sEH, WDR5 and AURKA. The goal of this project was twofold: first, to demonstrate the general applicability of our D2B platform; and second, to address key questions in PROTAC optimization, such as the roles of incubation times, linker attachment points and screening concentrations, as well as E3 ligase compatibility of a selected target.

### CRBN based PROTACs induce faster sEH degradation than VHL analogues

Targeting sEH, our CuAAC generated PROTACs were tested in a lytic HiBiT assay system established in HeLa cells at two different incubation times of 6 and 18 h, respectively. A projected PROTAC concentration of 200 nM was used. Applying a degradation threshold of 45% for sEH, we identified 10 highly active PROTACs and yielded an overall hit rate of 5%. Notably, the azide X8 exhibited degradation independent of the chemotype of the sEH ligand used (**Figure 5A**, left side, highlight in blue). In general, most potent sEH degradation was observed with alkyne A5 and CRBN-based azides. Within the same linker chemotype, longer linker moieties resulted in improved degradation potencies. For other alkynes (A6–A8), this trend was less consistent. Initially, no hits were detected with VHL-based azides after 6 h. However, at the 18-h time point, the combination of PROTAC A6 + X42 led to a 56% decrease of sEH levels (**Figure 5A and 5B**). Interestingly, VHL-based hits were exclusively observed using the POI ligand A6, underscoring the importance of utilizing diverse POI ligands when available (**Figure 5A**, right side, highlighted in blue). To date, no effective VHL-based PROTACs have been reported for sEH, and X42 may serve as a promising starting point for further development of VHL-recruiting sEH degraders. These results also highlight the value of screening across multiple time points to gain a more comprehensive understanding of degradation kinetics.

**Figure 5.**
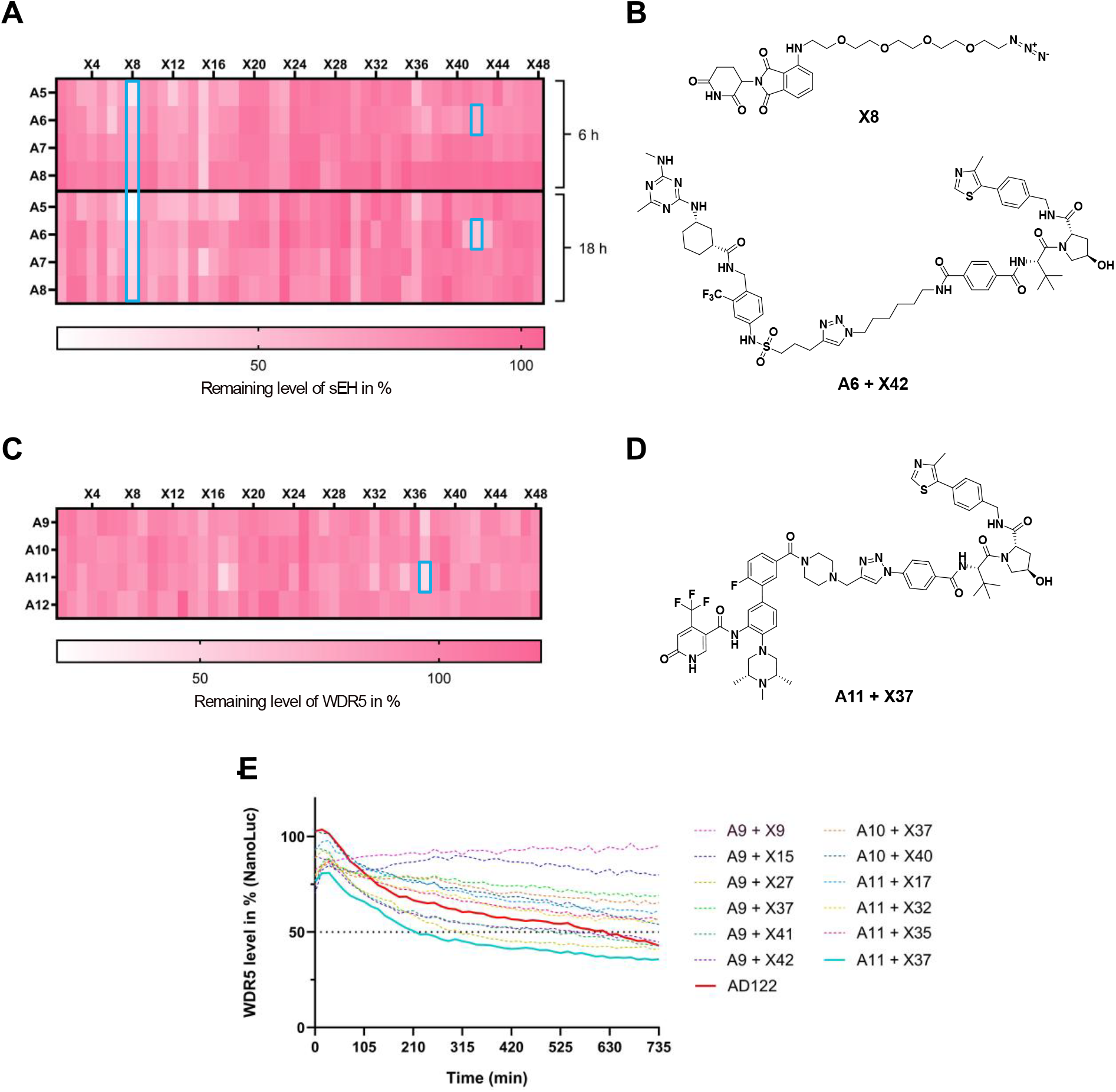
Identification of active PROTACs for sEH and WDR5: (A) HiBiT lytic assay-based screening data for sEH at a PROTAC concentration of 200 nM after 6- and 18-h incubation time. The best performing azide (X8) and the time dependent degradation effect of VHL containing PROTAC A6 + X42 are highlighted (blue). (B) Chemical structures of the azide X8 and PROTAC A6 + X42. (C) HiBiT lytic assay-based screening data for WDR5 at a PROTAC concentration of 1 μM after an incubation time of 6 h. (D) Chemical structure of the PROTAC A11 + X37 that induced the most efficient degradation of WDR5. (E) NanoLuc time dependent live cell measurement monitoring the degradation of WDR5 at a PROTAC concentration of 1 μM (crude reaction mixtures and AD122).

### Dimethyl-F-OICR-9429-COOH is the most efficient POI ligand

Next, we explored our D2B platform for a screen identifying degraders of the nuclear protein WDR5 for which we developed PROTACs previously and also established a HiBiT assay technology.^7,15^ In our screen for WDR5-targeting PROTACs, three PROTACs were identified that degraded WDR5 by more than 45% (**Figure 5C**). In line with previous results, we observed that degradation of WDR5 can be achieved with both CRBN- and VHL-based PROTACs.^15,24^ Among them, A11 + X37 emerged as the most potent hit (**Figure 5D**). Interestingly, the second ligand used in this study based on a recently developed ligand LH168^20^ yielded no hits, despite the overall excellent conversion rates observed for this ligand (**Figure S2**). This may be attributed to a different linker attachment point or the observed slow binding kinetics of this compound, potentially reducing the effectiveness of this ligand for PROTAC design.

In an orthogonal live-cell, NanoLuc luminescence assay in MV4-11 cells, A11 + X37 was confirmed as the most potent hit (**Figure 5E**), surpassing the reference PROTAC AD122^7,15^ during the 12 h incubation time. In general, more hits were identified using this screening method, possibly due to a higher sensitivity of this assay system.

### AURKA is a highly degradable kinase for CRBN but not VHL based PROTACs

The last case study focused on the mitotic kinase AURKA. Also, for this POI, we previously identified PROTACs using traditional PROTAC development strategies.^14,18^ In addition, AURKA has been identified as a highly “degradable” kinase using non-selective kinase inhibitors making it an excellent model system for validation of our platform.^25^ Indeed, when screening the generated D2B PROTAC set, we observed excellent hit rates (64% of the crude PROTACs show a degradation above 60% at 1 μM and 18% at 200 nM crude PROTAC concentration, respectively). Due to the large number of potent hits, we re-screened the library in HiBiT lytic assays (MV4-11 cells) at a reduced PROTAC concentration (50 nM) to better differentiate moderate from potent degraders (**Figure 6A**). Applying a 60% degradation threshold at this concentration, we still identified 8 potent hits at this low screening concentration (hit rate of 4%). We ranked the PROTACs according to their degradation potency at 50 nM and correlated the HiBiT degradation efficacy values with data measured at 1 μM (**Figure 6B**). The comparison of these data showed excellent overall correlation at both concentrations allowing us to identify most active PROTACs (A16 + X7 and A13 + X30) for this highly degradable POI (**Figure 6C**). The high hit rate allowed us to derive structure-activity relationship trends (**Figure 6D**, at 50 nM): The PEG linkers with an *N*-functionalization at the thalidomide (Thal) resulted in potent POI degradation, followed by alkyl linkers. The use of benzamide (BA)-derived ligands in combination with an alkyl linker was as effective as the combination of thalidomide and PEG linkers. Overall, ether-functionalized (C-O) phenyl dihydrouracil (PDHU) ligands appeared to be less favorable compared to C(sp^2^)-C(sp^3^) exit vectors. BA-derived CRBN ligands demonstrated comparable performance to those derived from thalidomide but it is likely that these PROTACs show a more favorable profile in terms of neo-substrate degradation and chemical stability.^22^ Overall, heterocyclic linker systems were less effective, indicating that it is necessary to test multiple and diverse heterocycles, as these linkers often demonstrate better metabolic stability^26^ and might enhance pharmacokinetic properties.^27^ Our parallel synthetic approach is well suited for this purpose, as a rapid evaluation of the unpurified PROTACs can be used to quickly investigate an alternative linker choice that might be better suited for an in vivo application.

**Figure 6.**
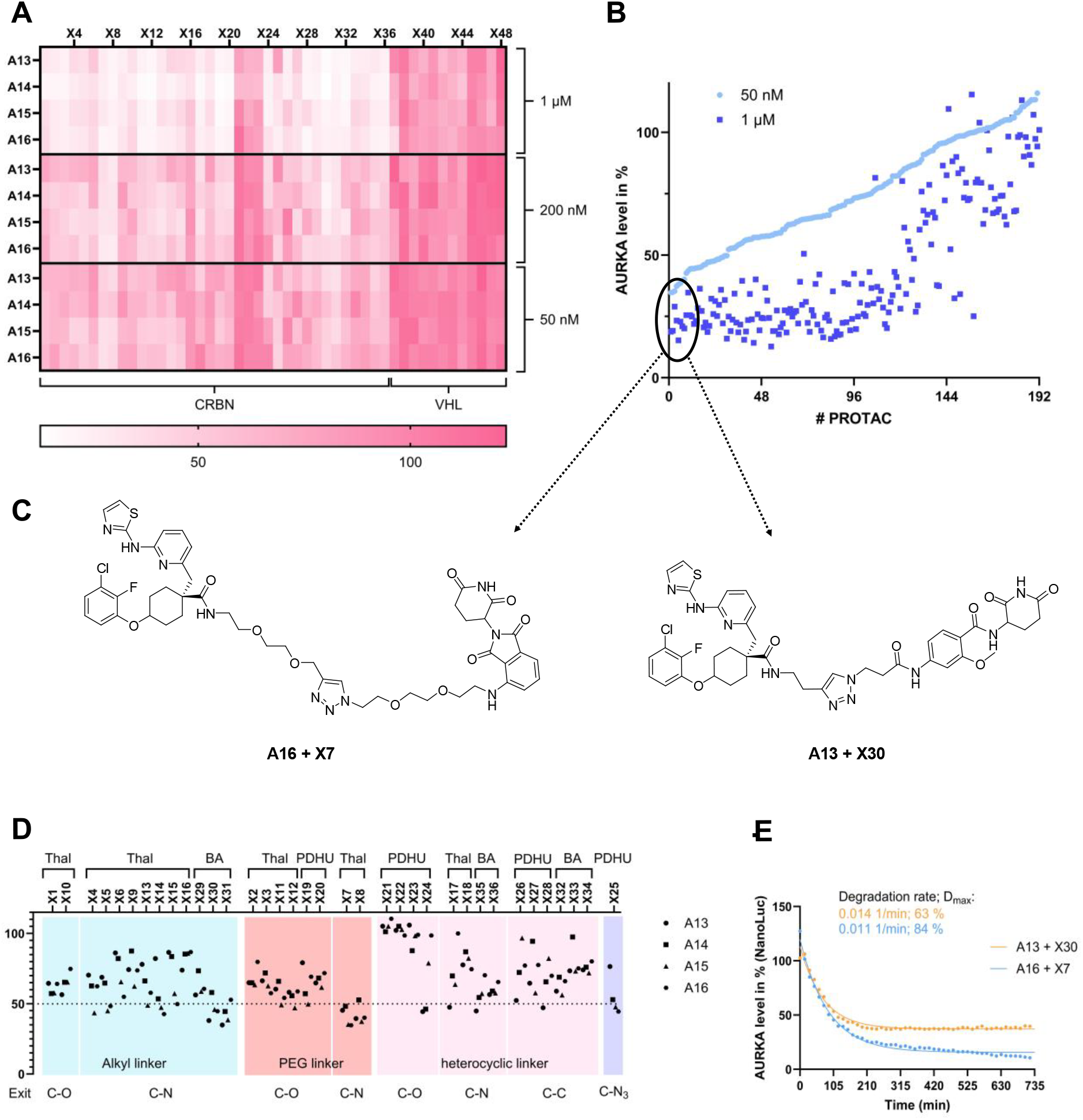
Identification of active PROTACs for AURKA: (A) HiBiT lytic assay-based screening data for AURKA at 50 nM, 200 nM and 1 μM PROTAC concentration after 6 hours incubation time. The clusters of CRBN and VHL containing PROTACs are labeled beneath the heatmap. (B) Screening hits sorted according to degradation efficacy using HiBiT lytic assay data measured at a PROTAC concentration of 50 nM. The data showed good reproducibility of screens carried out at 1 μM and 50 nM PROTAC concentration and identified eight PROTACs with more than 60% POI degradation at 50 nM. (C) Chemical structures of the PROTACs A16 + X7 and A13 + X30 that induce efficient degradation of AURKA (D_max_ > 60%) at a PROTAC concentration of 50 nM. (D) HiBiT-lytic assay-based screening data for AURKA of crude reaction mixtures of CRBN ligand-bearing PROTACs at a PROTAC concentration of 50 nM after 6 h treatment. The data is sorted by different components of the PROTACs: E3 ligand (Thalidomide: Thal, Benzamide: BA, PDHU: phenyl dihydrouracil), linker (alkyl, PEG, heterocyclic) and exit vector composition (C-O, C-N, C-C or C-N3). (E) NanoLuc time dependent live cell measurement monitoring the degradation of AURKA at a PROTAC concentration of 1 μM (crude reaction mixtures of PROTACs A16 + X7 and A13 + X30). The initial phase of each concentration-dependent degradation curve was fitted using a one-component exponential decay model in GraphPad Prism. From this analysis, the best fit parameters K (degradation rate) and plateau (minimum remaining fraction) were determined.

A detailed analysis of degradation kinetics using a NanoLuc live cell system allowed us to gain insight into time dependant effects (**Figure 6E**). We investigated both the degradation rate and the D_max_ within the defined time window. PROTACs A13 + X30 and A16 + X7 both showed a rapid onset of degradation with a degradation rate of 0.014 min^-1^ and 0.011 min^-1^ respectively. Interestingly, A13 + X30 reached a degradation plateau at D_max_ of 63% while A16 + X7 even outperformed the other hits with a D_max_ of 84%.

## DISCUSSION

The empirical nature of TPD drug discovery poses a significant hurdle towards the development of new degrader molecules. We used a D2B approach that allowed us to rapidly interrogate linker SAR, estimate trends in hijackable E3 ligases for degradation and quickly evaluate adequate POI ligand choices. In contrast to classical medicinal chemistry programs, with systematic synthesis and isolation of chemical matter, we were able to address all variables of PROTAC optimization in parallel without the need for time consuming PROTAC purification. This approach positively impacted the overall synthetic output and substantially reduced the time needed from molecule assembly to cellular assay. While it is true that significant synthetic work is needed beforehand for the synthesis of the library azide components and the design of corresponding alkynes, we believe that the overall value of this technology greatly outweighs the *ab initio* effort, especially since the same azide collection can be applied to multiple targets. In fact, sustainability is not only achieved via repurposing of the library - we managed to drastically decrease the scale of the CuAAC reaction with a consumption of a single azide in the two-digit μg range. D2B combined with high throughput screening methods enables the generation of large SAR datasets. Analysis and handling of this data dense output can become a bottleneck and should be combined with computational chemistry and machine learning systems to further streamline PROTAC design and enhance our understanding for adequate POI/E3 combinations, linker choice and overall degradability of a target. The D2B nanoscale synthesis reported in this study has been used to degrade proteins from four different families and we anticipate this methodology to be applied to disease relevant targets in future studies. We offer this method as an enabling technology to the TPD community interested in quick and efficient assessment of the degradability of a target of interest. This is further facilitated by introducing a ready-to-use, pre-plated storage format of the library that allows rapid assembly of degrader molecules. Future efforts will be focused on expanding the library with the aim of introducing more complex and rigid linker systems that have been reported to positively influence physicochemical and pharmacokinetic properties as well as new E3 ligands that may also allow tissue specific degradation based on E3 ligase expression and will increase the diversity of the E3-ligand/linker library.

## Limitations of the study

In this study we developed a technology for the fast and parallel synthesis of PROTACs in a miniaturized format with subsequent cellular evaluation. While the copper catalyzed click reaction is generally very efficient and high yielding, we encountered different degrees of reaction conversion ranging from 70%-95% depending on the scaffold of the POI ligand alkyne. We showed that increasing the amount of both POI ligand alkyne and E3 ligand azide can have a detrimental effect on the degradation efficacy and thus unreacted starting material can lead to a higher occurrence of false negative hits. Longer reaction times or scavenging materials can be employed that could conceivably remove excess starting material and lead to a cleaner conversion profile.

Our findings indicate that technical and reaction specific aspects including sample handling, reactant transfer and starting material conversion are crucial considerations for the overall success of the D2B approach. Indeed, degradation data can be easily confounded by these factors and should be considered a caveat when interpreting negative assay data. A comprehensive FAQ section that offers insights into technical aspects we encountered during the study is provided in the Supplementary Materials (**FAQ S1**).

Apart from copper catalyzed cycloadditions different reactions have been applied to D2B. Indeed, transformations such as the Staudinger ligation, Amide coupling, reductive amination and the inverse electron demand Diels Alder (iEDDA) cycloaddition have been shown to be amenable for plate-based synthesis.^28^ In this study we focused exclusively on the azide alkyne cycloaddition which limits linker modifications to one chemotype. Expanding the reaction toolbox will increase chemical variability and allow a more profound investigation of linkerology.

Lastly, this technology relies on the availability of high throughput, *in cellulo* on target assays with fast read out. Sensor cell lines tagged with e.g. HiBiT, eGFP/mCherry or NanoLuc are widely used to monitor degradation in cells. For novel targets these cell lines need to be generated beforehand, ideally with varying degrees of protein expression. Additionally, the non-availability of suitable control degrader molecules complicates the validation of the new sensor cell lines and thus the applicability to the target of interest. Global chemoproteomic analyses, computational mapping of protein-protein interactions of E3 ligases and POI and artificial proximity assays that allow to investigate suitable POI-E3 ligase pairs for the degradability of a target are pivotal to increasing the success rate of this methodology.

## SIGNIFICANCE

PROTAC development is one of the main scientific challenges in the field of targeted protein degradation. The empirical optimization of numerous parameters associated with PROTAC activity such as, cellular permeability, engagement in ternary complex formation or choice of E3 ligase poses a sincere bottleneck in medicinal chemistry campaigns towards active degrader molecules. We developed a streamlined, direct-to-biology synthesis platform designed to quickly assemble degrader molecules using copper-catalyzed azide-alkyne cycloaddition. We were able to synthesize 192 unique PROTACs per target and screen the molecules on four proteins of interest. This method increases the synthetic output and substantially cuts the time from compound assembly to screening thus enabling quick hit identification and degradability assessment.

## ACKNOWLEDGEMENTS

The authors are grateful for support by the Structural Genomics Consortium (SGC), a registered charity (no. 1097737) that receives funds from Bayer AG, Boehringer Ingelheim, Bristol Myers Squibb, Genentech, Genome Canada through Ontario Genomics Institute [OGI-196], EU/EFPIA/OICR/McGill/KTH/Diamond Innovative Medicines Initiative 2 Joint Undertaking [EUbOPEN grant 875510], Janssen, Merck KGaA (aka EMD in Canada and U.S.), Pfizer, and Takeda. F.A.G., M.P.S., and S.K. would like to acknowledge funding from the German Cancer Consortium (DKTK) at the German Cancer Research Center (DKFZ). M.P.S. is funded by the Deutsche Forschungsgemeinschaft (DFG, German Research Foundation), CRC1430 (Project-ID 424228829). S.K., M.M., K.H., N.L. are grateful for support by the translational cancer program of the German Cancer Aid TACTIC (70115201). Special thanks to Patrick Glenz (P.G.) and Luis Marc Rocktäschel (L.M.R.) for supporting the synthesis of this study during an internship under the supervision of M.M. and T.H.

## AUTHOR CONTRIBUTIONS

Conceptualization: S.K., E.P., F.A.G., T.H., K.H., M.M.; Data curation: M.M., Y.C.G.; Funding acquisition: S.K., E.W.; Investigation: M.M., Y.C.G., A.L., L.H., J.S., R.C., N.L., M.P.S., K.H., M.E., M.L., V.M.; methodology: K.H., M.M.; Project administration: M.M., F.A.G., T.H. K.H.; Writing the original draft: F.A.G, K.H., M.M.; Supervision: S.K., S.M., E.P., T.H., D.M., F.A.G., K.H.; Writing - review & editing: S.K., S.M., E.P., E.W.; Manuscript review: all authors.

## METHODS

- KEY RESOURCES TABLE
- RESOURCE AVAILABILITY
  ○ Lead contact
  ○ Materials availability
  ○ Data and code availability
- EXPERIMENTAL MODEL AND SUBJECT DETAILS
  ○ Cell lines
- METHOD DETAILS
  ○ Setup of the miniaturized CuAAC reaction in 384 well plate format
  ○ Synthetic procedures and characterization
  ○ Cell viability assay
  ○ HiBiT endpoint detection for BRD4 degradation
  ○ HiBiT endpoint detection for WDR5 and AURKA degradation
  ○ Nanoluciferase live cell measurement for WDR5 and AURKA degradation
  ○ HiBiT endpoint detection for sEH degradation
- QUANTIFICATION AND STATISTICAL ANALYSIS

