## Supplementary material for "Click. Screen. Degrade. A Miniaturized D2B Workflow for rapid PROTAC Discovery": Methods

### RESOURCE AVAILABILITY

#### Lead contact

#### Materials availability

The reagents generated in this study will be made available by the lead contact and subject to a completed material transfers agreement.

#### Data and code availability

- All data reported in this paper will be shared by the lead contact upon request.
- This paper does not report original code.
- Any additional information required to reanalyze the data reported in this paper is available from the lead contact upon request

### EXPERIMENTAL MODEL AND STUDY PARTICIPANT DETAILS

#### Cell lines

HEK293 (female, fetus) cells were regularly tested for mycoplasma infection. Cells were grown in DMEM medium supplemented with 10% fetal bovine serum (FBS) and 1% Penicillin/Streptomycin (100 U/ml penicillin and 100 mg/ml streptomycin) at 37 °C and 5% $CO_2$. Human MV4-11 (male, acute monocytic leukemia) cells were regularly tested for mycoplasma infection. Cells were grown in RPMI-1640 medium (Thermo Fisher Scientific) supplemented with 10% FBS and 1% Penicillin/Streptomycin (100 U/ml penicillin and 100 mg/ml streptomycin) at 37 °C in 5% $CO_2$. HeLa cells were stably transfected with sEH-HiBiT fusion protein and regularly tested for mycoplasma infection. Cells were grown in DMEM medium supplemented with 10% FBS, 1% Penicillin/Streptomycin (100 U/ml penicillin and 100 mg/ml streptomycin) and 1 mM sodium pyruvate at 37 °C and 5% $CO_2$.

### METHOD DETAILS

#### Setup of the miniaturized CuAAC reactions in 384 well plate format

Using the example of the synthesis of 192 unique BRD4 PROTACs: 50 mM stock solutions of alkynes **A1-4** and azides **X1-48** in DMSO as well as 60 mM stock solutions of

$CuSO_4 \cdot 5H_2O$ and sodium ascorbate in $H_2O$ were prepared. First, 4 x 2 µL of each azide stock solution were manually pipetted with an E1-ClipTip™ Electronic Multichannel pipette on to the 384 well plate (4 x 2 µL x 48, in total 192 occupied wells with 2 µL of the respective azide stock solution). Next, 2 µL of the respective alkyne (48 wells per alkyne), 0.5 µL of $CuSO_4 \cdot 5H_2O$ (in each well) and 0.5 µL of sodium ascorbate (in each well) stock solution were added to the 384 well plate with the MANTIS® Liquid Dispenser. The plate was sealed and put on a plate shaker (300 rpm) at rt for 24 h (In-depth description of miniaturized plate setup in Supplemental information, Figure S1).

**Synthetic procedures and characterization**

The synthesis of compounds will be explained in the following. All chemicals were purchased from common suppliers with a purity ≥ 95% and were used without further purification. The solvents with an analytical grade were obtained from VWR Chemicals and Merck. Dried solvents were purchased from Acros, stored over molecular sieves and kept under an inert atmosphere. Solvents used in column chromatography or flash column chromatography were technical grade. Perdeuterated solvents were purchased from Eurofins. All microwave-assisted reactions were carried out in sealed reaction vials (0.5 – 10 mL) with a Biotage Initator Microwave System with Robot Eight by Biotage. Flash column chromatography purifications were carried out with a puriFlash XS 520 Plus system from Interchim. For normal-phase chromatography PF-30SIHP-JP-F0024 columns were used with a gradient of DCM and methanol serving as the mobile phase. For reverse-phase chromatography PF-30C18HP-F0012 and PF-30C18HP-F0025 columns were used with a gradient of $H_2O$ and ACN serving as the mobile phase. To monitor the progression of the reactions and determining the purity of compounds analytical high-performance liquid chromatography (HPLC) was performed. A 1260 Infinity II LC System consisting of the multisampler G7167A, the column compartment G7116A, the multicolumn thermostat G7116A, the flexible pump G7104C, a single quadrupole LC/MSD system InfinityLab G6125B and the diode array detector HS G7117C by the company Agilent Technologies was used for this purpose. An ACE UltraCore Super C18 column (150 x 3.0 mm) from Avantor was used as the stationary phase and a gradient of $H_2O$ and ACN with 0.1% formic acid served as the mobile phase. UV detection took place at wavelengths of 254 and 280 nm. The following gradient was used: 0 min: 5% B -

2 min: 80% B – 5 min: 95% B - 7 min: 95% B (flow rate of 0.6 mL/min). Nuclear magnetic resonance (NMR) spectra were recorded with spectrometers DPX250 (250 MHz $^{1}H$), AV300 (300 MHz $^{1}H$, 282 MHz $^{19}F$), AV400 (400 MHz $^{1}H$, 101 MHz $^{13}C$, 377 MHz $^{19}F$) and AV500 (500 MHz $^{1}H$, 126 MHz $^{13}C$) from Bruker with all the measurements being performed at rt and in deuterated solvents. Chemical shifts (δ) are reported in parts per million (ppm) and refer to the internal standard tetramethylsilane at 0.00 ppm and to the residual solvent signal. DMSO-$d_6$, acetone-$d_6$, methylene chloride-$d_2$ and chloroform-*d* were used as a solvent, and the spectra were calibrated to the solvent signal: 2.50 ppm ($^{1}H$ NMR) or 39.52 ppm ($^{13}C$ NMR) for DMSO-$d_6$, 2.05 ppm ($^{1}H$ NMR) or 206.26 ppm ($^{13}C$ NMR) for acetone-$d_6$, 5.32 ppm ($^{1}H$ NMR) or 53.84 ppm ($^{13}C$ NMR) for methylene chloride-$d_2$ and 7.26 ppm ($^{1}H$ NMR) or 77.16 ppm ($^{13}C$ NMR) for chloroform-*d*. The coupling constant $J$ was stated in Hz. The multiplicity M of the signals in the spectra was characterized by following abbreviations: s (singlet), d (doublet), dd (doublet of doublets), t (triplet), td (triplet of doublets), quartet (q), quintet (quin), septet (sept), m (multiplet).

**General Procedure A for amide coupling**

Carboxylic acid (1.0 eq) and HATU (1.2 – 1.3 eq) were dissolved in DMF. Amine (1.0 eq) and DIPEA (3.0 eq) were added to the resulting mixture and it was stirred at rt for 2 h. The solvent was removed under reduced pressure and the crude product was purified by flash chromatography using acetonitrile/water as an eluent.

**General Procedure B for amide coupling**

Carboxylic acid (1.0 eq), 1-methyl-1*H*-imidazole (3.5 eq), amine (1.0 – 1.5 eq.) and *N,N,N',N'*-tetramethylchloroformamidinium hexafluorophosphate (TCFH) (1.2 eq) were dissolved in dry acetonitrile. The reaction mixture was stirred at rt for 18 h. The reaction mixture was diluted with EtOAc and washed with saturated $NaHCO_3$ (3x) and brine (1x). The organic phase was dried over $MgSO_4$ and the solvent was removed under reduced pressure. The crude product was purified by flash chromatography using acetonitrile/water as an eluent.

**General Procedure C for microwave-assisted nucleophilic aromatic substitution**

Fluoro derivative (1.0 eq), amine (1.2 – 1.5 eq) and DIPEA (3.0 – 4.0 eq) were charged in a microwave vial and dissolved in DMSO. The suspension was degassed with argon under sonication. The vial was heated in a microwave oven to 150 °C for five minutes. The solvent was removed under reduced pressure and the crude product was purified by flash chromatography using acetonitrile/water as an eluent.

**General Procedure D for *N*-Boc deprotection**

*N*-Boc protected amines were dissolved in dry DCM. The flask was cooled to 0 °C in an ice bath and trifluoroacetic acid (TFA) in an excess was added dropwise. The solution was stirred at rt for 2 h. The solvent was removed under reduced pressure and the crude product used without further purification.

**General Procedure E for copper-catalyzed azide-alkyne cycloaddition**

Azide (1.0 eq) and Alkyne (1.0 eq) were dissolved in DMSO. Copper sulfate pentahydrate (0.3 eq) and sodium ascorbate (0.3 eq) were dissolved in water and added to the reaction mixture. The solution was stirred at rt for 18 h. The solvent was removed under reduced pressure and the crude product was purified by flash chromatography using acetonitrile/water as an eluent.

**Synthesis of (*S*)-2-(4-(4-chlorophenyl)-2,3,9-trimethyl-6*H*-thieno[3,2-*f*][1,2,4]triazolo[4,3-*a*][1,4]diazepin-6-yl)-*N*-(prop-2-yn-1-yl)acetamide (A1)**

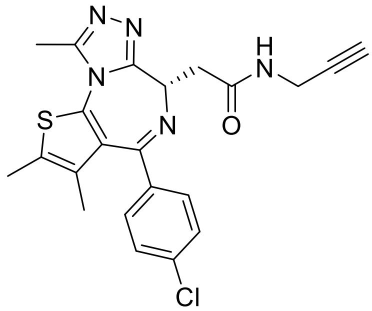


The title compound was prepared according to general procedure A, using (*S*)-2-(4-(4-chlorophenyl)-2,3,9-trimethyl-6*H*-thieno[3,2-*f*][1,2,4]triazolo[4,3-*a*][1,4]diazepin-6-yl)acetic acid (150 mg, 0.374 mmol), prop-2-yn-1-amine (20 mg, 0.374 mmol.), HATU (184 mg, 0.486 mmol) and DIPEA (195 µL, 1.12 mmol). The title compound was obtained as a colourless solid (152 mg, 93%). $^1$H NMR (400 MHz, DMSO-$d_6$): δ 8.67 (t, *J* = 5.6 Hz, 1H), 7.46 (dd, 4H), 4.50 (dd, *J* = 8.6, 5.7 Hz), 4.03 – 3.95 (m1H), 3.91 – 3.82 (m, 1H), 3.21 – 3.14 (m, 2H), 2.60 (s, 3H), 2.41 (s, 3H), 1.62 (s, 3H). $^{13}$C NMR (101 MHz, DMSO-$d_6$): δ 169.74, 163.04, 155.10, 149.83, 136.75, 135.24, 132.28, 130.70, 130.18, 129.84, 129.56, 128.46, 82.35, 72.05, 53.81, 37.91, 37.55, 18.82, 14.07, 12.68, 11.30. ESI-MS: *m/z* = 438.10 ([M+H]$^+$).

**Synthesis of (*S*)-*N*-(but-3-yn-1-yl)-2-(4-(4-chlorophenyl)-2,3,9-trimethyl-6*H*-thieno[3,2-*f*][1,2,4]triazolo[4,3-*a*][1,4]diazepin-6-yl)acetamide (A2)**

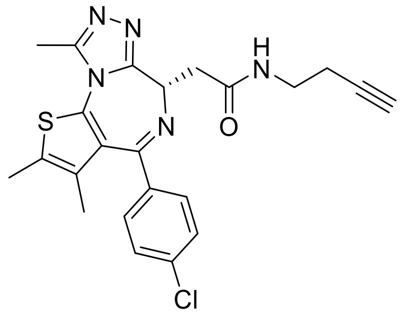


The title compound was prepared according to general procedure A, using (*S*)-2-(4-(4-chlorophenyl)-2,3,9-trimethyl-6*H*-thieno[3,2-*f*][1,2,4]triazolo[4,3-*a*][1,4]diazepin-6-yl)acetic acid (100 mg, 0.249 mmol), but-3-yn-1-amine (26 mg, 0.25 mmol.), HATU (123 mg, 0.324 mmol) and DIPEA (130 µL, 0.748 mmol). The title compound was obtained as a colourless solid (106 mg, 95%). $^1$H NMR (400 MHz, DMSO-$d_6$): δ 8.41 (t, *J* = 5.7 Hz), 7.46 (dd, 4H), 4.51 (dd, *J* = 8.4, 5.7 Hz, 1H), 3.33 – 3.13 (m, 4H), 2.86 (t, *J* = 2.7 Hz, 1H), 2.59 (s, 3H), 2.54 (s, 1H), 2.50 (s, 1H), 2.41 (s, 3H), 2.37 – 2.32 (m, 2H), 1.62 (s, 3H). $^{13}$C NMR (101 MHz, DMSO-$d_6$): δ 169.74, 163.04, 155.10, 149.83, 136.75, 135.24, 132.28, 130.70, 130.18, 129.84, 129.56, 128.46, 82.35, 72.05, 53.81, 37.91, 37.55, 18.82, 14.07, 12.68, 11.30. ESI-MS: *m/z* = 452.10 ([M+H]$^+$).

**Synthesis of (*S*)-2-(4-(4-chlorophenyl)-2,3,9-trimethyl-6*H*-thieno[3,2-*f*][1,2,4]triazolo[4,3-*a*][1,4]diazepin-6-yl)-*N*-(pent-4-yn-1-yl)acetamide (A3)**

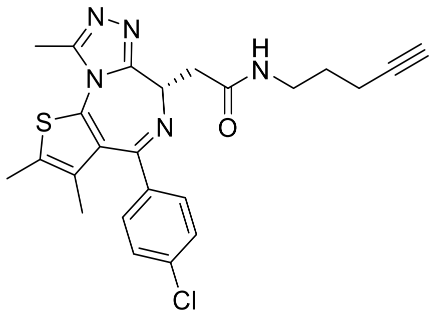

The title compound was prepared according to general procedure A, using (*S*)-2-(4-(4-chlorophenyl)-2,3,9-trimethyl-6*H*-thieno[3,2-*f*][1,2,4]triazolo[4,3-*a*][1,4]diazepin-6-yl)acetic acid (100 mg, 0.249 mmol), pent-4-yn-1-amine (30 mg, 0.25 mmol.), HATU (123 mg, 0.324 mmol) and DIPEA (130 µL, 0.748 mmol). The title compound was obtained as a colourless solid (112 mg, 97%). $^{1}$H NMR (400 MHz, DMSO-$d_6$): δ 8.23 (t, *J* = 5.6 Hz, 1H), 7.46 (dd, 4H), 4.51 (dd, *J* = 8.2, 6.0 Hz, 1H), 3.26 – 3.08 (m, 3H), 2.80 (t, *J* = 2.7 Hz, 1H), 2.59 (s, 3H), 2.41 (s, 3H), 2.21 (td, *J* = 6.9, 3.9 Hz, 2H), 1.63 (d, *J* = 4.9 Hz, 5H). $^{13}$C NMR (101 MHz, DMSO-$d_6$): δ 169.56, 163.08, 155.11, 149.82, 136.80, 135.23, 132.28, 130.70, 130.12, 129.84, 129.58, 128.47, 84.10, 71.32, 53.89, 37.64, 28.27, 15.39, 14.04, 12.67, 11.30. ESI-MS: *m/z* = 466.15 ([M+H]$^{+}$).

**Synthesis of (*S*)-2-(4-(4-chlorophenyl)-2,3,9-trimethyl-6*H*-thieno[3,2-*f*][1,2,4]triazolo[4,3-*a*][1,4]diazepin-6-yl)-1-(4-(prop-2-yn-1-yl)piperazin-1-yl)ethan-1-one (A4)**

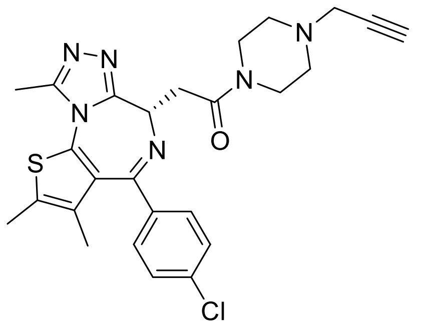

The title compound was prepared according to general procedure A, using (*S*)-2-(4-(4-chlorophenyl)-2,3,9-trimethyl-6*H*-thieno[3,2-*f*][1,2,4]triazolo[4,3-*a*][1,4]diazepin-6-yl)acetic acid (100 mg, 0.249 mmol), 1-(prop-2-yn-1-yl)piperazine (31 mg, 0.25 mmol.), HATU (123 mg, 0.324 mmol) and DIPEA (130 µL, 0.748 mmol). The title compound was obtained as a colourless solid (115 mg, 91%). $^1$H NMR (400 MHz, DMSO-$d_6$): δ 7.56 – 7.38 (m, 4H), 4.58 (t, *J* = 6.7 Hz, 1H), 3.67 (q, *J* = 4.4 Hz, 2H), 3.61 (dd, *J* = 16.4, 7.1 Hz, 1H), 3.50 (dt, *J* = 21.6, 5.3 Hz, 2H), 3.41 (dd, *J* = 16.4, 6.3 Hz, 1H), 3.19 (t, 1H), 2.60 (s, 3H), 2.56 – 2.52 (m, 2H), 2.41 (s, 4H), 2.07 (s, 3H), 1.63 (s, 3H). $^{13}$C NMR (101 MHz, DMSO-$d_6$): δ 168.11, 162.85, 155.27, 149.75, 136.78, 135.19, 132.20, 130.66, 130.15, 129.89, 129.64, 128.47, 79.06, 75.96, 54.14, 51.45, 50.90, 45.97, 44.79, 40.98, 40.43, 39.52, 34.76, 14.01, 12.68, 11.27. ESI-MS: *m/z* = 507.10 ([M+H]$^+$).

**Synthesis of methyl 1-(3-fluorobenzyl)-*1H*-indole-5-carboxylate (I1)**

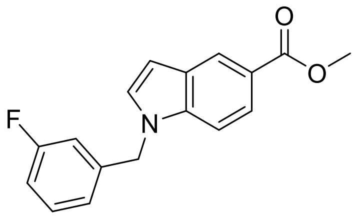


To a solution of methyl indole-5-carboxylate (2.00 g, 11.2 mmol, 1.0 eq) and 3-fluorobenzyl chloride (1.78 g, 12.3 mmol, 1.1 eq) in dry DMF (50 mL) were added NaH (60 % dispersion in mineral oil, 0.67 g, 16.8 mmol, 1.5 eq) and KI (catalytic amount). The mixture was sparged with argon and stirred at rt. for 3 h. Afterwards, the reaction was quenched with water (100 mL) and extracted with EtOAc (3 x 100 mL). The combined organic phase was dried over $MgSO_4$ and the solvent was removed under reduced pressure. The crude product was purified by flash chromatography (*n*-Hex: EtOAc = 90:10 → 20:80) to afford the title compound as a pale-yellow solid in 57% yield (1.82 g, 6.41 mmol). $^1$H NMR (300 MHz, DMSO-$d_6$): δ 8.28 (s, 1H), 7.74 (dd, *J* = 8.7, 1.5 Hz, 1H), 7.65 (d, *J* = 3.2 Hz, 1H), 7.57 (d, *J* = 8.7 Hz, 1H), 7.42 – 7.29 (m, 1H), 7.15 – 6.97 (m, 3H), 6.68 (d, *J* = 3.2 Hz, 1H), 5.50 (s, 2H), 3.83 (s, 3H). ESI-MS: *m/z* = 284.05 ([M+H]$^+$).

**Synthesis of 1-(3-fluorobenzyl)-*1H*-indole-5-carboxylic acid (I2)**

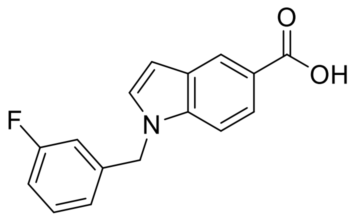


To a solution **I1** (656 mg, 2.32 mmol, 1.0 eq) in THF (12.5 mL) and MeOH (12.5 mL) was added dropwise a 0.5 M aqueous solution of KOH (11.60 mol, 5.0 eq). The mixture was stirred at 70 °C for 16 h. Afterwards, the reaction mixture was cooled to rt and the solvent mixture was evaporated. The residue was then acidified with 2 M aqueous HCl (pH = 3) and the resulting aqueous solution was extracted with DCM (3 x 20 mL). The combined organic phase was dried over $MgSO_4$ and the solvent was removed under reduced pressure to afford the title compound as a pale-yellow solid in 98% yield (610 mg, 2.27 mmol). $^1$H NMR (250 MHz, DMSO-$d_6$): δ 12.45 (bs, 1H), 8.25 (d, *J* = 1.1 Hz, 1H), 7.72 (dd, *J* = 8.7 Hz, *J* = 1.6 Hz, 1H), 7.63 (d, *J* = 3.2 Hz, 1H), 7.54 (d, *J* = 8.7 Hz, 1H), 7.43 – 7.29 (m, 1H), 7.13 – 6.98 (m, 3H), 6.66 (dd, *J* = 3.2 Hz, *J* = 0.6 Hz, 1H), 5.49 (s, 2H). ESI-MS: *m/z* = 268.10 ($[M+H]^+$).

**Synthesis of *N*-(4-amino-2-(trifluoromethyl)benzyl)-1-(3-fluorobenzyl)-1*H*-indole-5-carboxamide (I3)**

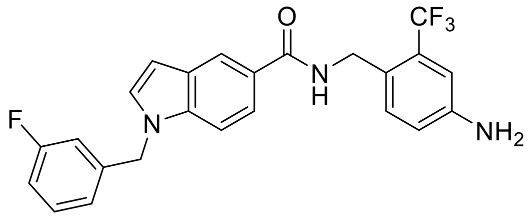


To a solution of **I2** (100 mg, 0.35 mmol, 1.0 eq) in dry THF (9 mL) were added DIPEA (0.2 mL, 1.06 mmol, 3.0 eq) and PyBOP (202 mg, 0.39 mmol, 1.1 eq). The mixture was stirred at rt for 10 min before 4-amino-3-(trifluoromethyl)benzylamine (69 mg, 0.35 mmol, 1.0 eq) and HOBt·$H_2O$ (27 mg, 0.18 mmol, 0.5 eq) were added. The resulting reaction mixture was stirred at rt for 16 h. Afterwards, the mixture was diluted with EtOAc (10 mL) and washed with brine (15 mL). The aqueous phase was extracted with EtOAc (3 x 15 mL) and the combined organic phase was dried over $MgSO_4$. After removing the solvent under reduced pressure, the crude product was purified by flash chromatography (*n*-Hex: EtOAc = 2:1 → 1:1) to afford the title compound as a yellow solid in 89% yield (138 mg, 0.31 mmol). $^1$H NMR (250 MHz, DMSO-$d_6$): δ 8.70 (t, *J* = 5.7 Hz, 1H), 8.20 (d, *J* = 1.2 Hz, 1H), 7.69 (dd, *J* = 8.7 Hz, *J* = 1.6 Hz, 1H), 7.61 (d, *J* = 3.2 Hz, 1H), 7.52 (d, *J* = 8.7 Hz, 1H), 7.41 – 7.28 (m, 1H), 7.16 (d, *J* = 8.4 Hz, 1H), 7.13 – 6.97 (m, 3H), 6.90 (d, *J* = 2.3 Hz, 1H), 6.74 (dd, *J* = 8.3 Hz, *J* = 2.1 Hz, 1H), 6.62 (d, *J* = 3.1 Hz, 1H), 5.49 (s, 2H), 5.43 (s, 2H), 4.49 (d, *J* = 5.4 Hz, 2H,). ESI-MS: *m/z* = 441.90 ([M+H]$^+$).

**Synthesis of 1-(3-fluorobenzyl)-*N*-(4-(pent-4-yn-1-ylsulfonamido)-2-(trifluoromethyl)-benzyl)-*1H*-indole-5-carboxamide (A5)**

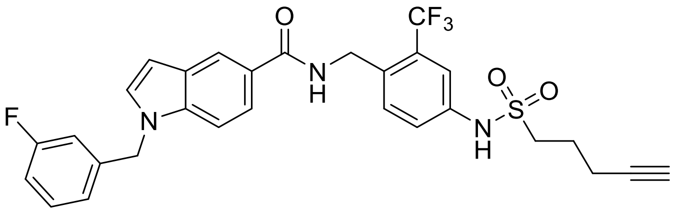


**I3** (100 mg, 0.25 mmol, 1.0 eq) was dissolved in 10 mL dry $CHCl_3$ and the solution was sparged with argon. Then, 1-sulfonylchloride-4-pentyne (52 mg, 0.29 mmol, 1.2 eq) and pyridine (0.1 mL, 1.09 mmol, 5.0 eq) were added. The reaction mixture was stirred at 55 °C for 48 h. After cooling to rt the solution was acidified with 2 M aqueous HCl (pH = 3) and extracted with DCM (3 x 15 mL). The combined organic phase was dried over $MgSO_4$ and the solvent was removed under reduced pressure. Purification using reversed phase flash chromatography (ACN/$H_2O$ = 30:70 → 10:90) afforded the title compound as a colourless solid in 74% yield (96 mg, 0.18 mmol). $^1$H NMR (400 MHz, DMSO-$d_6$): δ 10.16 (s, 1H, N*H*), 8.92 (t, $^3J_{HH}$ = 5.8 Hz, 1H, N*H*), 8.22 (d, *J* = 1.3 Hz, 1H), 7.70 (dd, *J* = 8.7 Hz, *J* = 1.6 Hz, 1H), 7.63 (d, *J* = 3.2 Hz, 1H), 7.54 (d, *J* = 8.9 Hz, 2H), 7.52 – 7.42 (m, 2H), 7.35 (m, 1H), 7.12 – 7.05 (m, 1H), 7.04 – 6.99 (m, 2H), 6.64 (dd, *J* = 3.2 Hz, *J* = 0.6 Hz, 1H*),* 5.50 (s, 2H), 4.61 (d, *J* = 5.4 Hz, 2H), 3.23 – 3.17 (m, 2H), 2.76 (t, *J* = 2.6 Hz, 1H), 2.28 (td, *J* = 7.0 Hz, *J* = 2.6 Hz, 2H), 1.82 (m, 2H). $^{13}$C NMR (126 MHz, DMSO-$d_6$): δ 167.4, 158.0, 137.3, 137.2, 130.73, 130.7, 130.61, 130.57 127.7, 125.5, 122.9, 120.9, 120.6, 120.2, 114.3, 113.94, 113.88, 113.6, 109.8, 102.4, 82.9, 74.9, 72.22, 72.18, 72.1, 49.9, 48.6, 22.4, 16.2. $^{19}$F NMR (282 MHz, DMSO-$d_6$): δ -59,7 (s), -112,1 (s). ESI-MS: *m/z* = 572.23 ([M+H]$^+$).

**Synthesis of *tert*-Butyl-((1*S*,3*R*)-3-((4-amino-2-(trifluoromethyl)benzyl)carbamoyl)-cyclohexyl)carbamate (I4)**

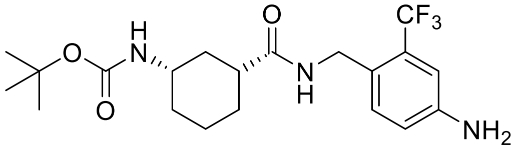

4-(Aminomethyl)-3-trifluoromethyl)aniline (200 mg, 0.78 mmol, 1.0 eq) and PyBOP (447 mg, 0.86 mmol, 1.1 eq) were dissolved in dry THF (15 mL). DIPEA (0.41 mL, 2.34 mmol, 3.0 eq), HOBt·$H_2O$ (60 mg, 0.39 mmol, 0.5 eq) and (1*R*,3*S*)-3-((*tert*-Butoxycarbonyl)amino)cyclohexane-1-carboxylic acid (150 mg, 0.78 mmol, 1.0 eq) were added and the reaction mixture was stirred at rt for 16 h. Afterwards, the mixture was diluted with EtOAc (15 mL) and washed with $H_2O$ (2 x 15 mL) and brine (15 mL). The aqueous phase was extracted with EtOAc (3 x 15 mL) and the combined organic phase was dried over $MgSO_4$. After removing the solvent under reduced pressure, the crude product was purified by flash chromatography (*n*-Hex: EtOAc = 100:0 → 70:30) to afford the title compound as a yellow solid in 82% yield (266 mg, 0.64 mmol). $^1$H NMR (250 MHz, $CDCl_3$): δ 7.27 (d, *J* = 8.2 Hz, 1H), 6.91 (d, *J* = 2.5 Hz, 1H), 6.76 (dd, *J* = 2.4 Hz, *J* = 8.2 Hz, 1H), 5.72 (t, *J* = 5.4 Hz, 1H), 4.45 (s, 2H), 4.43 (s, 1H), 3.85 (s, 2H), 3.48 – 3.38 (m, 2H), 2.13 – 2.07 (m, 2H), 1.95 – 1.90 (m, 2H), 1.86 – 1.76 (m, 1H), 1.82 – 1.76 (m, 2H), 1.42 (s, 9H), 1.08 – 1.02 (m, 1H). ESI-MS: *m/z* = 416.15 ($[M+H]^+$).

**Synthesis of *tert*-Butyl((1*S*,3*R*)-3-((4-(pent-4-yn-1-ylsulfonamido)-2-(trifluoromethyl)-benzyl)carbamoyl)cyclohexyl)carbamate (I5)**

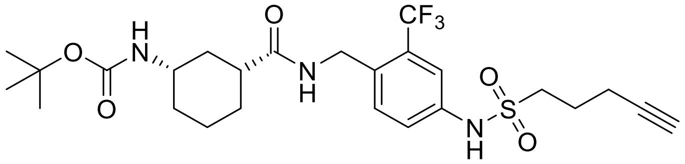

**I4** (100 mg, 0.24 mmol, 1.0 eq), Pent-4-yne-1-sulfonyl chloride (48 mg, 0.27 mmol, 1.1 eq) and Pyridine (0.1 mL, 1.2 mmol, 5.0eq) were dissolved in dry $CHCl_3$ (20 mL) and the reaction mixture was stirred at 60 °C for 48 h. The reaction mixture was diluted with EtOAc (15 mL) and washed with 1 M HCl (2 x 20 mL) and brine (20 mL). The aqueous phase was extracted with EtOAc (3 x 20 mL) and the combined organic phase was dried over $MgSO_4$. After removing the solvent under reduced pressure, the crude product was purified by flash chromatography (*n*-Hex: EtOAc = 100:0 → 80:20) to afford the title compound as a brown oil in 37% yield (99 mg, 0.18 mmol). $^1$H NMR (400 MHz, $CDCl_3$): δ 8.36 (s, 1H), 7.50 (d, *J* = 1.7 Hz, 1H), 7.36 – 7.34 (m, 1H), 7.31 – 7.29 (m, 1H), 6.32 -6.28 (m, 1H), 4.56 (dd, *J* = 7.1 Hz, *J* = 17.6 Hz, 2H), 4.46 (dd, *J* = 5.4 Hz, *J* = 15.7 Hz, 1H), 3.47 (s, 2H), 3.24 – 3.20 (m, 2H), 2.34 (dt, *J* = 2.6 Hz, *J* = 6.7 Hz, 2H), 2.29 – 2.23 (m, 1H), 2.14 – 2.11 (m, 1H), 2.02 (q, *J* = 7.2 Hz, 2H), 1.95 (t, *J* = 2.6 Hz, 1H), 1.90 – 1.86 (m, 1H), 1.86 – 1.78 (m, 2H), 1.43 (s, 9H), 1.31 – 1.29 (m, 2H), 1.14 – 1.04 (m, 1H). ESI-MS: m/z = 544.05 ($[M+H]^+$).

**Synthesis of (1*R*,3*S*)-3-Amino-*N*-(4-(pent-4-in-1-ylsulfonamido)-2-(trifluoromethyl)-benzyl)cyclohexane-1-carboxamide (I6)**

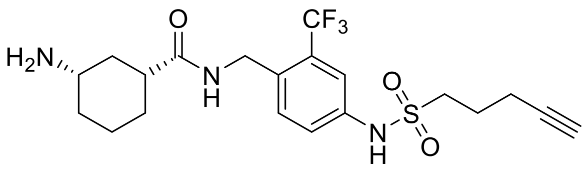


**I5** (83 mg, 0.15 mmol) was dissolved in dry DCM (2.5 mL). The flask was cooled to 0 °C in an ice bath and trifluoroacetic acid (TFA) in an excess was added dropwise. The solution was stirred at rt for two h. The reaction mixture was diluted with saturated aqueous $NaHCO_3$ (pH = 8) and the aqueous phase was extracted with EtOAc (5 x 10 mL). The combined organic phase was dried over $MgSO_4$. All volatiles were removed under reduced pressure to afford the title compound as a colourless solid in 99% yield (68 mg, 0.15 mmol). $^1$H NMR (400 MHz, MeOH-*d*$_4$): δ 7.56 – 7.55 (m, 1H), 7.47 – 7.42 (m, 2H), 4.91 (s,2H), 3.24 – 3.20 (m, 2H), 3.15 (tt, *J* = 11.7 Hz, *J* = 3.92 Hz, 1H), 2.45 (tt, *J* = 11.8 Hz, *J* = 3.5 Hz, 1H), 2.31 (td, *J* = 6.9 Hz, *J* = 2.3 Hz, 2H), 2.21 (t, *J* = 2.6 Hz, 1H), 2.12 – 2.01 (m, 2H), 1.98 – 1.90 (m, 2H), 1.94 – 1.86 (m, 2H), 1.47 – 1.27 (m, 4H). ESI-MS: *m/z* = 446.20 ($[M+H]^+$).

**Synthesis of (1*R*,3*S*)-3-((4-Methyl-6-(methylamino)-1,3,5-triazin-2-yl)amino)-*N*-(4-(pent-4-yn-1-ylsulfonamido)-2-(trifluoromethyl)benzyl)cyclohexane-1-carboxamide (A6)**

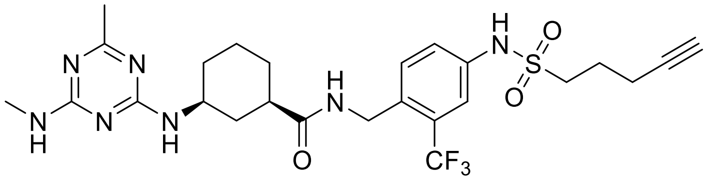

2,4-Dichloro-6-methyl-1,3,5-triazine (26 mg, 0.15 mmol, 1 eq) and methylamine (17 µL, 0.15 mmol, 1 eq) were combined in a 5 mL round bottom flask and mixed with 1M sodium hydroxide solution until a pH of 12 was reached. A solution of **I6** (68 mg, 0.15 mmol, 1 eq) in 1 mL of methanol was added and the pH was adjusted to 10 with 1 M sodium hydroxide solution. The reaction mixture was stirred at 90 °C for 16 h. The solvent was removed under reduced pressure and the crude product was purified by flash chromatography (DCM:MeOH = 100:0 → 90:10) to afford the title compound as a yellow solid in 20% yield (15 mg, 13 µmol). $^1$H NMR (400 MHz, $CD_2Cl_2$): δ 7.56 (d, *J* = 2.0 Hz, 1H), 7.39 (d, *J* = 8.4 Hz, 1H), 7.33 – 7.30 (m, 1H), 4.58 – 4.47 (m, 2H), 3.99 – 3.97 (m, 1H), 3.27 – 3.20 (m, 2H), 2.90 – 2.86 (m, 2H), 2.33 (dt, *J* = 2.6 Hz, *J* = 6.8 Hz, 3H), 2.17 – 3.13 (m, 2H), 2.04 – 1.97 (m, 4H), 1.86 – 1.80 (m, 2H), 1.40 – 1.39 (m, 2H), 1.26 (s, 3H), 1.63 (d, *J* = 6.1 Hz, 1H), 0.89 – 0.84 (m, 1H). ESI-MS: *m/z* = 568.35 ([M+H]$^+$).

**Synthesis of *tert*-butyl 2-((2-(2,6-dioxopiperidin-3-yl)-1,3-dioxoisoindolin-4-yl)oxy)acetate (A7)**

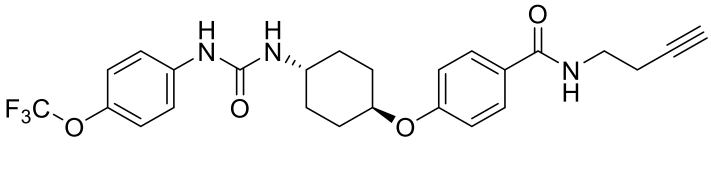

The title compound was prepared according to general procedure A, using 4-(((1*r*,4*r*)-4-(3-(4-(trifluoromethoxy)phenyl)ureido)cyclohexyl)oxy)benzoic acid (20 mg, 0.046 mmol), pent-4-yn-1-amine (5.5 mg, 0.046 mmol.), HATU (23 mg, 0.059 mmol) and DIPEA (24 µL,

0.14 mmol). The title compound was obtained as a colourless solid (13.4 mg, 54%). $^1$H NMR (300 MHz, DMSO-$d_6$) δ 8.51 (s, 1H), 8.30 (t, *J* = 5.5 Hz, 1H), 7.82 – 7.76 (m, 2H), 7.49 – 7.45 (m, 2H), 7.24 – 7.19 (m, 2H), 7.01 – 6.96 (m, 2H), 6.19 (d, *J* = 7.4 Hz, 1H), 4.46 – 4.40 (m, 1H), 3.58 – 3.48 (m, 1H), 3.30 – 3.25 (m, 2H), 2.78 (t, *J* = 2.6 Hz, 1H), 2.24 – 2.17 (m, 2H), 2.07 – 1.92 (m, 4H), 1.74 – 1.64 (m, 2H), 1.55 – 1.31 (m, 4H). $^{19}$F NMR (282 MHz, DMSO-$d_6$): δ – 57.1 (s). ESI-MS: *m/z* = 504.25 ([M+H]$^+$).

**Synthesis of *tert*-butyl 2-((2-(2,6-dioxopiperidin-3-yl)-1,3-dioxoisoindolin-4-yl)oxy)acetate (A8)**

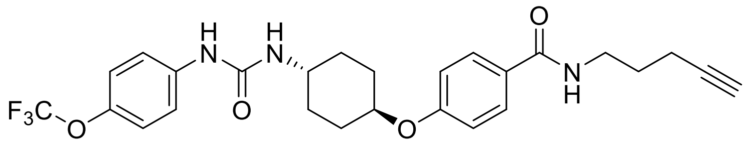


The title compound was prepared according to general procedure A, using 4-(((1*r*,4*r*)-4-(3-(4-(trifluoromethoxy)phenyl)ureido)cyclohexyl)oxy)benzoic acid (20 mg, 0.046 mmol), 3-butyn-1-amine hydrochloride (4.8 mg, 0.046 mmol.), HATU (23 mg, 0.059 mmol) and DIPEA (24 µL, 0.14 mmol). The title compound was obtained as a colourless solid (11.3 mg, 49%). $^1$H NMR (300 MHz, DMSO-$d_6$) δ 8.52 (s, 1H), 8.46 (t, *J* = 5.5 Hz, 1H), 7.82 – 7.77 (m, 2H), 7.50 – 7.45 (m, 2H), 7.25 – 7.19 (m, 2H), 7.04 – 6.99 (m, 2H), 6.20 (d, *J* = 7.6 Hz, 1H), 4.47 – 4.41 (m, 1H), 3.55 – 3.51 (m, 1H), 3.39 – 3.37 (m, 2H), 2.82 (t, *J* = 2.5 Hz, 1H), 2.41 (dt, *J* = 2.4 Hz, 7.1 Hz, 2H), 2.08 – 1.92 (m, 4H), 1.55 – 1.31 (m, 4H). $^{19}$F NMR (282 MHz, DMSO-$d_6$): δ – 57.1 (s). ESI-MS: *m/z* = 490.15 ([M+H]$^+$).

**Synthesis of *N*-(2'-fluoro-5'-(prop-2-yn-1-ylcarbamoyl)-4-((3*S*,5*R*)-3,4,5-trimethylpiperazin-1-yl)-[1,1'-biphenyl]-3-yl)-6-oxo-4-(trifluoromethyl)-1,6-dihydropyridine-3-carboxamide (A9)**

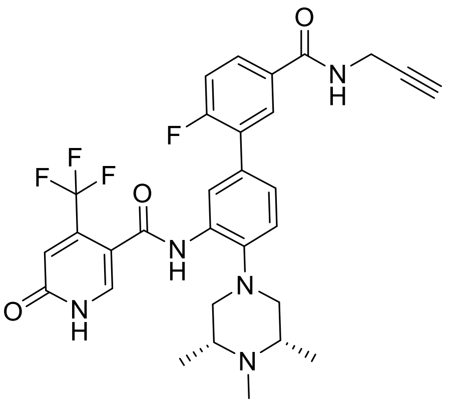

The title compound was prepared according to general procedure A, using 6-fluoro-3'-(6-oxo-4-(trifluoromethyl)-1,6-dihydropyridine-3-carboxamido)-4'-((3*S*,5*R*)-3,4,5-trimethylpiperazin-1-yl)-[1,1'-biphenyl]-3-carboxylic acid (10 mg, 18 µmol), prop-2-yn-1-amine (1.0 mg, 18 µmol), HATU (9.1 mg, 24 µmol) and DIPEA (2.2 µL, 55 µmol). The title compound was obtained as a colourless solid (10.4 mg, 97%). $^{1}$H NMR (400 MHz, DMSO-$d_6$): δ 12.58 (s, 1H), 9.55 (s, 1H), 9.06 (t, *J* = 5.5 Hz, 1H), 8.16 (s, 1H), 8.02 – 7.97 (m, 2H), 7.94 – 7.87 (m, 1H), 7.50 – 7.39 (m, 2H), 7.33 (d, *J* = 8.3 Hz, 1H), 6.83 (s, 1H), 4.07 (dd, *J* = 5.5, 2.5 Hz, 2H), 3.46 (d, *J* = 14.5 Hz, 2H), 3.36 (s, 4H), 3.14 (t, *J* = 2.5 Hz, 1H), 2.89 – 2.82 (m, 3H), 1.33 (d, *J* = 6.3 Hz, 6H). $^{13}$C NMR (101 MHz, DMSO-$d_6$): δ 164.83, 162.94, 161.77, 161.11, 159.77, 142.20, 138.54, 132.29, 130.63, 129.99, 128.78, 127.71, 125.87, 123.97, 123.14, 120.95, 120.64, 116.46, 116.27, 111.53, 89.45, 81.16, 73.02, 59.82, 55.59, 48.59, 36.48, 28.97, 28.58, 14.58. ESI-MS: *m/z* = 584.20 ([M+H]$^{+}$).

**Synthesis of *N*-(2'-fluoro-5'-((2-(prop-2-yn-1-yloxy)ethyl)carbamoyl)-4-((3*S*,5*R*)-3,4,5-trimethylpiperazin-1-yl)-[1,1'-biphenyl]-3-yl)-6-oxo-4-(trifluoromethyl)-1,6-dihydropyridine-3-carboxamide (A10)**

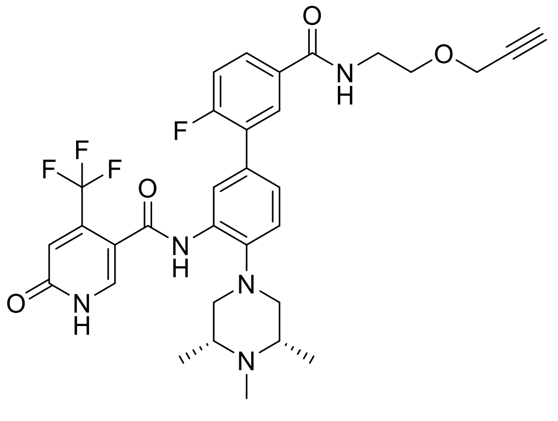

The title compound was prepared according to general procedure A, using 6-fluoro-3'-(6-oxo-4-(trifluoromethyl)-1,6-dihydropyridine-3-carboxamido)-4'-((3*S*,5*R*)-3,4,5-trimethylpiperazin-1-yl)-[1,1'-biphenyl]-3-carboxylic acid (10 mg, 18 µmol), 2-(prop-2-yn-1-yloxy)ethan-1-amine (1.8 mg, 18 µmol), HATU (9.1 mg, 24 µmol) and DIPEA (2.2 µL, 55 µmol). The title compound was obtained as a colourless solid (6.8 mg, 60%). $^{1}H$ NMR (400 MHz, DMSO-$d_6$): δ 12.58 – 11.21 (m, 1H), 9.53 (s, 1H), 8.68 (t, *J* = 5.9 Hz, 1H), 8.01 – 7.95 (m, 2H), 7.92 – 7.86 (m, 1H), 7.47 – 7.38 (m, 2H), 7.31 (s, 1H), 6.83 (s, 1H), 4.17 (d, *J* = 2.4 Hz, 1H), 3.64 – 3.51 (m, 4H), 3.49 – 3.41 (m, 4H), 3.34 (s, 4H), 2.90 (s, 4H), 1.23 (s, 6H). $^{13}C$ NMR (101 MHz, DMSO-$d_6$): δ 165.19, 162.76, 161.51, 161.12, 159.61, 131.14, 130.39, 129.84, 129.61, 128.55, 127.67, 127.56, 126.22, 125.93, 123.11, 120.92, 116.29, 116.10, 111.29, 108.48, 80.27, 77.51, 77.21, 67.84, 67.72, 57.55, 57.36, 43.96, 28.35, 27.24, 13.93. ESI-MS: *m/z* = 628.20 ([M+H]$^+$).

**Synthesis of *N*-(2'-fluoro-5'-(4-(prop-2-yn-1-yl)piperazine-1-carbonyl)-4-((3*S*,5*R*)-3,4,5-trimethylpiperazin-1-yl)-[1,1'-biphenyl]-3-yl)-6-oxo-4-(trifluoromethyl)-1,6-dihydropyridine-3-carboxamide (A11)**

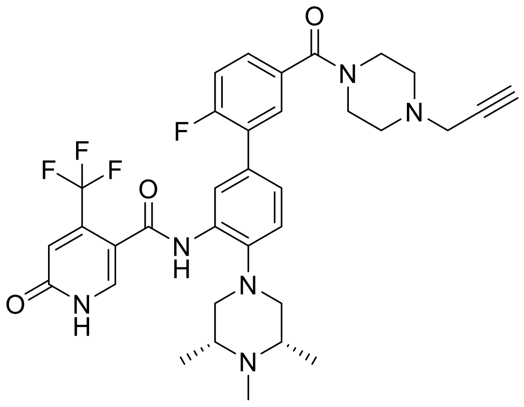


The title compound was prepared according to general procedure A, using 6-fluoro-3'-(6-oxo-4-(trifluoromethyl)-1,6-dihydropyridine-3-carboxamido)-4'-((3*S*,5*R*)-3,4,5-trimethylpiperazin-1-yl)-[1,1'-biphenyl]-3-carboxylic acid (10 mg, 18 µmol), 1-(prop-2-yn-1-yl)piperazine (2.3 mg, 18 µmol), HATU (9.1 mg, 24 µmol) and DIPEA (2.2 µL, 55 µmol). The title compound was obtained as a colourless solid (11.2 mg, 94%). $^{1}H$ NMR (400 MHz, $CD_2Cl_2$): δ 8.92 (s, 1H), 8.51 (s, 1H), 7.82 (s, 1H), 7.52 (dd, *J* = 7.4, 2.2 Hz, 1H), 7.43 – 7.35

(m, 1H), 7.30 (q, *J* = 8.3 Hz, 2H), 7.19 (dd, *J* = 10.3, 8.4 Hz, 1H), 6.88 (s, 1H), 3.76 (s, 2H), 3.51 (s, 2H), 3.34 (d, *J* = 2.5 Hz, 2H), 2.85 (d, *J* = 11.2 Hz, 2H), 2.74 (t, *J* = 11.0 Hz, 2H), 2.55 (s, 6H), 2.35 (s, 3H), 2.33 – 2.29 (m, 1H), 1.13 (s, 3H), 1.12 (s, 3H). $^{13}$C NMR (101 MHz, $CD_2Cl_2$): δ 169.56, 163.39, 162.51, 161.80, 159.81, 141.89, 140.41, 140.14, 138.41, 133.97, 132.93, 132.56, 130.50, 129.50, 129.39, 128.63, 125.87, 123.52, 121.51, 121.09, 120.50, 116.86, 116.67, 114.46, 78.90, 73.82, 59.44, 59.27, 47.18, 17.82. ESI-MS: *m/z* = 653.20 ($[M+H]^+$).

**Synthesis of 3-bromo-5-(bromomethyl)benzoic acid (I7)**

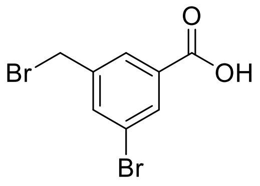


3-bromo-5-methylbenzoic acid (2.0 g, 9.3 mmol), *N*-bromsuccinimid (3.15 g, 17.7 mmol) and benzoyl peroxide (0.11 g, 0.47 mmol) were suspended in dry acetonitrile (120 mL) and stirred at 80 °C for 18 h. Upon cooling, the mixture was concentrated *in vacuo* and subsequently purified by flash chromatography using acetonitrile/water to afford the title compound as a white solid (1.63 g, 60%). $^1$H NMR (300 MHz, DMSO-$d_6$): δ 8.01 (t, *J* = 1.5 Hz, 1H), 7.95 (dt, *J* = 7.0, 1.9 Hz, 2H), 4.78 (s, 2H).

**Synthesis of 3-((*1H*-imidazol-1-yl)methyl)-5-bromobenzoic acid (I8)**

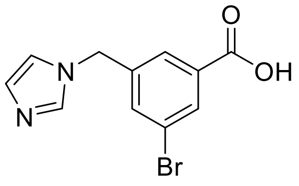


Imidazole (40 mg, 0.59 mmol) was dissolved in dry 1,4-dioxane (5 mL). A solution of **I7** (87 mg, 0.30 mmol) in dry 1,4-dioxane (4 mL) was added dropwise. After stirring for 3 h at 75 °C, the mixture was concentrated *in vacuo* and purified by flash chromatography using acetonitrile/water as an eluent to afford the title compound as a white solid (51 mg, 61%).

$^1H$ NMR (400 MHz, DMSO-$d_6$): δ 9.12 (s, 1H), 8.03 (d, *J* = 1.4 Hz, 1H), 7.97 (dt, *J* = 11.7, 1.6 Hz, 2H), 7.71 (d, *J* = 64.1 Hz, 2H), 5.49 (s, 2H). ESI-MS: *m/z* = 282.95 ($[M+H]^+$).

**Synthesis of 3-((*1H*-imidazol-1-yl)methyl)-5-(1-ethyl-3-(trifluoromethyl)-*1H*-pyrazol-4-yl)benzoic acid (I9)**

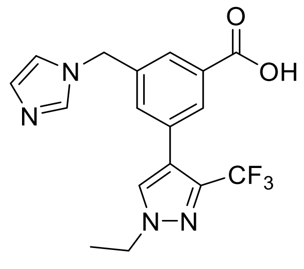


**I8** (250 mg, 0.895 mmol, 1.0 eq), (1-ethyl-3-(trifluoromethyl)-1H-pyrazol-4-yl)boronic acid (370 mg, 1.78 mmol, 2.0 eq), $K_3PO_3$ (475 mg, 2.23 mmol, 2.5 eq) and XPhos Pd G2 (35 mg, 0.045 mmol, 0.05 eq) were dissolved in 1,4-dioxane (8 mL) and $H_2O$ (2 mL). The reaction mixture was stirred at 80 °C for 3 h. Then, the reaction mixture was diluted with $H_2O$ (20 mL) and extracted with $CH_2Cl_2$ (3 x 20 mL). The combined organic phase was washed with brine (20 mL), dried over $MgSO_4$, filtered, and concentrated under reduced pressure. The crude product was purified by flash chromatography using acetonitrile/water as an eluent to afford the title compound (301 mg, 92%). $^1H$ NMR (600 MHz, DMSO-$d_6$): δ 9.29 (s, 1H), 8.32 (s, 1H), 7.98 (dd, *J* = 7.6, 4.8 Hz, 2H), 7.84 (t, *J* = 1.7 Hz, 1H), 7.72 (t, *J* = 1.6 Hz, 1H), 7.66 (s, 1H), 5.56 (s, 2H), 4.26 (q, *J* = 7.3 Hz, 2H), 1.45 (t, *J* = 7.3 Hz, 3H). ESI-MS: *m/z* = 365.05 ($[M+H]^+$).

**Synthesis of *tert*-butyl (*S*)-(1-(6,7-dihydrothieno[3,2-*c*]pyridin-5(*4H*)-yl)-1-oxopent-4-yn-2-yl)carbamate (I10)**

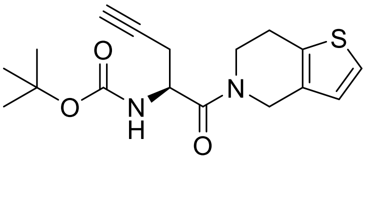

The title compound was prepared according to general procedure A, using (*S*)-2-((tert-butoxycarbonyl)amino)pent-4-ynoic acid (500 mg, 2.34 mmol), 4,5,6,7-tetrahydrothieno[3,2-c]pyridine hydrochloride (412 mg, 2.34 mmol), HATU (1.16 g, 3.04 mmol) and DIPEA (1.22 mL, 7.02 mmol). The title compound was obtained as a colourless solid (715 mg, 91%). $^1$H NMR (400 MHz, $CDCl_3$): δ 7.14 (d, *J* = 5.2 Hz, 1H), 6.79 (dd, *J* = 13.1, 5.1 Hz, 1H), 5.47 (dd, *J* = 26.2, 8.8 Hz, 1H), 4.91 (s, 1H), 4.70 (ddd, *J* = 42.8, 26.5, 16.7 Hz, 2H), 4.10 – 3.75 (m, 2H), 2.90 (dd, *J* = 20.1, 14.6 Hz, 2H), 2.74 – 2.53 (m, 2H), 1.58 (s, 1H), 1.44 (s, *J* = 3.8 Hz, 9H). ESI-MS: *m/z* = 357.05 ($[M+H]^+$).

**Synthesis of (*S*)-3-((1*H*-imidazol-1-yl)methyl)-*N*-(1-(6,7-dihydrothieno[3,2-c]pyridin-5(4*H*)-yl)-1-oxopent-4-yn-2-yl)-5-(1-ethyl-3-(trifluoromethyl)-1*H*-pyrazol-4-yl)benzamide (A12)**

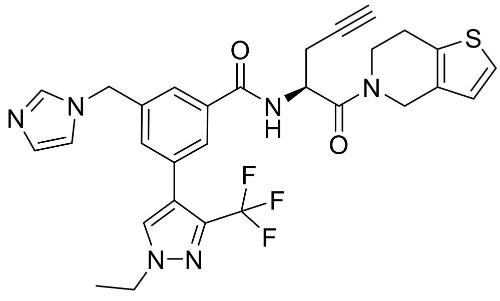

**I10** (333 mg, 0.996 mmol) was dissolved in dry DCM (14 mL). TFA (6 mL) was added and the reaction mixture was stirred at rt for 1 h. All volatiles were removed under reduced pressure and the crude product was dissolved in dry DMF (20 mL). **I9** (363 mg, 0.996 mmol), HATU (492 mg, 1.29 mmol) and DIPEA (520 µL, 2.99 mmol) were added to the solution and stirred at rt for two h. The crude product was purified by flash chromatography using acetonitrile/water as an eluent to afford the title compound as a light brown solid (307 mg, 53%). $^1$H NMR (600 MHz, DMSO-$d_6$): δ 8.93 (dd, *J* = 73.5, 8.4 Hz, 1H), 8.20 (d, *J* = 11.9 Hz, 1H), 7.89 – 7.71 (m, 3H), 7.41 – 7.27 (m, 2H), 7.19 (d, *J* = 6.3 Hz, 1H), 6.94 – 6.80 (m, 2H), 5.28 (d, *J* = 11.2 Hz, 2H), 5.18 (ddd, *J* = 44.8, 15.1, 8.0 Hz, 1H), 4.75 – 4.43 (m, 2H), 4.25 (q, *J* = 7.3 Hz, 2H), 3.94 – 3.69 (m, 2H), 2.92 – 2.58 (m, 5H), 1.43 (dd, *J* = 8.4, 6.2 Hz, 3H). $^{13}$C NMR (151 MHz, DMSO-$d_6$): δ 168.50, 165.23, 138.39, 136.32, 134.40, 133.06, 132.37, 131.89, 131.48, 131.04, 130.17, 126.80, 125.98, 125.34, 124.36,

123.64, 122.55, 120.76, 119.83, 80.92, 72.59, 49.20, 48.48, 47.23, 42.95, 42.60, 25.11, 21.43, 15.04. ESI-MS: *m/z* = 581.15 ($[M+H]^+$).

**Synthesis of *N*-(but-3-yn-1-yl)-4-(3-chloro-2-fluorophenoxy)-1-((6-(thiazol-2-ylamino)pyridin-2-yl)methyl)cyclohexane-1-carboxamide (A13)**

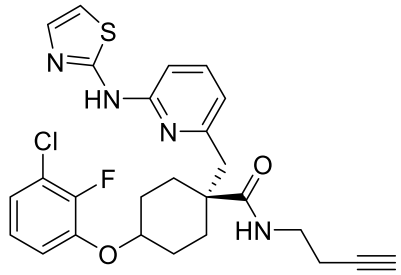

The title compound was prepared according to general procedure A, using 4-(3-chloro-2-fluorophenoxy)-1-((6-(thiazol-2-ylamino)pyridin-2-yl)methyl)cyclohexane-1-carboxylic acid (10 mg, 22 µmol), but-3-yn-1-amine (1.5 mg, 22 µmol), HATU (10.7 mg, 28.1 µmol) and DIPEA (11.3 µL, 65.9 µmol). The title compound was obtained as a colourless solid (10.1 mg, 91%). $^1H$ NMR (500 MHz, DMSO-$d_6$): δ 11.12 (s, 1H), 7.78 (t, *J* = 5.7 Hz, 1H), 7.55 (d, *J* = 7.8 Hz, 1H), 7.36 (d, *J* = 3.6 Hz, 1H), 7.18 – 7.13 (m, 1H), 7.10 (s, 1H), 7.09 (d, *J* = 3.2 Hz, 1H), 6.95 (d, *J* = 3.6 Hz, 1H), 6.86 (d, *J* = 8.2 Hz, 1H), 6.62 (d, *J* = 7.3 Hz, 1H), 4.55 (s, 1H), 3.14 (q, *J* = 7.1, 5.4 Hz, 2H), 2.89 (s, 2H), 2.79 (t, *J* = 2.6 Hz, 1H), 2.26 (td, *J* = 7.2, 2.7 Hz, 2H), 1.93 – 1.87 (m, 2H), 1.83 – 1.74 (m, 4H), 1.72 – 1.63 (m, 2H). $^{13}C$ NMR (125 MHz, DMSO-$d_6$): δ 174.22, 159.88, 155.12, 150.81, 149.94, 147.98, 146.21, 137.61, 137.44, 124.98, 121.90, 120.54, 120.42, 116.74, 116.27, 110.73, 108.17, 82.49, 74.29, 72.03, 46.67, 38.13, 28.31, 26.62, 18.56. ESI-MS: *m/z* = 513.10 ($[M+H]^+$).

**Synthesis of 4-(3-chloro-2-fluorophenoxy)-*N*-(hex-5-yn-1-yl)-1-((6-(thiazol-2-ylamino)pyridin-2-yl)methyl)cyclohexane-1-carboxamide (A14)**

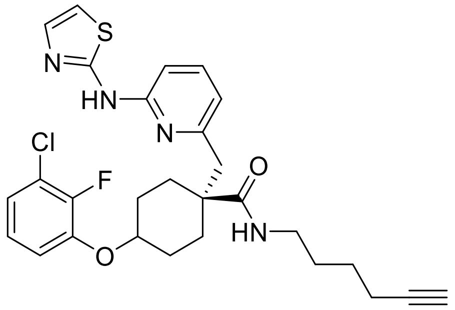

The title compound was prepared according to general procedure A, using 4-(3-chloro-2-fluorophenoxy)-1-((6-(thiazol-2-ylamino)pyridin-2-yl)methyl)cyclohexane-1-carboxylic acid (10 mg, 22 µmol), hex-5-yn-1-amine (2.1 mg, 22 µmol), HATU (10.7 mg, 28.1 µmol) and DIPEA (11.3 µL, 65.9 µmol). The title compound was obtained as a colourless solid (7.1 mg, 61%). $^{1}$H NMR (500 MHz, DMSO-$d_6$): δ 11.11 (s, 1H), 7.60 (t, *J* = 5.6 Hz, 1H), 7.54 (dd, *J* = 8.3, 7.3 Hz, 1H), 7.35 (d, *J* = 3.6 Hz, 1H), 7.21 – 7.13 (m, 1H), 7.10 (s, 1H), 7.09 (d, *J* = 3.0 Hz, 1H), 6.95 (d, *J* = 3.6 Hz, 1H), 6.86 (d, *J* = 8.2 Hz, 1H), 6.59 (d, *J* = 7.3 Hz, 1H), 4.55 (s, 1H), 3.03 (q, *J* = 6.5 Hz, 2H), 2.90 (s, 2H), 2.75 (t, *J* = 2.6 Hz, 1H), 2.14 (td, *J* = 6.9, 2.7 Hz, 2H), 1.94 – 1.87 (m, 2H), 1.84 – 1.74 (m, 4H), 1.69 – 1.61 (m, 2H), 1.51 – 1.42 (m, 2H), 1.42 – 1.33 (m, 2H). $^{13}$C NMR (125 MHz, DMSO-$d_6$): δ 173.99, 159.88, 155.26, 150.80, 149.91, 147.95, 146.22, 137.57, 137.44, 125.01, 121.87, 120.53, 120.41, 116.70, 116.20, 110.72, 108.13, 84.52, 74.35, 71.30, 46.61, 38.37, 28.40, 28.29, 26.73, 25.56, 17.46. ESI-MS: *m/z* = 541.15 ([M+H]$^+$).

**Synthesis of 4-(3-chloro-2-fluorophenoxy)-*N*-(2-(prop-2-yn-1-yloxy)ethyl)-1-((6-(thiazol-2-ylamino)pyridin-2-yl)methyl)cyclohexane-1-carboxamide (A15)**

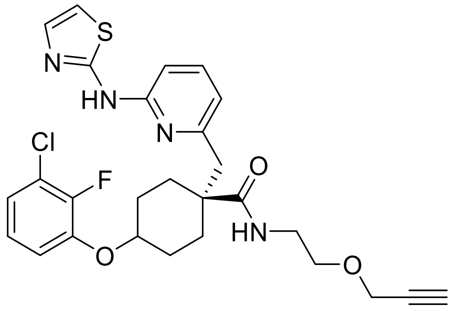

The title compound was prepared according to general procedure A, using 4-(3-chloro-2-fluorophenoxy)-1-((6-(thiazol-2-ylamino)pyridin-2-yl)methyl)cyclohexane-1-carboxylic acid (10 mg, 22 µmol), 2-(prop-2-yn-1-yloxy)ethan-1-amine (2.2 mg, 22 µmol), HATU (10.7 mg, 28.1 µmol) and DIPEA (11.3 µL, 65.9 µmol). The title compound was obtained as a colourless solid (7.6 mg, 65%). $^{1}$H NMR (500 MHz, DMSO-$d_6$): δ 11.11 (s, 1H), 7.70 (t, *J* = 5.7 Hz, 1H), 7.55 (dd, *J* = 8.2, 7.3 Hz, 1H), 7.35 (d, *J* = 3.6 Hz, 1H), 7.17 – 7.13 (m, 1H), 7.10 (s, 1H), 7.09 (d, *J* = 3.0 Hz, 1H), 6.95 (d, *J* = 3.6 Hz, 1H), 6.86 (d, *J* = 8.1 Hz, 1H), 6.63 (dd, *J* = 7.4, 0.8 Hz, 1H), 4.55 (s, 1H), 4.12 (d, *J* = 2.4 Hz, 2H), 3.44 (t, *J* = 2.4 Hz, 1H), 3.42 (d, *J* = 6.0 Hz, 2H), 3.21 (q, *J* = 6.0 Hz, 2H), 2.90 (s, 2H), 1.91 – 1.85 (m, 2H), 1.84 – 1.73 (m, 4H), 1.70 – 1.61 (m, 2H). $^{13}$C NMR (125 MHz, DMSO-$d_6$): δ 74.27, 159.89, 155.17, 150.78, 149.93, 147.98, 146.30, 146.22, 137.63, 137.44, 125.02, 121.88, 120.53, 120.41, 116.74, 116.20, 110.73, 108.14, 80.37, 77.21, 74.33, 67.78, 57.32, 46.69, 38.60, 28.38, 26.63. ESI-MS: *m/z* = 543.15 ([M+H]$^+$).

**Synthesis of 4-(3-chloro-2-fluorophenoxy)-*N*-(2-(2-(prop-2-yn-1-yloxy)ethoxy)ethyl)-1-((6-(thiazol-2-ylamino)pyridin-2-yl)methyl)cyclohexane-1-carboxamide (A16)**

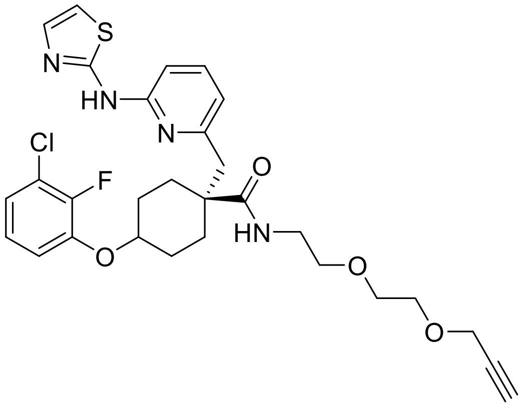


The title compound was prepared according to general procedure A, using 4-(3-chloro-2-fluorophenoxy)-1-((6-(thiazol-2-ylamino)pyridin-2-yl)methyl)cyclohexane-1-carboxylic acid (10 mg, 22 µmol), 2-(2-(prop-2-yn-1-yloxy)ethoxy)ethan-1-amine (3.1 mg, 22 µmol), HATU (10.7 mg, 28.1 µmol) and DIPEA (11.3 µL, 65.9 µmol). The title compound was obtained as a colourless solid (7.4 mg, 58%). $^{1}$H NMR (500 MHz, DMSO-$d_6$): δ 11.11 (s, 1H), 7.64 (t, *J* = 5.6 Hz, 1H), 7.54 (dd, *J* = 8.2, 7.3 Hz, 1H), 7.35 (d, *J* = 3.6 Hz, 1H), 7.19 – 7.13 (m, 1H), 7.11 (s, 1H), 7.09 (d, *J* = 2.9 Hz, 1H), 6.95 (d, *J* = 3.6 Hz, 1H), 6.86 (d, *J* = 7.9 Hz, 1H), 6.63 (d, *J* = 7.2 Hz, 1H), 4.55 (s, 1H), 4.13 (d, *J* = 2.4 Hz, 2H), 3.61 – 3.47 (m, 4H), 3.40 (t, *J* = 2.4 Hz, 1H), 3.38 – 3.32 (m, 2H), 3.19 (q, *J* = 6.0 Hz, 2H), 2.90 (s, 2H), 1.94 – 1.86

(m, 2H), 1.84 – 1.74 (m, 4H), 1.71 – 1.62 (m, 2H). $^{13}$C NMR (125 MHz, DMSO-$d_6$): δ 174.21, 159.89, 155.20, 150.78, 149.94, 147.98, 146.31, 146.23, 137.62, 137.43, 125.02, 124.98, 121.89, 120.53, 120.41, 116.74, 116.22, 110.73, 108.14, 80.37, 77.15, 74.37, 69.28, 68.89, 68.55, 57.57, 46.67, 38.78, 28.42, 26.64. ESI-MS: *m/z* = 587.15 ([M+H]$^+$).

**Synthesis of *tert*-butyl 2-((2-(2,6-dioxopiperidin-3-yl)-1,3-dioxoisoindolin-4-yl)oxy)acetate (I11)**

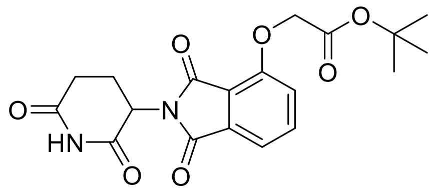

2-(2,6-dioxopiperidin-3-yl)-4-hydroxyisoindoline-1,3-dione (1.00 g, 3.65 mmol, 1.0 eq) and potassium carbonate (1.26 g, 9.12 mmol, 3.0 eq) were suspended in dry DMF (15 mL). The flask was cooled to 0 °C in an ice bath and *tert*-butyl bromoacetate (538 µL, 3.65 mmol, 1.0 eq) was added dropwise through a dropping funnel over 30 minutes. The solution was warmed to ambient temperature and stirred for two h. The reaction mixture was diluted with cold water and the precipitate was filtered. The colourless solid was dried under reduced pressure to obtain the title compound (1.21 g, 85%). $^1$H NMR (400 MHz, DMSO-$d_6$) δ 11.11 (s, 1H), 7.80 (t, *J* = 7.9 Hz, 1H), 7.48 (d, *J* = 7.3 Hz, 1H), 7.37 (d, *J* = 8.5 Hz, 1H), 5.10 (dd, *J* = 13.0, 5.3 Hz, 1H), 4.96 (s, 2H), 2.89 (t, *J* = 19.1, 14.0, 5.3 Hz, 1H), 2.58 (t, 2H), 2.15 – 1.92 (m, 1H), 1.42 (s, 9H). ESI-MS: *m/z* = 411.10 ([M+Na]$^+$).

**Synthesis of 2-((2-(2,6-dioxopiperidin-3-yl)-1,3-dioxoisoindolin-4-yl)oxy)acetic acid (I12)**

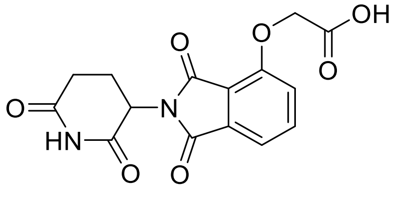

**I11** (1.00 g, 3.02 mmol) was dissolved in dry DCM (5 mL). The flask was cooled to 0 °C in an ice bath and trifluoroacetic acid (TFA) in an excess was added dropwise. The solution was stirred at rt for two h. The solvent was removed under reduced pressure and the crude product used without further purification. ESI-MS: *m/z* = 333.10 ([M+H]⁺).

**Synthesis of *N*-(3-azidopropyl)-2-((2-(2,6-dioxopiperidin-3-yl)-1,3-dioxoisoindolin-4-yl)-oxy)acetamide (X1)**

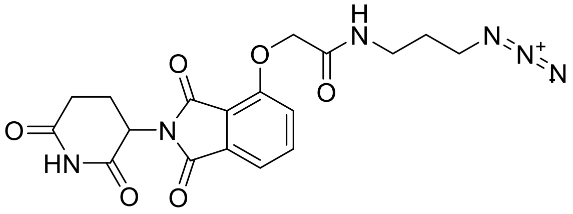

The title compound was prepared according to general procedure A, using **I12** (100 mg, 0.304 mmol), 3-azidopropan-1-amine (30 mg, 0.30 mmol), HATU (150 mg, 0.395 mmol) and DIPEA (158 µL, 0.912 mmol). The title compound was obtained as a light yellow solid (114 mg, 91%). $^1$H NMR (400 MHz, DMSO-$d_6$) δ 11.11 (s, 1H), 8.05 (t, *J* = 5.8 Hz, 1H), 7.81 (dd, *J* = 8.5, 7.3 Hz, 1H), 7.50 (d, *J* = 7.2 Hz, 1H), 7.39 (d, *J* = 8.5 Hz, 1H), 5.12 (dd, *J* = 12.9, 5.4 Hz, 1H), 4.78 (s, 2H), 3.37 (t, *J* = 6.8 Hz, 2H), 3.22 (q, *J* = 6.5 Hz, 2H), 2.90 (m, 1H), 2.63 – 2.51 (m, 2H), 2.16 – 1.99 (m, 1H), 1.69 (quin, *J* = 6.8 Hz, 2H). $^{13}$C NMR (101 MHz, DMSO-$d_6$) δ 172.80, 169.90, 166.97, 166.76, 165.52, 155.12, 136.96, 133.07, 120.45, 116.88, 116.10, 67.72, 48.83, 48.34, 35.80, 30.96, 28.33, 22.00. ESI-MS: *m/z* = 437.10 ([M+Na]⁺).

**Synthesis of *N*-(2-(2-(2-azidoethoxy)ethoxy)ethyl)-2-((2-(2,6-dioxopiperidin-3-yl)-1,3-dioxoisoindolin-4-yl)oxy)acetamide (X2)**

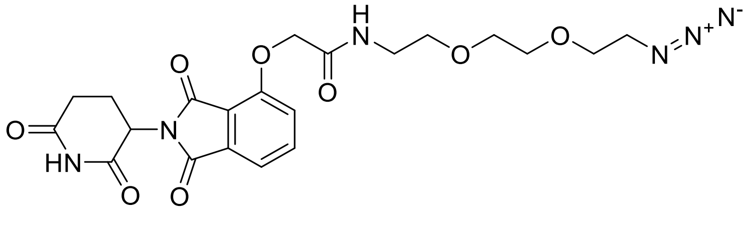

The title compound was prepared according to general procedure B, using **I12** (350 mg, 1.05 mmol) and 1-(2-aminoethoxy)-2-(2-azidoethoxy)ethane (219 mg, 1.26 mmol), 1-methyl-1*H*-imidazole (293 µL, 3.68 mmol) and TCFH (353 mg, 1.26 mmol). The title compound was obtained as a colourless solid (315 mg, 62%). $^{1}$H NMR (500 MHz, Acetone-$d_6$): δ 9.95 (s, 1H), 7.87 (dd, *J* = 8.4, 7.3 Hz, 1H), 7.63 (s, 1H), 7.55 – 7.48 (m, 2H), 5.19 – 5.11 (m, 1H), 4.75 (s, 2H), 3.71 – 3.56 (m, 8H), 3.48 (q, *J* = 5.5 Hz, 2H), 3.35 (t, *J* = 5.0 Hz, 2H), 3.04 – 2.93 (m, 1H), 2.83 – 2.74 (m, 2H), 2.29 – 2.19 (m, 1H). $^{13}$C NMR (126 MHz, Acetone-$d_6$): δ 171.7, 169.1, 166.7, 166.6, 165.9, 154.9, 137.0, 133.7, 120.3, 117.8, 116.3, 70.3, 70.2, 69.9, 69.3, 69.3, 68.0, 50.5, 49.3, 38.7, 31.1, 29.4, 29.3, 29.1, 29.0, 28.8, 28.6, 28.5, 22.4. ESI-MS: *m/z* = 489.2 ([M+H]$^+$).

**Synthesis of *N*-(3-azidopropyl)-2-((2-(2,6-dioxopiperidin-3-yl)-1,3-dioxoisoindolin-4-yl)-oxy)acetamide (X3)**

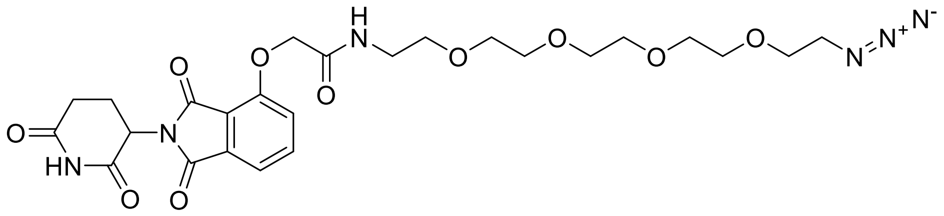

The title compound was prepared according to general procedure B, using **I12** (360 mg, 1.08 mmol) and 14-azido-3,6,9,12-tetraoxatetradecan-1-amine (283 µL, 1.19 mmol), 1-methyl-1*H*-imidazole (301 µL, 3.78 mmol) and TCFH (363 mg, 1.30 mmol). The title compound was obtained as a white solid (460 mg, 73%). $^{1}$H NMR (400 MHz, $CD_2Cl_2$): δ 8.56 (s, 1H), 7.67 (dd, *J* = 8.4, 7.3 Hz, 1H), 7.49 – 7.42 (m, 2H), 7.15 (dd, *J* = 8.4, 0.7 Hz, 1H), 4.92 – 4.82 (m, 1H), 4.56 (s, 2H), 3.60 – 3.49 (m, 16H), 3.45 (quin, *J* = 5.2 Hz, 2H), 3.29 (t, 2H), 2.81 – 2.60 (m, 3H), 2.13 – 2.02 (m, 1H). $^{13}$C NMR (101 MHz, $CD_2Cl_2$): δ 171.0, 168.3, 166.7, 166.6, 165.9, 154.5, 136.9, 133.7, 119.5, 118.0, 117.0, 70.6, 70.5, 70.4, 70.3, 69.8, 69.5, 68.1, 50.8, 49.3, 39.0, 31.4, 22.6. ESI-MS: *m/z* = 577.2 ([M+H]$^+$).

**Synthesis of 4-((3-azidopropyl)amino)-2-(2,6-dioxopiperidin-3-yl)isoindoline-1,3-dione (X4)**

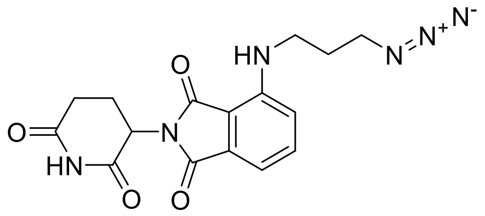

The title compound was prepared according to general procedure C, using 2-(2,6-dioxopiperidin-3-yl)-4-fluoroisoindoline-1,3-dione (150 mg, 0.543 mmol), 3-azidopropan-1-amine (65 mg, 0.65 mmol) and DIPEA (283 µL, 1.63 mmol). The title compound was obtained as a yellow solid (155 mg, 80%). $^{1}$H NMR (400 MHz, DMSO-$d_6$) δ 11.09 (s, 1H), 7.59 (dd, *J* = 8.6, 7.1 Hz, 1H), 7.11 (d, *J* = 8.6 Hz, 1H), 7.03 (d, *J* = 7.0 Hz, 1H), 6.67 (t, *J* = 6.1 Hz, 1H), 5.05 (dd, *J* = 12.9, 5.4 Hz, 1H), 3.45 (t, *J* = 6.7 Hz, 2H), 3.38 (q, *J* = 6.7 Hz, 2H), 2.95 – 2.81 (m, 1H), 2.64 – 2.53 (m, 2H), 2.08 – 1.94 (m, 1H), 1.83 (quin, *J* = 6.8 Hz, 2H). $^{13}$C NMR (101 MHz, DMSO-$d_6$) δ 172.83, 170.10, 168.80, 167.30, 146.22, 136.30, 132.27, 117.12, 110.56, 109.33, 48.51, 39.42, 30.98, 27.93, 22.16. ESI-MS: *m/z* = 357.15 ([M+H]$^+$).

**Synthesis of 4-((4-azidobutyl)amino)-2-(2,6-dioxopiperidin-3-yl)isoindoline-1,3-dione (X5)**

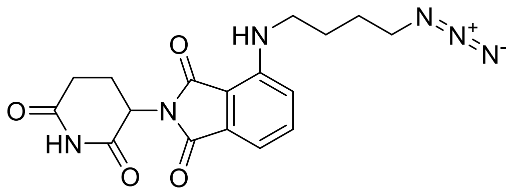

The title compound was prepared according to general procedure C, using 2-(2,6-dioxopiperidin-3-yl)-4-fluoroisoindoline-1,3-dione (200 mg, 0.724 mmol), 4-azidobutan-1-amine (99 mg, 0.87 mmol) and DIPEA (378 µL, 2.17 mmol). The title compound was obtained as a yellow solid (196 mg, 73%). $^{1}$H NMR (400 MHz, DMSO-$d_6$) δ 11.09 (s, 1H), 7.58 (dd, *J* = 8.6, 7.1 Hz, 1H), 7.11 (d, *J* = 8.6 Hz, 1H), 7.02 (d, *J* = 7.0 Hz, 1H), 6.61 (t, *J* = 6.1 Hz, 1H), 5.05 (dd, *J* = 12.9, 5.4 Hz, 1H), 3.41 – 3.33 (m, 4H), 2.96 – 2.81 (m, 1H), 2.62 – 2.52 (m, 2H), 2.06 – 1.96 (m, 1H), 1.66 – 1.59 (m, 4H). $^{13}$C NMR (101 MHz, DMSO-$d_6$) δ 172.81, 170.10, 168.89, 167.30, 146.33, 136.26, 132.24, 117.22, 110.44, 109.13, 50.37, 41.27, 39.94, 30.98, 25.91, 25.70, 22.15. ESI-MS: *m/z* = 371.10 ([M+H]$^+$).

**Synthesis of 4-((6-azidohexyl)amino)-2-(2,6-dioxopiperidin-3-yl)isoindoline-1,3-dione (X6)**

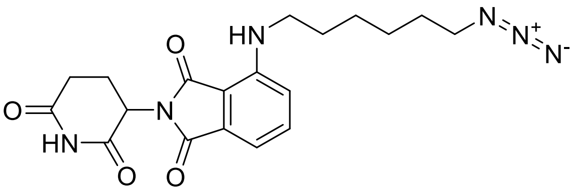


The title compound was prepared according to general procedure C, using 2-(2,6-dioxopiperidin-3-yl)-4-fluoroisoindoline-1,3-dione (200 mg, 0.724 mmol), 6-azidohexan-1-amine (123 mg, 0.868 mmol) and DIPEA (378 µL, 2.17 mmol). The title compound was obtained as a yellow solid (195 mg, 68%). $^1$H NMR (400 MHz, DMSO-$d_6$) δ 11.09 (s, 1H), 7.57 (dd, $J$ = 8.6, 7.1 Hz, 1H), 7.08 (d, $J$ = 8.6 Hz, 1H), 7.02 (d, $J$ = 7.0 Hz, 1H), 6.53 (t, $J$ = 6.0 Hz, 1H), 5.05 (dd, $J$ = 12.9, 5.4 Hz, 1H), 3.33 – 3.25 (m, 4H), 2.95 – 2.77 (m, 1H), 2.64 – 2.51 (m, 2H), 2.06 – 1.96 (m, 1H), 1.68 – 1.49 (m, 4H), 1.36 (quin, $J$ = 3.5 Hz, 4H). $^{13}$C NMR (101 MHz, DMSO-$d_6$) δ 172.81, 170.10, 168.96, 167.31, 146.42, 136.27, 132.20, 117.17, 110.38, 109.04, 50.57, 48.54, 41.75, 30.98, 28.54, 28.17, 25.88, 22.15. ESI-MS: $m/z$ = 399.20 ([M+H]$^+$).

**Synthesis of 4-({2-[2-(2-Azidoethoxy)ethoxy]ethyl}amino)-2-(2,6-dioxopiperidin-3-yl)-2,3-dihydro-1*H*-isoindole-1,3-dione (X7)**

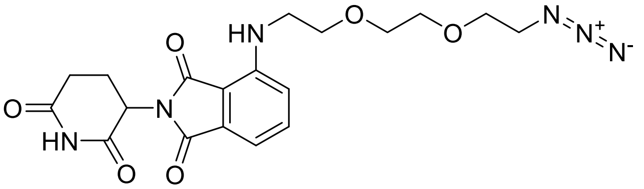


2-(2,6-dioxopiperidin-3-yl)-4-fluoroisoindoline-1,3-dione (231 mg, 0.821 mmol, 1.0 eq) and 1-(2-aminoethoxy)-2-(2-azidoethoxy)ethane (157 mg, 0.903 mmol, 1.1 eq) were dissolved in DMF (4 mL). *N*,*N*-Diisopropylethylamine (0.281 µL, 1.64 mmol, 2.0 eq) was added and the reaction mixture was heated up to 90 °C for 14 h. After TLC confirmed completion of the reaction, EtOAc (50 mL) was added to the mixture and the solution was washed with brine (2 x 50 mL) and water (50 mL). The organic phase was dried over $MgSO_4$ and the solvent was removed under reduced pressure. The crude product was purified by flash chromatography using acetonitrile/water as an eluent to obtain the title compound as a yellow solid (220 mg, 62%). $^1$H NMR (400 MHz, Acetone-$d_6$) δ 9.98 – 9.75 (m, 1H), 7.63 – 7.55 (m, 1H), 7.17 – 7.11 (m, 1H), 7.08 – 7.01 (m, 1H), 6.81 – 6.50 (m, 1H), 5.11 – 5.02 (m, 1H), 3.80 – 3.73 (m, 2H), 3.72 – 3.62 (m, 5H), 3.59 – 3.51 (m, 2H), 3.43 – 3.31 (m, 2H), 3.03 – 2.88 (m, 1H), 2.82 – 2.72 (m, 4H), 2.28 – 2.15 (m, 1H). $^{13}$C NMR (101 MHz, Acetone-$d_6$) δ 172.65, 170.28, 170.20, 168.28, 147.84, 136.87, 133.63, 117.86, 111.48, 111.14, 71.35, 71.21, 70.82, 70.24, 51.42, 49.89, 43.04, 32.02, 23.43. ESI-MS: *m/z* = 431.2 ([M+H]$^+$).

**Synthesis of 4-(15-Azido-4,7,10,13-tetraoxa-1-azapentadecan-1-yl)-2-(2,6-dioxopiperidin-3-yl)-2,3-dihydro-1*H*-isoindole-1,3-dione (X8)**

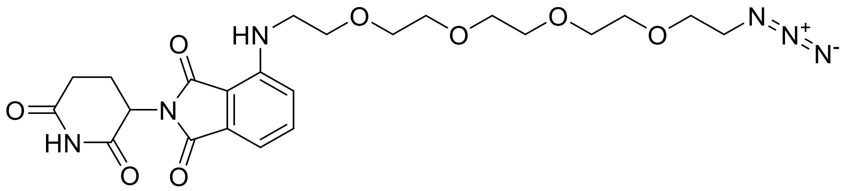


2-(2,6-dioxopiperidin-3-yl)-4-fluoroisoindoline-1,3-dione (556 mg, 1.97 mmol, 1.0 eq) and 1-(2-aminoethoxy)-2-(2-azidoethoxy)ethane (569 mg, 2.18 mmol, 1.1 eq) were dissolved in DMF (10 mL). DIPEA (728 µL, 4.26 mmol, 2.2 eq) was added and the reaction mixture was heated up to 90 °C for 14 h. After TLC confirmed completion of the reaction, EtOAc (50 mL) was added to the mixture and the solution was washed with brine (2 x 50 mL) and water (50 mL). The organic phase was dried over $MgSO_4$ and the solvent was removed under reduced pressure. The crude product was purified by flash chromatography using acetonitrile/water as an eluent to obtain the title compound as a yellow solid (338 mg, 33%). $^1$H NMR (500 MHz, Acetone-$d_6$) δ 9.86 (s, 1H), 7.63 – 7.56 (m, 1H), 7.14 (d, *J* = 8.5 Hz, 1H), 7.05 (dd, *J* = 7.1, 0.6 Hz, 1H), 6.62 (t, *J* = 5.7 Hz, 1H), 5.07 (dd, *J* = 12.6, 5.4 Hz, 1H), 3.75 (t, *J* = 5.4 Hz, 2H), 3.68 – 3.65 (m, 2H), 3.65 – 3.62 (m, 4H), 3.60 – 3.60 (m, 4H), 3.59 – 3.58 (m, 4H), 3.54 (q, *J* = 5.5 Hz, 2H), 3.37 (t, *J* = 5.0 Hz, 2H), 3.03 – 2.89 (m, 1H), 2.80 – 2.72 (m, 2H), 2.28 – 2.12 (m, 1H). $^{13}$C NMR (126 MHz, Acetone-$d_6$) δ 172.66, 170.25, 170.20, 168.28, 147.85, 136.88, 133.63, 117.87, 111.46, 111.11, 71.34,

71.31, 71.28, 71.20, 70.69, 70.22, 51.40, 49.89, 43.08, 32.03, 23.44. ESI-MS: *m/z* = 519.2 ($[M+H]^+$).

**Synthesis of *tert*-butyl (2-((2-(2,6-dioxopiperidin-3-yl)-1,3-dioxoisoindolin-4-yl)amino)-ethyl)carbamate (I13)**

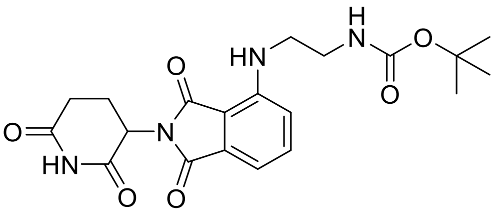

The title compound was prepared according to general procedure C, using 2-(2,6-dioxopiperidin-3-yl)-4-fluoroisoindoline-1,3-dione (1.30 mg, 4.71 mmol), *tert*-butyl (2-aminoethyl)carbamate (905 mg, 5.65 mmol) and DIPEA (2.46 mL, 14.1 mmol). The title compound was obtained as a yellow solid (899 mg, 46%). $^1$H NMR (400 MHz, DMSO-$d_6$) δ 11.08 (s, 1H), 7.58 (dd, *J* = 8.6, 7.1 Hz, 1H), 7.14 (d, *J* = 8.6 Hz, 1H), 7.06 – 6.95 (m, 2H), 6.71 (t, *J* = 6.2 Hz, 1H), 5.05 (dd, *J* = 12.5, 5.4 Hz, 1H), 3.41 – 3.34 (m, 2H), 3.12 (q, *J* = 6.1 Hz, 2H), 2.98 – 2.79 (m, 1H), 2.64 – 2.53 (m, 2H), 2.06 – 1.95 (m, 1H), 1.36 (s, 9H). ESI-MS: *m/z* = 439.10 ($[M+Na]^+$).

**Synthesis of 4-((2-aminoethyl)amino)-2-(2,6-dioxopiperidin-3-yl)isoindoline-1,3-dione (I14)**

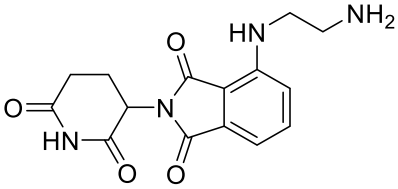

The title compound was prepared according to general procedure D, using **I13** (169 mg, 0.406 mmol). The crude product was used without further purification. ESI-MS: *m/z* = 317.10 ($[M+H]^+$).

**Synthesis of 2-azido-*N*-(2-((2-(2,6-dioxopiperidin-3-yl)-1,3-dioxoisoindolin-4-yl)amino)-ethyl)acetamide (X9)**

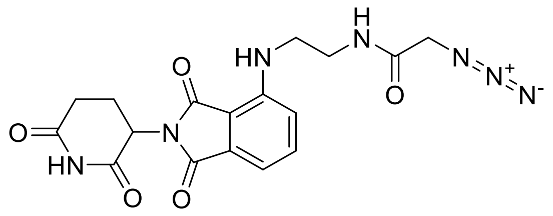

The title compound was prepared according to general procedure A, using **I14** (128 mg, 0.406 mmol), 2-azidoacetic acid (30.4 µL, 0.406 mmol), HATU (185 mg, 0.487 mmol) and DIPEA (212 µL, 1.22 mmol). The title compound was obtained as a yellow solid (93 mg, 57%). $^{1}$H NMR (400 MHz, DMSO-$d_6$) δ 11.09 (s, 1H), 8.33 (t, *J* = 5.6 Hz, 1H), 7.59 (dd, *J* = 8.6, 7.0 Hz, 1H), 7.17 (d, *J* = 8.6 Hz, 1H), 7.04 (d, *J* = 7.0 Hz, 1H), 6.74 (t, *J* = 6.2 Hz, 1H), 5.06 (dd, *J* = 12.9, 5.4 Hz, 1H), 3.82 (s, 2H), 3.42 (q, *J* = 6.3 Hz, 2H), 3.32 – 3.26 (m, 2H), 2.96 – 2.78 (m, 1H), 2.63 – 2.51 (m, 2H), 2.10 – 1.89 (m, 1H). $^{13}$C NMR (101 MHz, DMSO-$d_6$) δ 172.83, 170.11, 168.72, 167.79, 167.30, 146.29, 136.24, 132.25, 117.12, 110.65, 109.37, 50.83, 48.54, 41.22, 38.19, 30.98, 22.16. ESI-MS: *m/z* = 400.20 ([M+H]$^+$).

**Synthesis of *tert*-butyl 2-((2-(2,6-dioxopiperidin-3-yl)-1,3-dioxoisoindolin-5-yl)oxy)acetate (I15)**

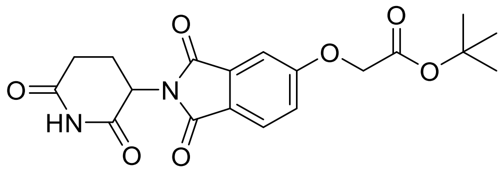

2-(2,6-dioxopiperidin-3-yl)-5-hydroxyisoindoline-1,3-dione (1.00 g, 3.65 mmol, 1 eq) and potassium carbonate (1.26 g, 9.12 mmol, 3 eq) were suspended in dry DMF (15 mL). The flask was cooled to 0 °C in an ice bath and *tert*-butyl bromoacetate (538 µL, 3.65 mmol, 1 eq) was added dropwise through a dropping funnel over 30 minutes. The solution was warmed to ambient temperature and stirred for two h. The reaction mixture was diluted with cold water and the precipitate was filtered. The colourless solid was dried under reduced pressure to obtain the title compound (1.13 g, 80%). $^{1}$H NMR (400 MHz, DMSO-$d_6$) δ 11.11 (s, 1H), 7.85 (d, *J* = 8.2 Hz, 1H), 7.40 (s, 1H), 7.35 (dd, *J* = 8.3, 2.4 Hz, 1H), 5.11

(dd, *J* = 12.9, 5.4 Hz, 1H), 4.93 (s, 2H), 2.97 – 2.81 (m, 1H), 2.65 – 2.53 (m, 2H), 2.13 – 2.01 (m, 1H), 1.43 (s, 9H). ESI-MS: *m/z* = 389.10 ($[M+H]^+$).

**Synthesis of 2-((2-(2,6-dioxopiperidin-3-yl)-1,3-dioxoisoindolin-5-yl)oxy)acetic acid (I16)**

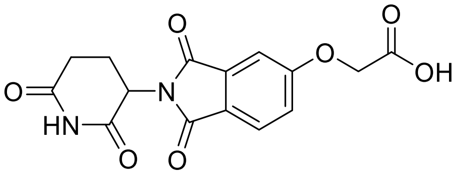


**I15** (1.00 g, 3.02 mmol) was dissolved in dry DCM (5 mL). The flask was cooled to 0 °C in an ice bath and trifluoroacetic acid (TFA) in an excess was added dropwise. The solution was stirred at rt for two h. The solvent was removed under reduced pressure and the crude product used without further purification. ESI-MS: *m/z* = 333.05 ($[M+H]^+$).

**Synthesis of *N*-(3-azidopropyl)-2-((2-(2,6-dioxopiperidin-3-yl)-1,3-dioxoisoindolin-5-yl)-oxy)acetamide (X10)**

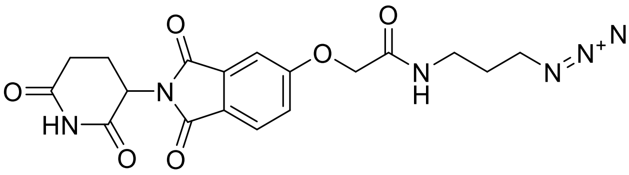


The title compound was prepared according to general procedure A, using **I16** (100 mg, 0.304 mmol), 3-azidopropan-1-amine (30 mg, 0.30 mmol), HATU (150 mg, 0.395 mmol) and DIPEA (158 µL, 0.912 mmol). The title compound was obtained as a light yellow solid (121 mg, 96%). $^1H$ NMR (400 MHz, DMSO-$d_6$) δ 11.10 (s, 1H), 8.28 (t, *J* = 5.8 Hz, 1H), 7.87 (d, *J* = 8.3 Hz, 1H), 7.44 (d, *J* = 2.3 Hz, 1H), 7.39 (dd, *J* = 8.3, 2.3 Hz, 1H), 5.12 (dd, *J* = 12.9, 5.4 Hz, 1H), 4.73 (s, 2H), 3.35 (t, *J* = 6.8 Hz, 2H), 3.20 (q, *J* = 6.5 Hz, 2H), 2.98 – 2.82 (m, 1H), 2.66 – 2.51 (m, 2H), 2.09 – 2.01 (m, 1H), 1.69 (quin, *J* = 6.8 Hz, 2H). $^{13}C$ NMR (101 MHz, DMSO-$d_6$) δ 172.78, 169.91, 166.90, 166.74, 163.07, 133.73, 125.32, 123.56, 120.91, 109.42, 67.36, 49.00, 48.37, 35.78, 30.95, 28.36, 22.05. ESI-MS: *m/z* = 437.10 ($[M+Na]^+$).

**Synthesis of *N*-(2-(2-(2-azidoethoxy)ethoxy)ethyl)-2-((2-(2,6-dioxopiperidin-3-yl)-1,3-dioxoisoindolin-5-yl)oxy)acetamide (X11)**

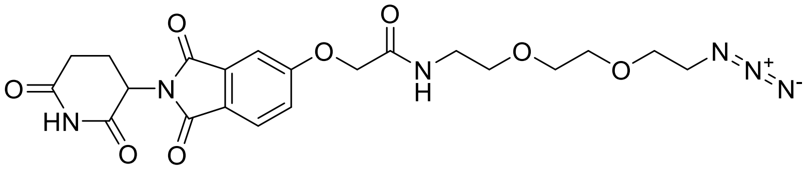

The title compound was prepared according to general procedure B, using **I16** (150 mg, 0.451 mmol) and 1-(2-aminoethoxy)-2-(2-azidoethoxy)ethane (102 mg, 0.587 mmol), 1-methyl-1*H*-imidazole (126 µL, 1.58 mmol) and TCFH (152 mg, 0.542 mmol). The title compound was obtained as a colourless solid (100 mg, 45%). $^{1}$H NMR (500 MHz, Acetone-$d_6$) δ 9.91 (s, 1H), 7.85 (d, *J* = 8.2 Hz, 1H), 7.59 (s, 1H), 7.45 (d, *J* = 2.2 Hz, 1H), 7.43 (dd, *J* = 8.2, 2.3 Hz, 1H), 5.12 (dd, *J* = 12.7, 5.5 Hz, 1H), 4.75 (s, 2H), 3.67 (t, *J* = 4.8 Hz, 2H), 3.63 – 3.58 (m, 4H), 3.56 (t, *J* = 5.7 Hz, 2H), 3.46 (q, *J* = 5.7 Hz, 2H), 3.37 (t, *J* = 4.9 Hz, 2H), 3.08 – 2.90 (m, 1H), 2.81 – 2.73 (m, 2H), 2.29 – 2.17 (m, 1H). $^{13}$C NMR (126 MHz, Acetone-$d_6$) δ 172.60, 170.04, 167.67, 167.61, 167.53, 164.06, 135.33, 126.03, 125.43, 121.56, 110.27, 71.11, 71.01, 70.71, 70.21, 68.58, 51.37, 50.34, 39.55, 31.99, 23.31. ESI-MS: *m/z* = 489.1 ([M+H]$^{+}$).

**Synthesis of *N*-(2-(2-(2-(2-azidoethoxy)ethoxy)ethoxy)ethyl)-2-((2-(2,6-dioxopiperidin-3-yl)-1,3-dioxoisoindolin-5-yl)oxy)acetamide (X12)**

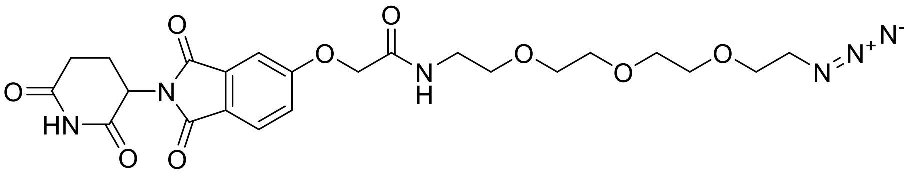

The title compound was prepared according to general procedure B, using **I16** (180 mg, 0.542 mmol) and 1-[2-(2-aminoethoxy)ethoxy]-2-(2-azidoethoxy)ethane (130 mg, 0.596 mmol), 1-methyl-1*H*-imidazole (151 µL, 1.90 mmol) and TCFH (182 mg, 0.650 mmol). The title compound was obtained as a colourless solid (110 mg, 38%). $^{1}$H NMR (400 MHz, Acetone-$d_6$) δ 9.90 (s, 1H), 7.85 (dd, *J* = 8.1, 0.7 Hz, 1H), 7.60 (s, 1H), 7.45 (dd, *J* = 2.4, 0.6 Hz, 1H), 7.43 (dd, *J* = 8.1, 2.4 Hz, 1H), 5.12 (dd, *J* = 12.6, 5.5 Hz, 1H), 4.75 (s,

2H), 3.68 – 3.64 (m, 2H), 3.62 – 3.60 (m, 4H), 3.57 (q, *J* = 1.2 Hz, 4H), 3.56 – 3.53 (m, 2H), 3.45 (q, *J* = 5.7 Hz, 2H), 3.38 (t, *J* = 5.0 Hz, 2H), 3.04 – 2.91 (m, 1H), 2.78 – 2.70 (m, 2H), 2.28 – 2.17 (m, 1H). $^{13}$C NMR (101 MHz, Acetone-$d_6$) δ 171.70, 169.13, 166.78, 166.72, 166.61, 163.17, 134.43, 125.15, 124.53, 120.66, 109.41, 70.37, 70.30, 70.09, 69.80, 69.26, 67.70, 50.49, 49.45, 38.69, 31.09, 22.41. ESI-MS: *m/z* = 533.2 ([M+H]$^+$).

**Synthesis of 5-((3-azidopropyl)amino)-2-(2,6-dioxopiperidin-3-yl)isoindoline-1,3-dione (X13)**

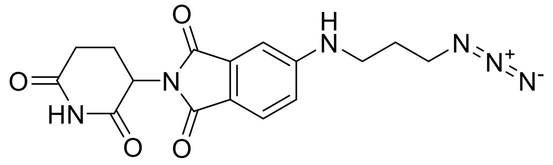

The title compound was prepared according to general procedure C, using 2-(2,6-dioxopiperidin-3-yl)-5-fluoroisoindoline-1,3-dione (200 mg, 0.724 mmol), 3-azidopropan-1-amine (87 mg, 0.87 mmol) and DIPEA (378 µL, 2.17 mmol). The title compound was obtained as a yellow solid (60 mg, 23%). $^1$H NMR (400 MHz, DMSO-$d_6$) δ 11.05 (s, 1H), 7.57 (d, *J* = 8.3 Hz, 1H), 7.15 (t, *J* = 5.5 Hz, 1H), 6.96 (d, *J* = 2.1 Hz, 1H), 6.86 (dd, *J* = 8.4, 2.2 Hz, 1H), 5.03 (dd, *J* = 12.9, 5.4 Hz, 1H), 3.46 (t, *J* = 6.7 Hz, 2H), 3.24 (q, *J* = 6.5 Hz, 2H), 2.96 – 2.80 (m, 1H), 2.65 – 2.52 (m, 2H), 2.08 – 1.95 (m, 1H), 1.82 (quin, *J* = 6.8 Hz, 2H). $^{13}$C NMR (101 MHz, DMSO-$d_6$) δ 172.83, 170.18, 167.67, 167.15, 154.29, 134.23, 125.13, 116.21, 48.64, 48.37, 39.65, 30.99, 27.53, 22.23. ESI-MS: *m/z* = 357.20 ([M+H]$^+$).

**Synthesis of 5-((4-azidobutyl)amino)-2-(2,6-dioxopiperidin-3-yl)isoindoline-1,3-dione (X14)**

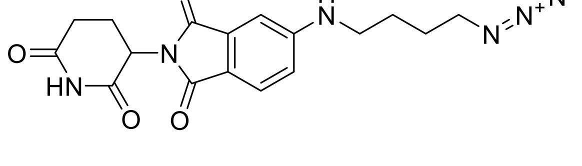

The title compound was prepared according to general procedure C, using 2-(2,6-dioxopiperidin-3-yl)-5-fluoroisoindoline-1,3-dione (200 mg, 0.724 mmol), 4-azidobutan-1-amine (99 mg, 0.87 mmol) and DIPEA (378 µL, 2.17 mmol). The title compound was obtained as a yellow solid (93 mg, 35%). $^1$H NMR (400 MHz, DMSO-$d_6$) δ 11.05 (s, 1H), 7.56 (d, *J* = 8.4 Hz, 1H), 7.13 (t, *J* = 5.5 Hz, 1H), 6.96 (d, *J* = 2.1 Hz, 1H), 6.85 (dd, *J* = 8.4, 2.1 Hz, 1H), 5.03 (dd, *J* = 12.8, 5.5 Hz, 1H), 3.44 – 3.34 (m, 2H), 3.19 (q, *J* = 6.1 Hz, 2H), 2.95 – 2.78 (m, 1H), 2.64 – 2.50 (m, 2H), 2.07 – 1.89 (m, 1H), 1.73 – 1.49 (m, 4H). $^{13}$C NMR (101 MHz, DMSO-$d_6$)δ 172.83, 170.19, 167.71, 167.16, 154.40, 134.24, 125.09, 115.96, 50.39, 48.62, 41.92, 30.99, 25.91, 25.46, 22.23. ESI-MS: *m/z* = 371.20 ([M+H]$^+$).

**Synthesis of 5-((6-azidohexyl)amino)-2-(2,6-dioxopiperidin-3-yl)isoindoline-1,3-dione (X15)**

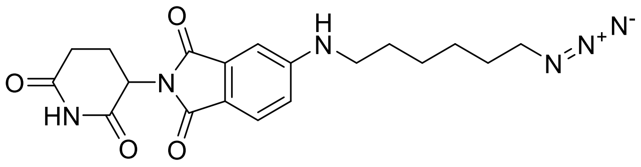

The title compound was prepared according to general procedure C, using 2-(2,6-dioxopiperidin-3-yl)-5-fluoroisoindoline-1,3-dione (200 mg, 0.724 mmol), 6-azidohexan-1-amine (124 mg, 0.867 mmol) and DIPEA (378 µL, 2.17 mmol). The title compound was obtained as a yellow solid (107 mg, 37%). $^1$H NMR (400 MHz, DMSO-$d_6$) δ 11.05 (s, 1H), 7.56 (d, *J* = 8.4 Hz, 1H), 7.09 (t, *J* = 5.4 Hz, 1H), 6.94 (d, *J* = 2.1 Hz, 1H), 6.84 (dd, *J* = 8.4, 2.1 Hz, 1H), 5.03 (dd, *J* = 12.9, 5.4 Hz, 1H), 3.44 – 3.24 (m, 2H), 3.15 (q, *J* = 6.5 Hz, 2H), 2.93 – 2.80 (m, 1H), 2.62 – 2.50 (m, 2H), 2.05 – 1.85 (m, 1H), 1.65 – 1.46 (m, 4H), 1.43 – 1.33 (m, 4H). $^{13}$C NMR (101 MHz, DMSO-$d_6$) δ 172.82, 170.19, 167.71, 167.16, 154.46, 134.22, 125.10, 115.82, 50.58, 48.61, 42.38, 30.99, 28.20, 28.09, 26.04, 25.91, 22.24. ESI-MS: *m/z* = 421.20 ([M+H]$^+$).

**Synthesis of *tert*-butyl (2-((2-(2,6-dioxopiperidin-3-yl)-1,3-dioxoisoindolin-5-yl)amino)-ethyl)carbamate (I17)**

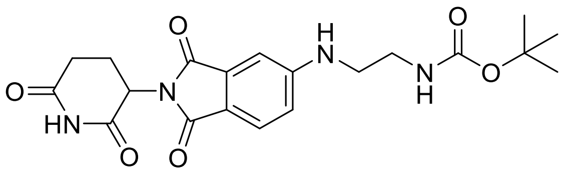

The title compound was prepared according to general procedure C, using 2-(2,6-dioxopiperidin-3-yl)-5-fluoroisoindoline-1,3-dione (2.00 g, 7.24 mmol), *tert*-butyl (2-aminoethyl)carbamate (1.39 g, 8.69 mmol) and DIPEA (3.78 mL, 21.7 mmol). The title compound was obtained as a yellow solid (293 mg, 10%). $^{1}$H NMR (400 MHz, DMSO-$d_6$) δ 11.05 (s, 1H), 7.56 (d, $J$ = 8.4 Hz, 1H), 7.13 (t, $J$ = 5.9 Hz, 1H), 6.99 – 6.96 (m, 1H), 6.91 (t, $J$ = 4.7 Hz, 1H), 6.86 (dd, $J$ = 8.4, 2.2 Hz, 1H), 5.03 (dd, $J$ = 12.9, 5.4 Hz, 1H), 3.22 (quin, $J$ = 6.6 Hz, 2H), 3.15 – 3.09 (m, 2H), 2.96 – 2.81 (m, 1H), 2.60 – 2.51 (m, 2H), 2.04 – 1.94 (m, 1H), 1.37 (s, 9H). ESI-MS: $m/z$ = 439.10 ([M+Na]$^+$).

**Synthesis of 5-((2-aminoethyl)amino)-2-(2,6-dioxopiperidin-3-yl)isoindoline-1,3-dione (I18)**

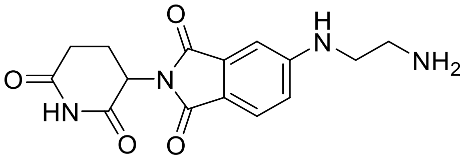

The title compound was prepared according to general procedure D, using **I17** (293 mg, 0.704 mmol). The crude product was used without further purification. ESI-MS: $m/z$ = 317.15 ([M+H]$^+$).

**Synthesis of 2-azido-*N*-(2-((2-(2,6-dioxopiperidin-3-yl)-1,3-dioxoisoindolin-5-yl)amino)-ethyl)acetamide (X16)**

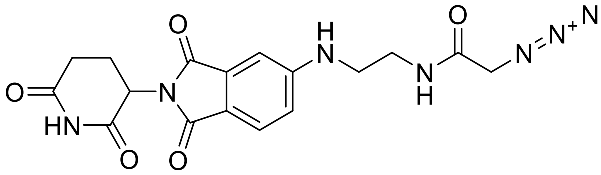

The title compound was prepared according to general procedure A, using **I18** (128 mg, 0.405 mmol), 2-azidoacetic acid (41 mg, 0.41 mmol), HATU (185 mg, 487 mmol) and DIPEA (282 µL, 1.62 mmol). The title compound was obtained as a yellow solid (70 mg, 43%). $^{1}$H NMR (400 MHz, DMSO-$d_6$) δ 11.06 (s, 1H), 8.25 (t, $J$ = 5.1 Hz, 1H), 7.58 (d, $J$ = 8.4 Hz, 1H), 7.17 (t, $J$ = 5.2 Hz, 1H), 7.00 (d, $J$ = 2.1 Hz, 1H), 6.88 (dd, $J$ = 8.4, 2.2 Hz, 1H), 5.03 (dd, $J$ = 12.9, 5.4 Hz, 1H), 3.84 (s, 2H), 3.31 – 3.24 (m, 4H), 2.93 – 2.83 (m, 1H), 2.63 – 2.51

(m, 2H), 2.03 – 1.96 (m, 1H). $^{13}C$ NMR (101 MHz, DMSO-$d_6$) δ 172.83, 170.17, 167.63, 167.14, 154.28, 134.25, 125.12, 116.38, 50.89, 48.65, 41.66, 37.78, 30.99, 22.23. ESI-MS: *m/z* = 400.15 ($[M+H]^+$).

**Synthesis of *tert*-butyl 4-(1-(2-(2,6-dioxopiperidin-3-yl)-1,3-dioxoisoindolin-5-yl)-piperidin-4-yl)piperazine-1-carboxylate (I19)**

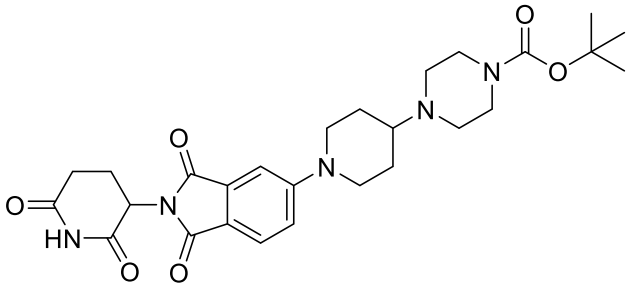

The title compound was prepared according to general procedure C, using 2-(2,6-dioxopiperidin-3-yl)-5-fluoroisoindoline-1,3-dione (500 mg, 1.81 mmol), *tert*-butyl 4-(piperidin-4-yl)piperazine-1-carboxylate (731 mg, 2.72 mmol) and DIPEA (1.26 mL, 7.24 mmol). The title compound was obtained as a yellow solid (551 mg, 58%). $^1H$ NMR (400 MHz, DMSO-$d_6$) δ 11.07 (s, 1H), 7.65 (d, *J* = 8.5 Hz, 1H), 7.31 (d, *J* = 2.3 Hz, 1H), 7.23 (dd, *J* = 8.7, 2.4 Hz, 1H), 5.06 (dd, *J* = 12.9, 5.4 Hz, 1H), 4.06 (d, *J* = 13.3 Hz, 2H), 3.28 (t, *J* = 5.0 Hz, 4H), 2.94 (t, 2H), 2.90 – 2.82 (m, 1H), 2.65 – 2.53 (m, 2H), 2.49 – 2.47 (m, 1H), 2.42 (t, *J* = 5.1 Hz, 4H), 2.04 – 1.97 (m, 1H), 1.82 (d, *J* = 10.8 Hz, 2H), 1.53 – 1.41 (m, 2H), 1.38 (s, 9H). ESI-MS: *m/z* = 526.20 ($[M+H]^+$).

**Synthesis of 2-(2,6-dioxopiperidin-3-yl)-5-(4-(piperazin-1-yl)piperidin-1-yl)isoindoline-1,3-dione (I20)**

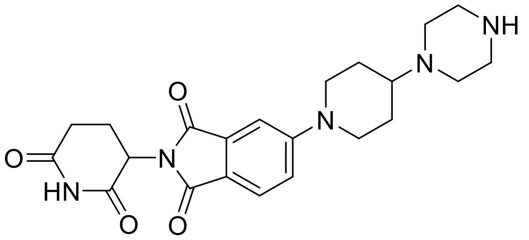

The title compound was prepared according to general procedure D, using **I19** (500 mg, 0.951 mmol). The crude product was used without further purification. ESI-MS: *m/z* = 426.15 ($[M+H]^+$).

**Synthesis of 5-(4-(4-(2-azidoacetyl)piperazin-1-yl)piperidin-1-yl)-2-(2,6-dioxopiperidin-3-yl)isoindoline-1,3-dione (X17)**

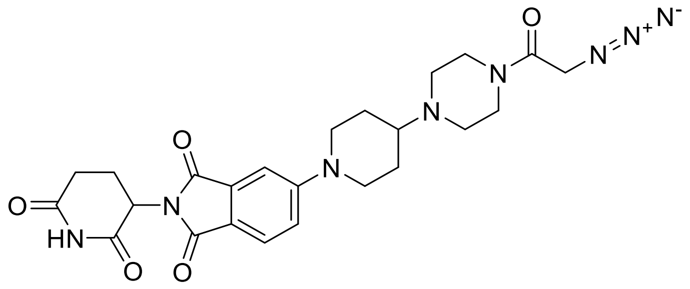

The title compound was prepared according to general procedure A, using **I20** (134 mg, 0.315 mmol), 2-azidoacetic acid (32 mg, 0.32 mmol), HATU (156 mg, 0.409 mmol) and DIPEA (165 µL, 0.945 mmol). The title compound was obtained as a yellow solid (102 mg, 64%). $^1$H NMR (400 MHz, DMSO-$d_6$) δ 11.08 (s, 1H), 7.65 (d, *J* = 8.5 Hz, 1H), 7.32 (d, *J* = 2.3 Hz, 1H), 7.24 (dd, *J* = 8.7, 2.3 Hz, 1H), 5.06 (dd, *J* = 13.0, 5.4 Hz, 1H), 4.12 (s, 2H), 4.07 (d, *J* = 13.4 Hz, 2H), 3.44 (t, *J* = 4.9 Hz, 2H), 3.28 (t, *J* = 5.0 Hz, 2H), 2.96 (t, *J* = 12.3 Hz, 2H), 2.91 – 2.81 (m, 1H), 2.64 – 2.52 (m, 3H), 2.48 – 2.46 (m, 4H), 2.04 – 1.99 (m, 1H), 1.81 (d, *J* = 10.4 Hz, 2H), 1.51 – 1.40 (m, 2H). $^{13}$C NMR (101 MHz, DMSO-$d_6$) δ 172.85, 170.15, 167.63, 167.02, 166.98, 165.73, 154.75, 134.05, 125.03, 117.64, 107.83, 60.56, 49.62, 48.74, 48.62, 48.33, 46.64, 44.45, 41.85, 30.99, 27.04, 22.19. ESI-MS: *m/z* = 509.20 ([M+H]$^+$).

**Synthesis of 5-(4-(4-(4-azidobutanoyl)piperazin-1-yl)piperidin-1-yl)-2-(2,6-dioxopiperidin-3-yl)isoindoline-1,3-dione (X18)**

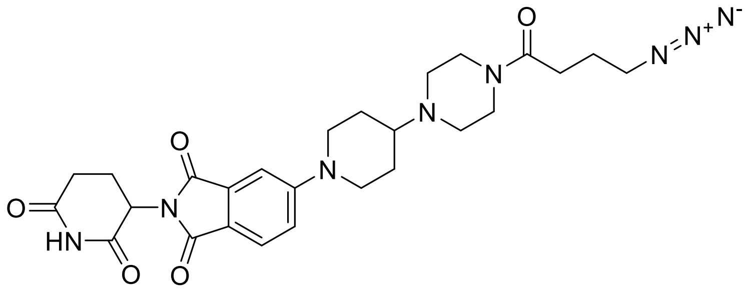

The title compound was prepared according to general procedure A, using **I20** (134 mg, 0.315 mmol), 4-azidobutanoic acid (37 mg, 0.32 mmol), HATU (156 mg, 0.409 mmol) and DIPEA (165 µL, 0.945 mmol). The title compound was obtained as a yellow solid (82 mg, 50%). $^1$H NMR (400 MHz, DMSO-$d_6$) δ 11.08 (s, 1H), 7.65 (d, *J* = 8.5 Hz, 1H), 7.32 (d, *J* = 2.3

Hz, 1H), 7.24 (dd, $J$ = 8.7, 2.3 Hz, 1H), 5.06 (dd, $J$ = 12.9, 5.4 Hz, 1H), 4.07 (d, $J$ = 12.9 Hz, 2H), 3.45 – 3.38 (m, 4H), 3.36 (s, 1H), 3.33 – 3.32 (m, 1H), 2.96 (t, $J$ = 12.6 Hz, 2H), 2.90 – 2.81 (m, 1H), 2.63 – 2.52 (m, 3H), 2.49 – 2.45 (m, 2H), 2.42 (t, $J$ = 5.0 Hz, 2H), 2.36 (t, $J$ = 7.3 Hz, 2H), 2.04 – 1.97 (m, 1H), 1.82 (d, $J$ = 12.5 Hz, 2H), 1.73 (quin, $J$ = 7.1 Hz, 2H), 1.55 – 1.38 (m, 2H). $^{13}$C NMR (101 MHz, DMSO-$d_6$) δ 172.82, 170.12, 169.56, 167.61, 166.97, 154.75, 134.04, 125.01, 117.69, 117.63, 107.81, 60.55, 50.28, 48.94, 48.73, 48.50, 46.62, 45.17, 41.40, 30.98, 29.06, 27.07, 24.09, 22.18. ESI-MS: $m/z$ = 537.20 ([M+H]$^+$).

**Synthesis of 1-(3-hydroxy-2-methylphenyl)dihydropyrimidine-2,4(1*H*,3*H*)-dione (I21)**

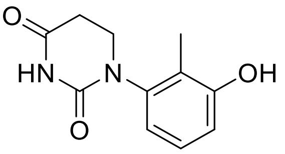


3-amino-2-methylphenol (4.00 g, 32.5 mmol) and acrylic acid (3.34 mL, 48.7 mmol) were dissolved in dry toluene (20 mL) and stirred at 110 °C for 4 h. The solvent was removed under reduced pressure and the crude reaction mixture diluted with acetic acid (25 mL). Urea (5.85 g, 97.4 mmol) was added and stirred at 100 °C for 18 h. The reaction mixture was cooled in an ice bath (0 °C) and diluted with water, filtered and the precipitate was washed with water (20 mL) and *n*-hexane (20 mL). The precipitate was dried under reduced pressure to give the title compound as a colourless solid (6.61 g, 92%). $^1$H NMR (400 MHz, DMSO-$d_6$) δ 10.27 (s, 1H), 9.48 (s, 1H), 7.01 (t, $J$ = 7.8 Hz, 1H), 6.77 (dd, $J$ = 8.1, 1.2 Hz, 1H), 6.70 (dd, $J$ = 7.9, 1.2 Hz, 1H), 3.80 – 3.63 (m, 1H), 3.53 – 3.40 (m, 1H), 2.87 – 2.58 (m, 2H), 1.96 (s, 3H). ESI-MS: $m/z$ = 221.05 ([M+H]$^+$).

**Synthesis of *tert*-butyl 2-(3-(2,4-dioxotetrahydropyrimidin-1(2*H*)-yl)-2-methylphenoxy)-acetate (I22)**

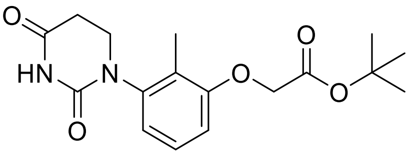

**I21** (2.00 g, 9.08 mmol) and potassium carbonate (3.77 g, 27.2 mmol) were dissolved in dry DMF (20 mL). *Tert*-butyl bromoacetate (1.41 ml, 9.54 mmol) was added dropwise and the reaction mixture was stirred at rt for 2 h. The reaction mixture was diluted with water, filtered and the precipitate was washed with water (10 mL) and *n*-hexane (10 mL). The precipitate was dried under reduced pressure to give the title compound (2.49 g, 82%). $^{1}$H NMR (400 MHz, DMSO-$d_6$) δ 10.31 (s, 1H), 7.17 (t, $J$ = 8.2 Hz, 1H), 6.90 (d, $J$ = 7.9 Hz, 1H), 6.80 (d, $J$ = 8.3 Hz, 1H), 4.69 (s, 2H), 3.84 – 3.66 (m, 1H), 3.54 – 3.40 (m, 1H), 2.86 – 2.60 (m, 2H), 2.05 (s, 3H), 1.43 (s, 9H). ESI-MS: $m/z$ = 357.05 ([M+Na]$^+$).

**Synthesis of 2-(3-(2,4-dioxotetrahydropyrimidin-1(2*H*)-yl)-2-methylphenoxy)acetic acid (I23)**

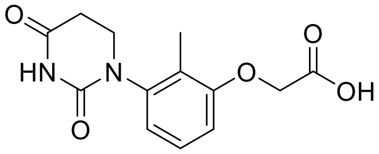


**I22** (1.00 g, 2.99 mmol) was dissolved in dry DCM (10 mL). The flask was cooled to 0 °C in an ice bath and trifluoroacetic acid (TFA) in an excess was added dropwise. The solution was stirred at rt for two h. The solvent was removed under reduced pressure and the crude product used without further purification. ESI-MS: $m/z$ = 279.95 ([M+H]$^+$).

**Synthesis of *N*-(2-(2-(2-azidoethoxy)ethoxy)ethyl)-2-(3-(2,4-dioxotetrahydropyrimidin-1(2*H*)-yl)-2-methylphenoxy)acetamide (X19)**

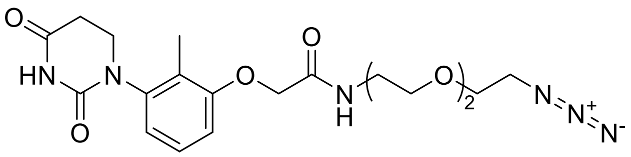


The title compound was prepared according to general procedure A, using **I23** (125 mg, 0.449 mmol), 2-(2-(2-azidoethoxy)ethoxy)ethan-1-amine (78 mg, 0.45 mmol), HATU (256 mg, 0.674 mmol) and DIPEA (234 µL, 1.35 mmol). The title compound was obtained as a colourless solid (122 mg, 62%). $^{1}$H NMR (400 MHz, DMSO-$d_6$) δ 10.33 (s, 1H), 7.95 (t,

*J* = 5.7 Hz, 1H), 7.17 (t, *J* = 8.1 Hz, 1H), 6.91 (d, *J* = 7.6 Hz, 1H), 6.84 (d, *J* = 8.1 Hz, 1H), 4.51 (s, 2H), 3.80 – 3.67 (m, 1H), 3.61 – 3.57 (m, 2H), 3.57 – 3.51 (m, 6H), 3.51 – 3.45 (m, 3H), 3.42 – 3.36 (m, 2H), 2.82 – 2.63 (m, 2H), 2.08 (s, 3H). $^{13}$C NMR (101 MHz, DMSO-*d*$_6$) δ 170.74, 167.70, 156.37, 151.76, 141.79, 126.66, 124.51, 119.92, 110.78, 69.65, 69.60, 69.27, 68.87, 67.58, 49.98, 44.68, 38.27, 31.07, 10.86. ESI-MS: *m/z* = 435.20 ([M+H]$^+$).

**Synthesis of *N*-(20-azido-3,6,9,12,15,18-hexaoxaicosyl)-2-(3-(2,4-dioxotetrahydro-pyrimidin-1(2*H*)-yl)-2-methylphenoxy)acetamide (X20)**

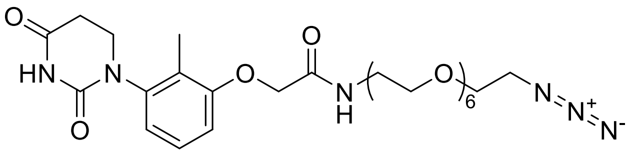

The title compound was prepared according to general procedure A, using **I23** (125 mg, 0.449 mmol), 20-azido-3,6,9,12,15,18-hexaoxaicosan-1-amine (157 mg, 0.449 mmol), HATU (256 mg, 0.674 mmol) and DIPEA (234 µL, 1.35 mmol). The title compound was obtained as a colourless solid (163 mg, 59%). $^1$H NMR (400 MHz, DMSO-*d*$_6$) δ 10.33 (s, 1H), 7.96 (t, *J* = 5.7 Hz, 1H), 7.18 (t, *J* = 8.1 Hz, 1H), 6.91 (d, *J* = 7.9 Hz, 1H), 6.84 (d, *J* = 8.3 Hz, 1H), 4.52 (s, 2H), 3.81 – 3.70 (m, 1H), 3.59 (t, *J* = 4.9 Hz, 2H), 3.56 – 3.44 (m, 25H), 3.38 (t, *J* = 4.9 Hz, 2H), 2.84 – 2.61 (m, 2H), 2.08 (s, 3H). $^{13}$C NMR (101 MHz, DMSO-*d*$_6$) δ 170.73, 167.69, 156.38, 151.75, 141.79, 126.66, 124.51, 119.90, 110.78, 69.83, 69.80, 69.70, 69.61, 69.26, 68.84, 67.58, 50.00, 44.69, 39.37, 38.29, 31.08, 10.86. ESI-MS: *m/z* = 633.25 ([M+Na]$^+$).

**Synthesis of *tert*-butyl 4-(2-(3-(2,4-dioxotetrahydropyrimidin-1(2*H*)-yl)-2-methyl-phenoxy)acetyl)piperazine-1-carboxylate (I24)**

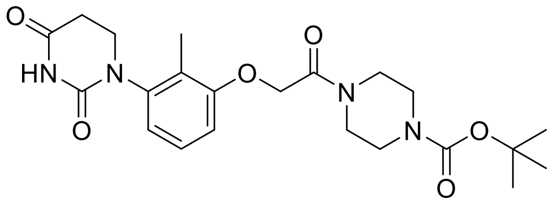

The title compound was prepared according to general procedure A, using **I23** (487 mg, 1.75 mmol), *tert*-butyl piperazine-1-carboxylate (326 mg, 1.75 mmol), HATU (865 mg, 2.27 mmol) and DIPEA (915 µL, 3.42 mmol). The title compound was obtained as a colourless solid (721 mg, 92%). $^{1}$H NMR (400 MHz, DMSO-$d_6$) δ 10.32 (s, 1H), 7.16 (t, $J$ = 8.1 Hz, 1H), 6.88 (d, $J$ = 7.9 Hz, 1H), 6.85 (d, $J$ = 8.3 Hz, 1H), 4.87 (s, 2H), 3.83 – 3.68 (m, 1H), 3.50 (t, $J$ = 6.1 Hz, 1H), 3.48 – 3.44 (m, 4H), 3.38 (s, 2H), 3.30 (s, 2H), 2.84 – 2.61 (m, 2H), 2.05 (s, 3H), 1.41 (s, 9H). ESI-MS: *m/z* = 469.20 ([M+Na]$^+$).

**Synthesis of 1-(2-methyl-3-(2-oxo-2-(piperazin-1-yl)ethoxy)phenyl)dihydropyrimidine-2,4(1H,3H)-dione (I25)**

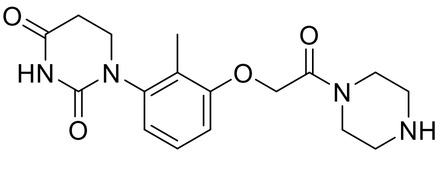


The title compound was prepared according to general procedure D, using **I24** (710 mg, 1.59 mmol). The crude product was used without further purification. ESI-MS: *m/z* = 347.15 ([M+H]$^+$).

**Synthesis of 1-(3-(2-(4-(2-azidoacetyl)piperazin-1-yl)-2-oxoethoxy)-2-methylphenyl)-dihydropyrimidine-2,4(1*H*,3*H*)-dione (X21)**

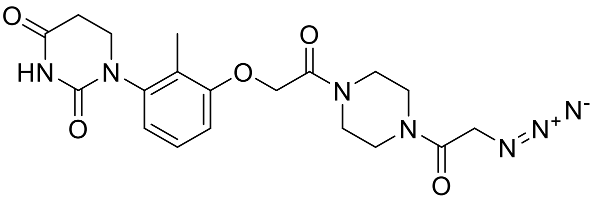


The title compound was prepared according to general procedure A, using **I25** (138 mg, 0.398 mmol), 2-azidoacetic acid (40 mg, 0.39 mmol), HATU (197 mg, 0.518 mmol) and DIPEA (207 µL, 1.19 mmol). The title compound was obtained as a colourless solid (132 mg, 77%). $^{1}$H NMR (400 MHz, DMSO-$d_6$) δ 10.32 (s, 1H), 7.17 (t, $J$ = 8.1 Hz, 1H), 6.87 (t, $J$ = 8.4 Hz, 2H), 4.90 (s, 2H), 4.18 (s, 2H), 3.82 – 3.66 (m, 1H), 3.55 – 3.47 (m, 7H), 3.40 (s, 2H), 2.86 – 2.61 (m, 2H), 2.05 (s, 3H). $^{13}$C NMR (101 MHz, DMSO-$d_6$) δ 170.76, 166.20,

166.15, 156.57, 151.77, 141.72, 126.52, 124.18, 119.56, 110.72, 66.46, 49.74, 44.70, 44.02, 43.76, 41.09, 31.07, 10.76. ESI-MS: *m/z* = 430.15 ([M+H]$^+$).

**Synthesis of 1-(3-(2-(4-(3-azidopropanoyl)piperazin-1-yl)-2-oxoethoxy)-2-methylphenyl)-dihydropyrimidine-2,4(1*H*,3*H*)-dione (X22)**

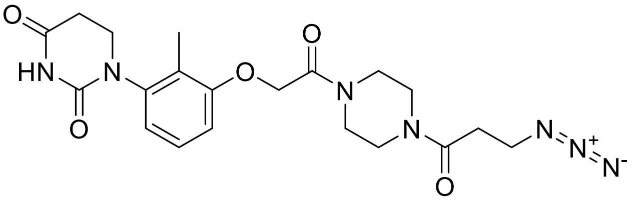

The title compound was prepared according to general procedure A, using **I25** (138 mg, 0.398 mmol), 3-azidopropanoic acid (46 mg, 0.39 mmol), HATU (197 mg, 0.518 mmol) and DIPEA (207 µL, 1.19 mmol). The title compound was obtained as a colourless solid (118 mg, 67%). $^1$H NMR (400 MHz, DMSO-*d*$_6$) δ 10.32 (s, 1H), 7.17 (t, *J* = 8.1 Hz, 1H), 6.88 (t, *J* = 8.3 Hz, 2H), 4.90 (s, 2H), 3.84 – 3.67 (m, 1H), 3.57 – 3.43 (m, 11H), 2.85 – 2.73 (m, 1H), 2.71 – 2.59 (m, 3H), 2.05 (s, 3H). $^{13}$C NMR (101 MHz, DMSO-*d*$_6$) δ 170.75, 168.70, 166.13, 156.57, 151.76, 141.70, 126.51, 124.18, 119.54, 110.72, 66.42, 46.68, 44.69, 44.37, 43.92, 41.16, 40.81, 31.75, 31.08, 10.76. ESI-MS: *m/z* = 444.25 ([M+H]$^+$).

**Synthesis of 1-(3-(2-(4-(4-azidobutanoyl)piperazin-1-yl)-2-oxoethoxy)-2-methylphenyl)-odihydropyrimidine-2,4(1*H*,3*H*)-dione (X23)**

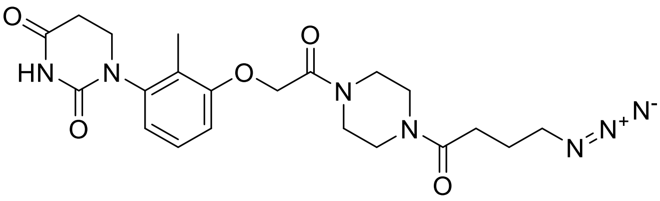

The title compound was prepared according to general procedure A, using **I25** (138 mg, 0.398 mmol), 4-azidobutanoic acid (51 mg, 0.39 mmol), HATU (197 mg, 0.518 mmol) and DIPEA (207 µL, 1.19 mmol). The title compound was obtained as a colourless solid (130 mg, 71%). $^1$H NMR (400 MHz, DMSO-*d*$_6$) δ 10.32 (s, 1H), 7.17 (t, *J* = 8.1 Hz, 1H), 6.87 (t, *J* = 8.8 Hz, 2H), 4.89 (s, 2H), 3.81 – 3.70 (m, 1H), 3.66 – 3.43 (m, 9H), 3.36 (t, *J* = 7.1 Hz,

2H), 2.87 – 2.58 (m, 2H), 2.42 (t, *J* = 7.3 Hz, 2H), 2.05 (s, 3H), 1.77 (quin, *J* = 7.1 Hz, 2H). $^{13}C$ NMR (101 MHz, DMSO-$d_6$) δ 170.75, 170.11, 166.12, 156.58, 151.77, 141.71, 126.51, 124.19, 119.55, 110.72, 66.45, 50.29, 48.61, 44.77, 44.70, 43.97, 41.39, 41.14, 31.08, 29.16, 24.02, 10.75. ESI-MS: *m/z* = 458.20 ([M+H]$^+$).

**Synthesis of 1-(3-(2-(4-(4-azidobenzoyl)piperazin-1-yl)-2-oxoethoxy)-2-methylphenyl)-dihydropyrimidine-2,4(1*H*,3*H*)-dione (X24)**

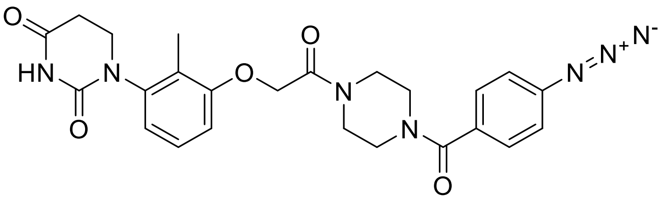

The title compound was prepared according to general procedure A, using **I25** (138 mg, 0.398 mmol), 4-azidobenzoic acid (65 mg, 0.39 mmol), HATU (197 mg, 0.518 mmol) and DIPEA (207 µL, 1.19 mmol). The title compound was obtained as a colourless solid (131 mg, 67%). $^1H$ NMR (400 MHz, DMSO-$d_6$) δ 10.32 (s, 1H), 7.49 (d, *J* = 8.0 Hz, 2H), 7.22 – 7.12 (m, 3H), 6.87 (dd, *J* = 10.8, 8.0 Hz, 2H), 4.90 (s, 2H), 3.81 – 3.69 (m, 1H), 3.60 – 3.37 (m, 9H), 2.85 – 2.58 (m, 2H), 2.05 (s, 3H). $^{13}C$ NMR (101 MHz, DMSO-$d_6$) δ 170.70, 168.54, 166.10, 156.54, 151.73, 141.69, 140.83, 132.13, 129.12, 126.47, 124.18, 119.54, 119.09, 110.68, 66.47, 44.66, 31.06, 10.74. ESI-MS: *m/z* = 492.15 ([M+H]$^+$).

**Synthesis of 1-(3-iodo-2-methylphenyl)dihydropyrimidine-2,4(1*H*,3*H*)-dione (I26)**

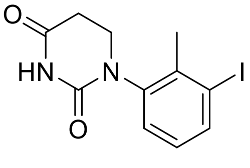

3-iodo-2-methylaniline (10.0 g, 42.9 mmol) and acrylic acid (4.42 mL, 64.4 mmol) were dissolved in dry toluene (30 mL) and stirred at 110 °C for 4 h. The solvent was removed under reduced pressure and the crude reaction mixture diluted with acetic acid (35 mL). Urea (7.73 g, 0.129 mol) was added and stirred at 100 °C for 18 h. The reaction mixture was cooled in an ice bath (0 °C) and diluted with water, filtered and the precipitate was washed with water (20 mL) and *n*-hexane (20 mL). The precipitate was dried under reduced pressure to give the title compound as a light-brown solid (11.9 g, 84%). $^{1}$H NMR (400 MHz, DMSO-$d_6$) δ 10.39 (s, 1H), 7.82 (dd, *J* = 7.9, 1.2 Hz, 1H), 7.33 (dd, *J* = 7.9, 1.2 Hz, 1H), 7.03 (t, *J* = 8.0 Hz, 1H), 3.84 – 3.66 (m, 1H), 3.57 – 3.48 (m, 1H), 2.85 – 2.61 (m, 2H), 2.28 (s, 3H). ESI-MS: *m/z* = 331.85 ([M+H]$^+$).

**Synthesis of 1-(3-azido-2-methylphenyl)dihydropyrimidine-2,4(1*H*,3*H*)-dione (X25)**

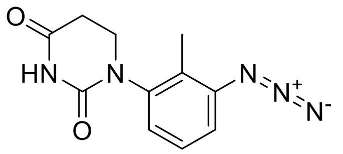


**I26** (200 mg, 0.606 mmol), sodium azide (79 mg, 1.2 mmol), copper iodide (23 mg, 0.12 mmol), *L*-proline (28 mg, 0.24 mmol) and potassium carbonate (236 mg, 1.82 mmol) were dissolved in dry DMSO (2 mL) and stirred at 80 °C for 18 h. The solvent was removed under reduced pressure and the crude product was purified by flash chromatography using acetonitrile/water as an eluent. The title compound was obtained as a colourless solid (48 mg, 32%). $^{1}$H NMR (400 MHz, DMSO-$d_6$) δ 10.38 (s, 1H), 7.34 (t, *J* = 7.9 Hz, 1H), 7.24 (dd, *J* = 8.1, 1.2 Hz, 1H), 7.13 (dd, *J* = 7.8, 1.2 Hz, 1H), 3.89 – 3.69 (m, 1H), 3.54 – 3.41 (m, 1H), 2.87 – 2.73 (m, 1H), 2.71 – 2.60 (m, 1H), 2.03 (s, 3H). $^{13}$C NMR (101 MHz, DMSO-$d_6$) δ 170.74, 151.78, 142.35, 138.70, 127.51, 127.32, 123.68, 117.48, 44.59, 31.06, 12.09. ESI-MS: *m/z* = 246.10 ([M+H]$^+$).

**Synthesis of *tert*-butyl (*E*)-4-(3-(2,4-dioxotetrahydropyrimidin-1(2*H*)-yl)-2-methylstyryl)-piperidine-1-carboxylate (I27)**

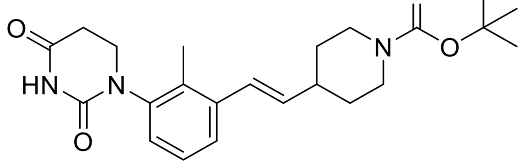


**I26** (500 mg, 1.52 mmol), *tert*-butyl 4-vinylpiperidine-1-carboxylate (640 mg, 3.03 mmol), potassium acetate (297 mg, 3.03 mmol) and palladium(II) acetate (17 mg, 0.076 mmol)

were dissolved in dry DMF (5 mL) and stirred at 100 °C for 18 h. The solvent was removed under reduced pressure and the crude product was purified by flash chromatography using acetonitrile/water as an eluent. The title compound was obtained as a colourless solid (590 mg, 94%). $^{1}$H NMR (400 MHz, DMSO-$d_6$) δ 10.32 (s, 1H), 7.39 (d, *J* = 7.4 Hz, 1H), 7.24 – 7.08 (m, 3H), 6.63 (d, *J* = 15.8 Hz, 1H), 6.09 (dd, *J* = 15.8, 6.8 Hz, 1H), 3.97 (d, *J* = 12.9 Hz, 2H), 3.81 – 3.67 (m, 1H), 3.55 – 3.44 (m, 1H), 2.86 – 2.61 (m, 4H), 2.41 – 2.30 (m, 1H), 2.12 (s, 3H), 2.11 – 2.04 (m, 1H), 1.74 (d, *J* = 12.5 Hz, 2H), 1.41 (s, 9H). ESI-MS: *m/z* = 436.25 ([M+Na]$^{+}$).

**Synthesis of (*E*)-1-(2-methyl-3-(2-(piperidin-4-yl)vinyl)phenyl)dihydropyrimidine-2,4(1*H*,3*H*)-dione (I28)**

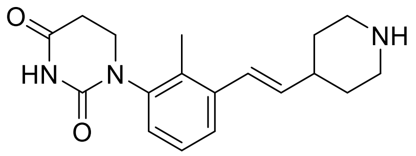


The title compound was prepared according to general procedure D, using **I27** (310 mg, 0.939 mmol). The crude product was used without further purification. ESI-MS: *m/z* = 314.20 ([M+H]$^{+}$).

**Synthesis of (*E*)-1-(3-(2-(1-(2-azidoacetyl)piperidin-4-yl)vinyl)-2-methylphenyl)dihydro-pyrimidine-2,4(1*H*,3*H*)-dione (X26)**

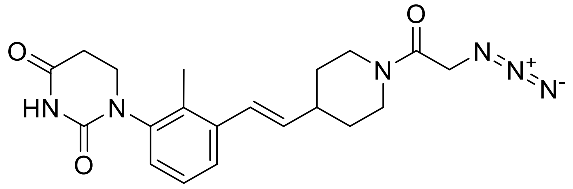


The title compound was prepared according to general procedure A, using **I28** (118 mg, 0.377 mmol), 2-azidoacetic acid (38 mg, 0.38 mmol), HATU (186 mg, 0.489 mmol) and DIPEA (197 µL, 1.13 mmol). The title compound was obtained as a colourless solid (106 mg, 71%). $^{1}$H NMR (400 MHz, DMSO-$d_6$) δ 10.33 (s, 1H), 7.39 (d, *J* = 7.4 Hz, 1H), 7.28 – 7.06 (m, 3H), 6.64 (d, *J* = 15.8 Hz, 1H), 6.10 (dd, *J* = 15.9, 6.7 Hz, 1H), 4.35 (q, *J* = 16.2 Hz,

1H), 4.25 – 4.01 (m, 3H), 3.85 – 3.57 (m, 2H), 3.58 – 3.37 (m, 1H), 3.17 – 2.89 (m, 1H), 2.90 – 2.54 (m, 4H), 2.47 – 2.36 (m, 1H), 1.89 – 1.54 (m, 2H), 1.55 – 1.23 (m, 2H).

$^{13}$C NMR (101 MHz, DMSO-$d_6$) δ 170.74, 165.57, 151.90, 142.45, 141.23, 137.75, 136.79, 132.74, 126.41, 125.99, 124.77, 113.24, 49.74, 44.71, 43.97, 41.37, 40.19, 39.52, 31.16, 31.05, 13.77. ESI-MS: *m/z* = 397.15 ([M+H]$^+$).

**Synthesis of (*E*)-1-(3-(2-(1-(4-azidobenzoyl)piperidin-4-yl)vinyl)-2-methylphenyl)-dihydropyrimidine-2,4(1*H*,3*H*)-dione (X27)**

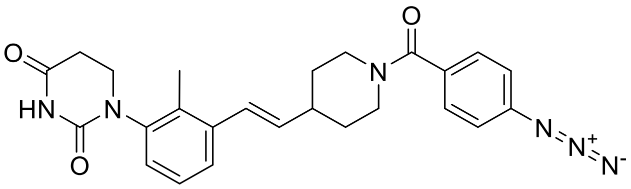


The title compound was prepared according to general procedure A, using **I28** (118 mg, 0.377 mmol), 4-azidobenzoic acid (61 mg, 0.38 mmol), HATU (186 mg, 0.489 mmol) and DIPEA (197 µL, 1.13 mmol). The title compound was obtained as a colourless solid (88 mg, 51%). $^1$H NMR (400 MHz, DMSO-$d_6$) δ 10.34 (s, 1H), 7.41 (dd, *J* = 7.7, 1.6 Hz, 1H), 7.28 – 7.10 (m, 3H), 6.66 (d, *J* = 15.8 Hz, 1H), 6.12 (dd, *J* = 15.8, 6.7 Hz, 1H), 4.49 – 4.32 (m, 1H), 4.23 – 4.04 (m, 3H), 3.84 – 3.66 (m, 2H), 3.56 – 3.42 (m, 1H), 3.06 (t, *J* = 12.5 Hz, 1H), 2.84 – 2.64 (m, 3H), 2.14 (s, 3H), 1.87 – 1.60 (m, 3H), 1.49 – 1.20 (m, 2H). $^{13}$C NMR (101 MHz, DMSO-$d_6$) δ 170.74, 165.56, 151.89, 142.45, 141.23, 137.75, 136.78, 132.73, 126.41, 126.03, 125.99, 124.76, 113.24, 49.73, 44.70, 43.96, 41.36, 31.66, 31.14, 31.04, 13.77. ESI-MS: *m/z* = 459.20 ([M+H]$^+$).

**Synthesis of *tert*-butyl 4-(3-(2,4-dioxotetrahydropyrimidin-1(2*H*)-yl)-2-methylphenethyl)-piperidine-1-carboxylate (I29)**

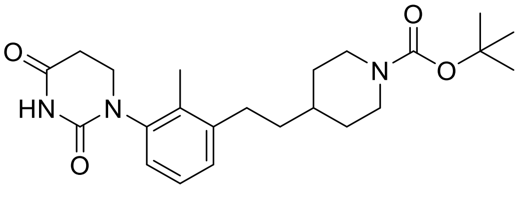

**I27** (280 mg, 0.677 mmol) was dissolved in dry DMF (10 mL). Pd/C (5%, 72 mg, 0.034 mmol) was added and the argon atmosphere was replaced with a hydrogen atmosphere. The reaction mixture was stirred at rt for 2 h. The reaction mixture was filtered through a pad of celite and the filter cake was rinsed with methanol (20 mL). The solvents were removed under reduced pressure to give the title compound as a colourless solid (202 mg, 69%). $^1$H NMR (400 MHz, DMSO-$d_6$) δ 10.31 (s, 1H), 7.22 – 7.00 (m, 3H), 3.94 (d, *J* = 12.8 Hz, 2H), 3.78 – 3.63 (m, 1H), 3.55 – 3.37 (m, 1H), 2.82 – 2.58 (m, 6H), 2.10 (s, 3H), 1.72 (d, *J* = 12.9 Hz, 2H), 1.55 – 1.41 (m, 5H), 1.39 (s, 9H). ESI-MS: *m/z* = 438.25 ([M+Na]$^+$).

**Synthesis of 1-(2-methyl-3-(2-(piperidin-4-yl)ethyl)phenyl)dihydropyrimidine-2,4(1*H*,3*H*)-dione (I30)**

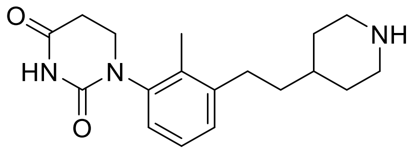


The title compound was prepared according to general procedure D, using **I29** (119 mg, 0.286 mmol). The crude product was used without further purification. ESI-MS: *m/z* = 338.25 ([M+Na]$^+$).

**Synthesis of 1-(3-(2-(1-(2-azidoacetyl)piperidin-4-yl)ethyl)-2-methylphenyl)dihydro-pyrimidine-2,4(1*H*,3*H*)-dione (X28)**

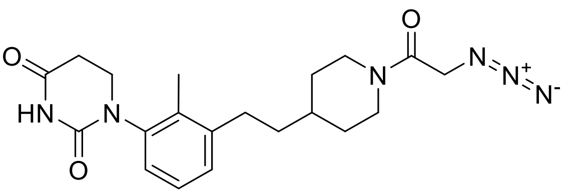


The title compound was prepared according to general procedure A, using **I30** (90 mg, 0.286 mmol), 2-azidoacetic acid (29 mg, 0.29 mmol), HATU (141 mg, 0.372 mmol) and DIPEA (149 µL, 0.859 mmol). The title compound was obtained as a colourless solid

(99 mg, 87%). $^1$H NMR (400 MHz, DMSO-$d_6$) δ 10.31 (s, 1H), 7.43 – 6.71 (m, 3H), 4.35 (d, $J$ = 13.0 Hz, 1H), 4.13 (d, $J$ = 6.4 Hz, 2H), 3.76 – 3.68 (m, 1H), 3.63 (d, $J$ = 13.4 Hz, 1H), 3.52 – 3.43 (m, 1H), 2.95 (t, $J$ = 12.6 Hz, 1H), 2.82 – 2.72 (m, 1H), 2.71 – 2.56 (m, 4H), 2.10 (s, 3H), 1.78 (s, 2H), 1.62 – 1.49 (m, 0H), 1.53 – 1.39 (m, 2H), 1.26 (t, $J$ = 5.7 Hz, 1H), 1.23 – 0.96 (m, 2H). $^{13}$C NMR (101 MHz, DMSO-$d_6$) δ 170.77, 165.47, 151.86, 141.87, 141.19, 133.48, 127.93, 126.23, 124.88, 49.73, 44.77, 44.28, 41.72, 36.75, 35.28, 31.96, 31.37, 31.07, 30.34, 13.23. ESI-MS: $m/z$ = 399.20 ([M+H]$^+$).

**Synthesis of *N*-(2,6-dioxopiperidin-3-yl)-2-methoxy-4-nitrobenzamide (I31)**

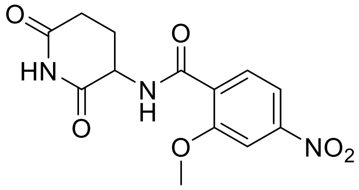


2-methoxy-4-nitrobenzoic acid (5.14 g, 26.1 mmol), 3-aminopiperidine-2,6-dione hydrochloride (8.58 g, 52.1 mmol), 1-ethyl-3-(3-dimethylaminopropyl)carbodiimide hydrochloride (5.50 g, 28.7 mmol), 1-hydroxybenzotriazole hydrate (4.39 g, 28.7 mmol) were dissolved in dry DMF (40 mL). DIPEA (15.2 mL, 117 mmol) was added and the reaction mixture was stirred at rt for 18 h. The reaction mixture was cooled in an ice bath (0 °C) and diluted with water, filtered and the precipitate was washed with water (50 mL). The precipitate was dried under reduced pressure to give the title compound as a colourless solid (6.72 g, 84%). $^1$H NMR (400 MHz, DMSO-$d_6$) δ 10.90 (s, 1H), 8.75 (d, $J$ = 7.8 Hz, 1H), 7.97 – 7.88 (m, 3H), 4.86 – 4.65 (m, 1H), 4.01 (s, 3H), 2.86 – 2.70 (m, 1H), 2.58 – 2.51 (m, 1H), 2.19 – 1.92 (m, 2H). ESI-MS: $m/z$ = 308.05 ([M+H]$^+$).

**Synthesis of 4-amino-*N*-(2,6-dioxopiperidin-3-yl)-2-methoxybenzamide (I32)**

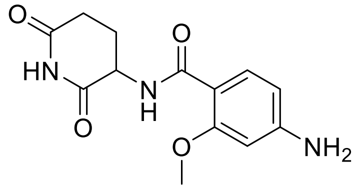

**I31** (1.29 mg, 4.20 mmol), iron powder (1.17 g, 20.9 mmol) and ammonium chloride (1.12 g, 20.9 mmol) were dissolved in dry methanol (10 mL) and water (5 mL) and the reaction mixture was stirred at 65 °C for 2 h. The reaction mixture was filtered through a pad of celite and the filter cake was rinsed with methanol (15 mL). The solvent was removed under reduced pressure. The crude product was purified by flash chromatography using acetonitrile/water as an eluent to obtain the title compound as a yellow solid (449 mg, 39%). $^1$H NMR (400 MHz, DMSO-$d_6$) δ 10.85 (s, 1H), 8.35 (d, *J* = 6.9 Hz, 1H), 7.66 (d, *J* = 8.5 Hz, 1H), 6.25 (s, 1H), 6.21 (dd, *J* = 8.5, 1.9 Hz, 1H), 5.79 (s, 2H), 4.68 (dt, *J* = 12.3, 6.0 Hz, 1H), 3.83 (s, 3H), 2.81 – 2.68 (m, 1H), 2.54 – 2.42 (m, 1H), 2.18 – 1.97 (m, 2H). ESI-MS: *m/z* = 310.10 ([M+H]$^+$).

**Synthesis of 4-(2-azidoacetamido)-*N*-(2,6-dioxopiperidin-3-yl)-2-methoxybenzamide (X29)**

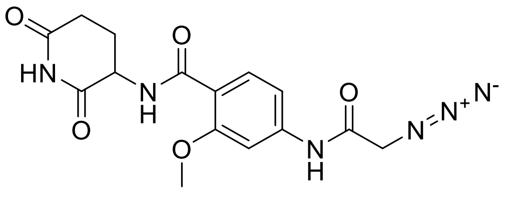


The title compound was prepared according to general procedure A, using **I31** (110 mg, 0.397 mmol), 2-azidoacetic acid (40 mg, 0.40 mmol), HATU (196 mg, 0.517 mmol) and DIPEA (208 µL, 1.19 mmol). The title compound was obtained as a colourless solid (115 mg, 80%). $^1$H NMR (400 MHz, DMSO-$d_6$) δ 10.88 (s, 1H), 10.41 (s, 1H), 8.54 (d, *J* = 7.3 Hz, 1H), 7.87 (d, *J* = 8.5 Hz, 1H), 7.56 (d, *J* = 1.9 Hz, 1H), 7.22 (dd, *J* = 8.6, 1.9 Hz, 1H), 4.74 (q, *J* = 8.5 Hz, 1H), 4.08 (s, 2H), 3.91 (s, 3H), 2.84 – 2.69 (m, 1H), 2.56 – 2.50 (m, 1H), 2.17 – 2.02 (m, 2H). $^{13}$C NMR (101 MHz, DMSO-$d_6$) δ 172.98, 172.44, 166.95, 163.93, 157.94, 142.66, 131.97, 116.35, 111.04, 102.36, 55.90, 51.37, 50.08, 31.01, 24.19. ESI-MS: *m/z* = 361.15 ([M+H]$^+$).

**Synthesis of 4-(3-azidopropanamido)-*N*-(2,6-dioxopiperidin-3-yl)-2-methoxybenzamide (X30)**

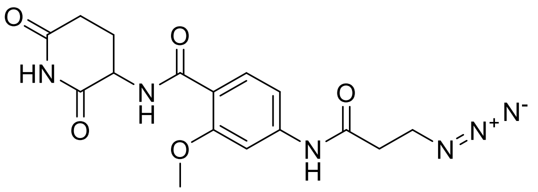

The title compound was prepared according to general procedure A, using **I31** (200 mg, 0.721 mmol), 3-azidopropanoic acid (125 mg, 1.08 mmol), HATU (356 mg, 0.937 mmol) and DIPEA (377 µL, 2.16 mmol). The title compound was obtained as a colourless solid (156 mg, 58%). $^{1}$H NMR (400 MHz, DMSO-$d_6$) δ 10.88 (s, 1H), 10.33 (s, 1H), 8.53 (d, *J* = 7.2 Hz, 1H), 7.86 (d, *J* = 8.5 Hz, 1H), 7.56 (d, *J* = 1.9 Hz, 1H), 7.23 (dd, *J* = 8.6, 1.9 Hz, 1H), 4.81 – 4.69 (m, 1H), 3.90 (s, 3H), 3.63 (t, *J* = 6.3 Hz, 2H), 2.83 – 2.74 (m, 1H), 2.66 (t, *J* = 6.3 Hz, 2H), 2.57 – 2.52 (m, 1H), 2.15 – 2.10 (m, 2H). $^{13}$C NMR (101 MHz, DMSO-$d_6$) δ 172.95, 172.44, 169.26, 163.95, 157.92, 143.23, 131.88, 115.90, 110.85, 102.09, 55.83, 50.07, 46.66, 35.71, 30.99, 24.19. ESI-MS: *m/z* = 375.10 ([M+H]$^+$).

**Synthesis of 4-(4-azidobutanamido)-*N*-(2,6-dioxopiperidin-3-yl)-2-methoxybenzamide (X31)**

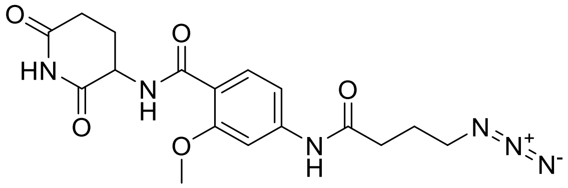

The title compound was prepared according to general procedure A, using **I31** (200 mg, 0.721 mmol), 4-azidobutanoic acid (140 mg, 1.08 mmol), HATU (356 mg, 0.937 mmol) and DIPEA (377 µL, 2.16 mmol). The title compound was obtained as a colourless solid (190 mg, 68%). $^{1}$H NMR (400 MHz, DMSO-$d_6$) δ 10.87 (s, 1H), 10.23 (s, 1H), 8.52 (d, *J* = 7.2 Hz, 1H), 7.85 (d, *J* = 8.5 Hz, 1H), 7.58 (d, *J* = 1.9 Hz, 1H), 7.21 (dd, *J* = 8.6, 1.8 Hz, 1H), 4.79 – 4.63 (m, 1H), 3.89 (s, 3H), 3.40 (t, *J* = 6.8 Hz, 2H), 2.86 – 2.70 (m, 1H), 2.55 – 2.52 (m, 1H), 2.44 (t, *J* = 7.3 Hz, 2H), 2.18 – 2.02 (m, 2H), 1.85 (quin, *J* = 7.0 Hz, 2H). $^{13}$C NMR (101 MHz, DMSO-$d_6$) δ 172.94, 172.45, 170.96, 163.96, 157.91, 143.48, 131.81, 115.65, 110.79, 102.05, 55.81, 50.23, 50.06, 33.34, 30.99, 24.19, 24.07. ESI-MS: *m/z* = 389.10 ([M+H]$^+$).

**Synthesis of *N*-(2,6-dioxopiperidin-3-yl)-4-iodo-2-methoxybenzamide (I32)**

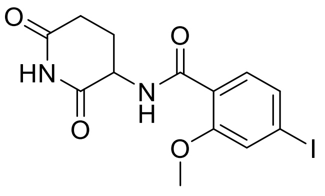

4-iodo-2-methoxybenzoic acid (10.0 g, 35.9 mmol), 3-aminopiperidine-2,6-dione hydrochloride (11.84 g, 71.9 mmol), 1-ethyl-3-(3-dimethylaminopropyl)carbodiimide hydrochloride (7.58 g, 39.6 mmol), 1-hydroxybenzotriazole hydrate (6.06 g, 39.6 mmol) were dissolved in dry DMF (50 mL). DIPEA (28.1 mL, 162 mmol) was added and the reaction mixture was stirred at rt for 18 h. The reaction mixture was cooled in an ice bath (0 °C) and diluted with water, filtered and the precipitate was washed with water (50 mL). The precipitate was dried under reduced pressure to give the title compound as a light grey solid (12.3 g, 89%). $^{1}$H NMR (400 MHz, DMSO-$d_6$) δ 10.87 (s, 1H), 8.56 (d, *J* = 7.5 Hz, 1H), 7.57 (d, *J* = 8.1 Hz, 1H), 7.50 (d, *J* = 1.5 Hz, 1H), 7.45 (dd, *J* = 8.1, 1.5 Hz, 1H), 4.78 – 4.68 (m, 1H), 3.92 (s, 3H), 2.83 – 2.70 (m, 1H), 2.54 (t, *J* = 3.7 Hz, 1H), 2.16 – 2.00 (m, 2H). ESI-MS: *m/z* = 388.95 ([M+H]$^{+}$).

**Synthesis of *tert*-butyl 3-((4-((2,6-dioxopiperidin-3-yl)carbamoyl)-3-methoxyphenyl)amino)azetidine-1-carboxylate (I33)**

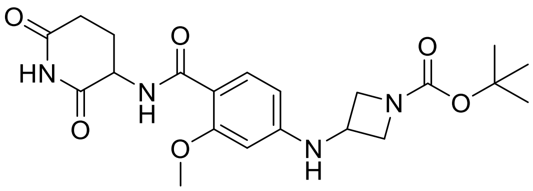

**I32** (1.50 g, 3.86 mmol), *tert*-butyl 3-aminoazetidine-1-carboxylate (1.33 g, 7.73 mmol), copper iodide (147 mg, 0.775 mmol), *L*-proline (178 mg, 1.55 mmol) and potassium carbonate (1.60 g, 11.6 mmol) were dissolved in dry DMSO (10 mL) and stirred at 80 °C for 18 h. The solvent was removed under reduced pressure and the crude product was purified by flash chromatography using acetonitrile/water as an eluent. The title compound was obtained as a colourless solid (830 mg, 50%). $^{1}$H NMR (400 MHz, DMSO-

$d_6$) δ 10.85 (s, 1H), 8.02 (d, *J* = 6.6 Hz, 1H), 7.93 (d, *J* = 8.6 Hz, 1H), 6.18 (dd, *J* = 8.6, 2.1 Hz, 1H), 6.07 (d, *J* = 2.1 Hz, 1H), 4.76 – 4.61 (m, 2H), 4.32 – 4.18 (m, 5H), 3.93 (s, 3H), 3.81 – 3.69 (m, 4H), 1.42 (s, 9H). ESI-MS: *m/z* = 433.15 ([M+H]$^+$).

**Synthesis of 4-(azetidin-3-ylamino)-*N*-(2,6-dioxopiperidin-3-yl)-2-methoxybenzamide (I34)**

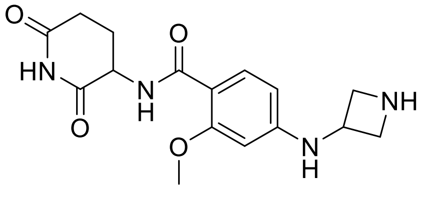

The title compound was prepared according to general procedure D, using **I33** (800 mg, 0.185 mmol). The crude product was used without further purification. ESI-MS: *m/z* = 333.20 ([M+H]$^+$).

**Synthesis of 4-((1-(2-azidoacetyl)azetidin-3-yl)amino)-*N*-(2,6-dioxopiperidin-3-yl)-2-methoxybenzamide (X32)**

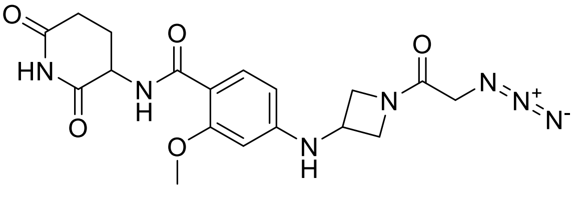

The title compound was prepared according to general procedure A, using **I34** (130 mg, 0.352 mmol), 2-azidoacetic acid (39 mg, 0.39 mmol), HATU (201 mg, 0.582 mmol) and DIPEA (245 µL, 1.41 mmol). The title compound was obtained as a colourless solid (140 mg, 96%). $^1$H NMR (400 MHz, DMSO-$d_6$) δ 10.86 (s, 1H), 8.38 (d, *J* = 6.9 Hz, 1H), 7.74 (d, *J* = 8.4 Hz, 1H), 6.99 (d, *J* = 5.7 Hz, 1H), 6.23 – 6.13 (m, 2H), 4.69 (dt, *J* = 12.5, 6.3 Hz, 1H), 4.48 (t, *J* = 7.6 Hz, 1H), 4.36 – 4.23 (m, 2H), 3.91 (s, 2H), 3.88 (s, 3H), 3.76 (dd, *J* = 9.1, 4.0 Hz, 1H), 2.89 (s, 1H), 2.75 (ddd, *J* = 17.2, 13.3, 5.7 Hz, 1H), 2.54 – 2.52 (m, 1H), 2.21 – 1.93 (m, 2H). $^{13}$C NMR (101 MHz, DMSO-$d_6$) δ 172.98, 172.74, 167.22, 164.46, 159.21,

151.37, 132.86, 109.55, 104.64, 95.07, 60.38, 56.88, 55.67, 54.95, 50.09, 47.93, 42.35, 31.06, 24.44. ESI-MS: *m/z* = 416.25 ([M+H]$^+$).

**Synthesis of 4-((1-(3-azidopropanoyl)azetidin-3-yl)amino)-*N*-(2,6-dioxopiperidin-3-yl)-2-methoxybenzamide (X33)**

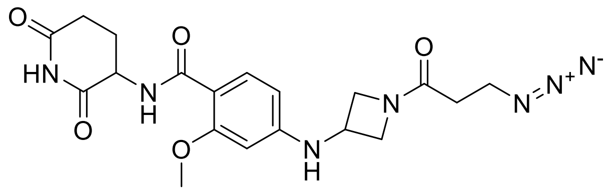

The title compound was prepared according to general procedure A, using **I34** (130 mg, 0.352 mmol), 3-azidopropanoic acid (45 mg, 0.39 mmol), HATU (201 mg, 0.582 mmol) and DIPEA (245 µL, 1.41 mmol). The title compound was obtained as a colourless solid (149 mg, 99%). $^1$H NMR (400 MHz, DMSO-$d_6$) δ 10.86 (s, 1H), 8.38 (d, *J* = 7.0 Hz, 1H), 7.74 (d, *J* = 8.5 Hz, 1H), 6.98 (d, *J* = 6.1 Hz, 1H), 6.25 – 5.99 (m, 2H), 4.76 – 4.63 (m, 1H), 4.51 (t, *J* = 7.8 Hz, 1H), 4.36 – 4.28 (m, 1H), 4.25 (t, *J* = 9.4, 7.5 Hz, 1H), 3.91 (d, *J* = 4.7 Hz, 1H), 3.88 (s, 3H), 3.74 – 3.69 (m, 1H), 3.51 (t, *J* = 6.3 Hz, 2H), 2.75 (ddd, *J* = 17.2, 13.5, 5.7 Hz, 1H), 2.55 – 2.51 (m, 1H), 2.37 (t, *J* = 6.3 Hz, 2H), 2.20 – 1.94 (m, 2H). $^{13}$C NMR (101 MHz, DMSO-$d_6$) δ 172.99, 172.75, 169.80, 164.49, 159.23, 151.47, 132.85, 118.07, 109.47, 104.66, 95.02, 56.98, 55.67, 54.43, 50.10, 46.31, 41.78, 31.07, 30.25, 24.45. ESI-MS: *m/z* = 430.15 ([M+H]$^+$).

**Synthesis of 4-((1-(4-azidobutanoyl)azetidin-3-yl)amino)-*N*-(2,6-dioxopiperidin-3-yl)-2-methoxybenzamide (X34)**

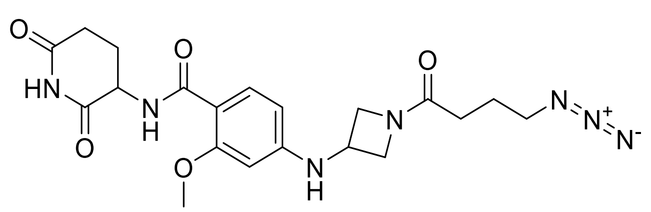

The title compound was prepared according to general procedure A, using **I34** (130 mg, 0.352 mmol), 4-azidobutanoic acid (50 mg, 0.39 mmol), HATU (201 mg, 0.582 mmol) and

DIPEA (245 µL, 1.41 mmol). The title compound was obtained as a colourless solid (127 mg, 77%). $^{1}$H NMR (400 MHz, DMSO-$d_6$) δ 10.86 (s, 1H), 8.38 (d, *J* = 7.0 Hz, 1H), 7.73 (d, *J* = 8.5 Hz, 1H), 6.96 (d, *J* = 6.2 Hz, 1H), 6.28 – 6.11 (m, 2H), 4.69 (quin, *J* = 6.2 Hz, 1H), 4.49 (t, *J* = 7.7 Hz, 1H), 4.33 – 4.25 (m, 1H), 4.25 – 4.18 (m, 1H), 3.88 (s, 3H), 3.87 – 3.84 (m, 2H), 3.67 (dd, *J* = 9.6, 4.8 Hz, 1H), 2.75 (ddd, *J* = 17.2, 13.4, 5.7 Hz, 1H), 2.52 (s, 1H), 2.20 – 1.99 (m, 5H), 1.73 (quin, *J* = 7.1 Hz, 2H). $^{13}$C NMR (101 MHz, DMSO-$d_6$) δ 172.98, 172.75, 171.44, 164.48, 159.22, 151.49, 132.84, 109.45, 104.66, 95.00, 56.97, 55.66, 54.36, 50.18, 50.09, 41.71, 31.06, 27.55, 24.45, 23.56. ESI-MS: *m/z* = 444.20 ([M+H]$^+$).

**Synthesis of *tert*-butyl 4-(4-((2,6-dioxopiperidin-3-yl)carbamoyl)-3-methoxyphenyl)-piperazine-1-carboxylate (I35)**

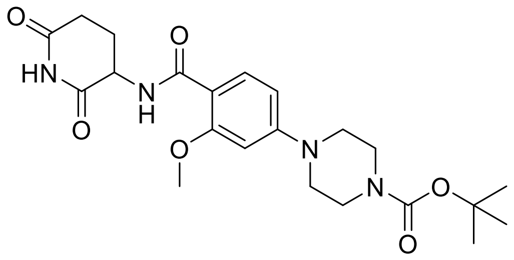


**I32** (1.50 g, 3.86 mmol), *tert*-butyl piperazine-1-carboxylate (1.44 g, 7.73 mmol), copper iodide (147 mg, 0.775 mmol), *L*-proline (178 mg, 1.55 mmol) and potassium carbonate (1.60 g, 11.6 mmol) were dissolved in dry DMSO (10 mL) and stirred at 80 °C for 18 h. The solvent was removed under reduced pressure and the crude product was purified by flash chromatography using acetonitrile/water as an eluent. The title compound was obtained as a colourless solid (415 mg, 24%). $^{1}$H NMR (400 MHz, DMSO-$d_6$) δ 10.87 (s, 1H), 8.44 (d, *J* = 7.0 Hz, 1H), 7.79 (d, *J* = 8.8 Hz, 1H), 6.60 (dd, *J* = 8.9, 2.2 Hz, 1H), 6.56 (d, *J* = 2.3 Hz, 1H), 4.79 – 4.61 (m, 1H), 3.92 (s, 3H), 3.50 – 3.41 (m, 4H), 3.31 – 3.27 (m, 4H), 2.82 – 2.67 (m, 1H), 2.57 – 2.51 (m, 1H), 2.19 – 1.96 (m, 2H), 1.42 (s, 9H). ESI-MS: *m/z* = 447.15 ([M+H]$^+$).

**Synthesis of *N*-(2,6-dioxopiperidin-3-yl)-2-methoxy-4-(piperazin-1-yl)benzamide (I36)**

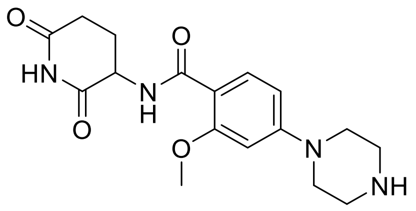


The title compound was prepared according to general procedure D, using **I35** (415 mg, 0.929 mmol). The crude product was used without further purification. ESI-MS: *m/z* = 347.05 ([M+H]$^+$).

**Synthesis of 4-(4-(2-azidoacetyl)piperazin-1-yl)-*N*-(2,6-dioxopiperidin-3-yl)-2-methoxy-benzamide (X35)**

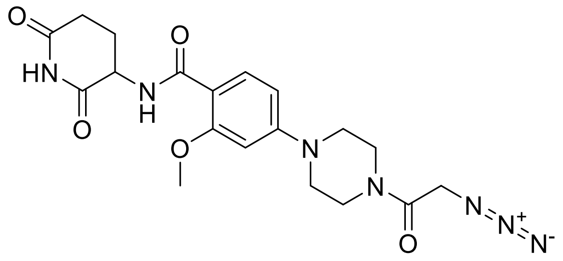


The title compound was prepared according to general procedure A, using **I36** (115 mg, 0.301 mmol), 2-azidoacetic acid (33 mg, 0.33 mmol), HATU (171 mg, 0.451 mmol) and DIPEA (157 µL, 0.902 mmol). The title compound was obtained as a colourless solid (78 mg, 60%). $^1$H NMR (400 MHz, DMSO-$d_6$) δ 10.87 (s, 1H), 8.45 (d, *J* = 7.0 Hz, 1H), 7.80 (d, *J* = 8.8 Hz, 1H), 6.61 (dd, *J* = 8.9, 2.2 Hz, 1H), 6.56 (d, *J* = 2.2 Hz, 1H), 4.70 (dt, *J* = 12.4, 6.3 Hz, 1H), 4.21 (s, 2H), 3.93 (s, 3H), 3.73 – 3.59 (m, 2H), 3.53 – 3.44 (m, 2H), 3.39 – 3.34 (m, 4H), 2.91 – 2.66 (m, 1H), 2.55 – 2.52 (m, 1H), 2.19 – 1.94 (m, 2H). $^{13}$C NMR (101 MHz, DMSO-$d_6$) δ 172.99, 172.68, 166.13, 164.28, 158.90, 154.05, 132.41, 110.97, 106.71, 97.78, 55.93, 50.10, 49.68, 46.85, 46.74, 43.55, 41.02, 31.05, 24.38. ESI-MS: *m/z* = 430.20 ([M+H]$^+$).

**Synthesis of 4-(4-(4-azidobutanoyl)piperazin-1-yl)-*N*-(2,6-dioxopiperidin-3-yl)-2-methoxybenzamide (X36)**

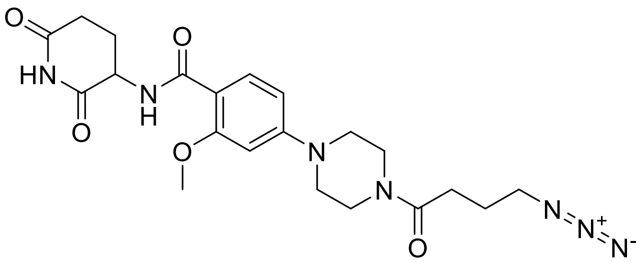


The title compound was prepared according to general procedure A, using **I36** (200 mg, 0.721 mmol), 4-azidobutanoic acid (125 mg, 1.08 mmol), HATU (356 mg, 0.937 mmol) and DIPEA (377 µL, 2.16 mmol). The title compound was obtained as a colourless solid (156 mg, 58%). $^{1}$H NMR (400 MHz, DMSO-$d_6$) δ 10.86 (s, 1H), 8.44 (d, *J* = 7.0 Hz, 1H), 7.80 (d, *J* = 8.8 Hz, 1H), 6.61 (dd, *J* = 8.9, 2.2 Hz, 1H), 6.56 (d, *J* = 2.2 Hz, 1H), 4.70 (ddd, *J* = 12.4, 7.0, 5.6 Hz, 1H), 3.93 (s, 3H), 3.60 (q, *J* = 5.1 Hz, 4H), 3.37 (t, *J* = 6.9 Hz, 4H), 3.31 – 3.28 (m, 2H), 2.76 (ddd, *J* = 17.3, 13.4, 5.8 Hz, 1H), 2.56 – 2.51 (m, 1H), 2.44 (t, *J* = 7.2 Hz, 2H), 2.21 – 1.99 (m, 2H), 1.78 (quin, *J* = 7.1 Hz, 2H). $^{13}$C NMR (101 MHz, DMSO-$d_6$) δ 172.94, 172.64, 169.95, 164.28, 158.88, 154.14, 132.37, 110.91, 106.68, 97.71, 55.90, 50.29, 50.09, 47.12, 46.85, 44.26, 40.64, 31.03, 29.09, 24.37, 24.03. ESI-MS: *m/z* = 458.15 ([M+H]$^+$).

**Synthesis of (2*S*,4*R*)-1-((*S*)-2-(4-azidobenzamido)-3,3-dimethylbutanoyl)-4-hydroxy-*N*-(4-(4-methylthiazol-5-yl)benzyl)pyrrolidine-2-carboxamide (X37)**

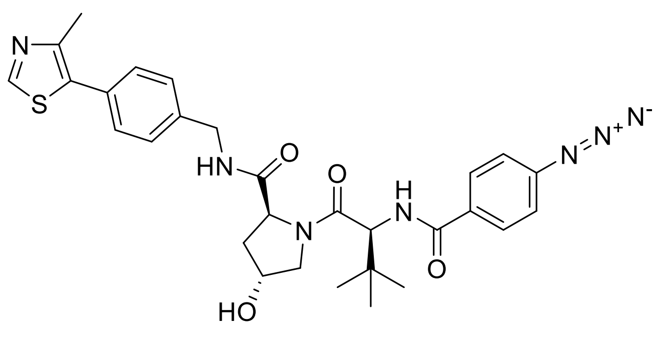


The title compound was prepared according to general procedure A, using (2*S*,4*R*)-1-((*S*)-2-amino-3,3-dimethylbutanoyl)-4-hydroxy-*N*-(4-(4-methylthiazol-5-yl)benzyl)pyrrolidine-2-carboxamide (100 mg, 0.232 mmol), 4-azidobenzoic acid (38 mg, 0.23 mmol), HATU (115 mg, 0.302 mmol) and DIPEA (121 µL, 0.697 mmol). The title compound was obtained as a colourless solid (110 mg, 82%). $^{1}$H NMR (400 MHz, DMSO-$d_6$) δ 8.98 (s, 1H), 8.58 (t, *J* = 6.1 Hz, 1H), 8.05 (d, *J* = 9.0 Hz, 1H), 7.94 (d, *J* = 8.2 Hz, 2H),

7.44 – 7.36 (m, 4H), 7.18 (d, *J* = 8.2 Hz, 2H), 5.15 (d, *J* = 3.6 Hz, 1H), 4.77 (d, *J* = 9.0 Hz, 1H), 4.50 – 4.35 (m, 3H), 4.24 (dd, *J* = 15.9, 5.5 Hz, 1H), 3.73 (s, 2H), 2.45 (s, 3H), 2.16 – 1.85 (m, 2H), 1.03 (s, 9H). $^{13}$C NMR (101 MHz, DMSO-$d_6$) δ 171.91, 169.45, 165.60, 151.47, 147.75, 142.39, 139.49, 131.17, 130.62, 129.68, 128.69, 127.47, 118.76, 68.92, 58.82, 57.37, 56.43, 41.68, 37.95, 35.55, 26.53, 15.95. ESI-MS: *m/z* = 476.20 ([M+H]$^+$).

**Synthesis of (2*S*,4*R*)-1-((*S*)-2-(2-azidoacetamido)-3,3-dimethylbutanoyl)-4-hydroxy-*N*-(4-(4-methylthiazol-5-yl)benzyl)pyrrolidine-2-carboxamide (X38)**

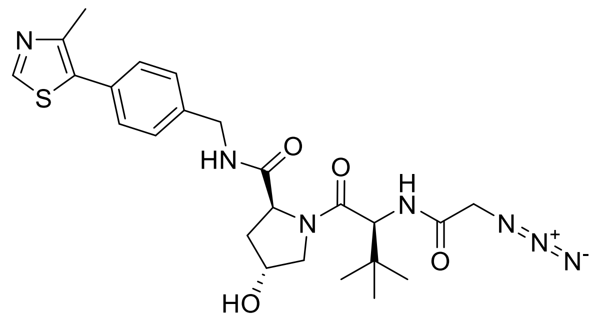


The title compound was prepared according to general procedure A, using (2*S*,4*R*)-1-((*S*)-2-amino-3,3-dimethylbutanoyl)-4-hydroxy-*N*-(4-(4-methylthiazol-5-yl)benzyl)pyrrolidine-2-carboxamide (100 mg, 0.232 mmol), 2-azidoacetic acid (23 mg, 0.23 mmol), HATU (115 mg, 0.302 mmol) and DIPEA (121 µL, 0.697 mmol). The title compound was obtained as a colourless solid (105 mg, 88%). $^1$H NMR (400 MHz, DMSO-$d_6$) δ 8.98 (s, 1H), 8.60 (t, *J* = 6.0 Hz, 1H), 8.23 (d, *J* = 9.3 Hz, 1H), 7.45 – 7.35 (m, 4H), 5.17 (d, *J* = 3.5 Hz, 1H), 4.56 (d, *J* = 9.3 Hz, 1H), 4.48 – 4.40 (m, 2H), 4.36 (s, 1H), 4.21 (dd, *J* = 15.9, 5.4 Hz, 1H), 3.99 – 3.84 (m, 2H), 3.73 – 3.59 (m, 2H), 2.44 (s, 3H), 2.04 (d, *J* = 8.6 Hz, 1H), 1.95 – 1.86 (m, 1H), 0.95 (s, 9H). $^{13}$C NMR (101 MHz, DMSO-$d_6$) δ 171.88, 169.10, 167.38, 151.48, 147.74, 139.50, 131.17, 129.66, 128.65, 127.42, 68.90, 58.76, 56.58, 50.32, 41.66, 37.95, 35.48, 26.26, 15.95. ESI-MS: *m/z* = 514.20 ([M+H]$^+$).

**Synthesis of (2*S*,4*R*)-1-((*S*)-2-(4-azidobutanamido)-3,3-dimethylbutanoyl)-4-hydroxy-*N*-(4-(4-methylthiazol-5-yl)benzyl)pyrrolidine-2-carboxamide (X39)**

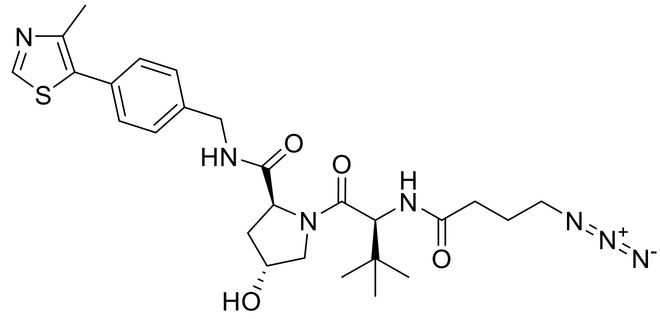


The title compound was prepared according to general procedure A, using (2*S*,4*R*)-1-((*S*)-2-amino-3,3-dimethylbutanoyl)-4-hydroxy-*N*-(4-(4-methylthiazol-5-yl)benzyl)pyrrolidine-2-carboxamide (100 mg, 0.232 mmol), 4-azidobutanoic acid (30 mg, 0.23 mmol), HATU (115 mg, 0.302 mmol) and DIPEA (121 µL, 0.697 mmol). The title compound was obtained as a colourless solid (114 mg, 91%). $^{1}$H NMR (400 MHz, DMSO-$d_6$) δ 8.97 (s, 1H), 8.56 (t, *J* = 6.1 Hz, 1H), 7.97 (d, *J* = 9.3 Hz, 1H), 7.46 – 7.27 (m, 4H), 5.14 (d, *J* = 3.5 Hz, 1H), 4.54 (d, *J* = 9.3 Hz, 1H), 4.47 – 4.39 (m, 2H), 4.36 – 4.32 (m, 1H), 4.21 (dd, *J* = 15.9, 5.4 Hz, 1H), 3.72 – 3.58 (m, 2H), 3.30 (td, *J* = 6.9, 1.1 Hz, 2H), 2.44 (s, 3H), 2.40 – 2.18 (m, 2H), 2.10 – 1.98 (m, 1H), 1.95 – 1.85 (m, 1H), 1.81 – 1.66 (m, 2H), 0.93 (s, 9H). $^{13}$C NMR (101 MHz, DMSO-$d_6$) δ 172.04, 171.35, 169.69, 151.53, 147.78, 139.55, 131.24, 129.70, 128.70, 127.49, 68.94, 58.77, 56.51, 56.42, 50.36, 41.71, 37.98, 35.27, 31.86, 26.40, 24.73, 15.98. ESI-MS: *m/z* = 542.35 ([M+H]$^{+}$).

**Synthesis of methyl 4-(((*S*)-1-((2*S*,4*R*)-4-hydroxy-2-((4-(4-methylthiazol-5-yl)benzyl)carbamoyl)pyrrolidin-1-yl)-3,3-dimethyl-1-oxobutan-2-yl)carbamoyl)benzoate (I37)**

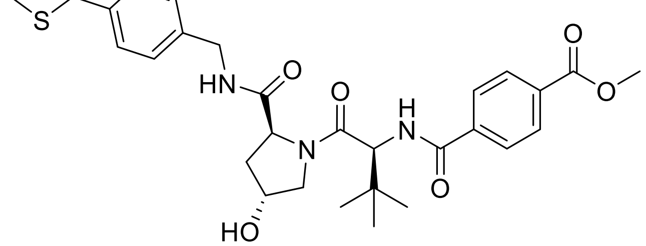


The title compound was prepared according to general procedure A, using (2*S*,4*R*)-1-((*S*)-2-amino-3,3-dimethylbutanoyl)-4-hydroxy-*N*-(4-(4-methylthiazol-5-

yl)benzyl)pyrrolidine-2-carboxamide (1.00 g, 2.32 mmol), 4-(methoxycarbonyl)benzoic acid (418 mg, 2.32 mmol), HATU (1.15 g, 3.02 mmol) and DIPEA (1.21 mL, 6.97 mmol). The title compound was obtained as a colourless solid (1.29 g, 94%). $^1$H NMR (400 MHz, DMSO-$d_6$) δ 8.98 (s, 1H), 8.58 (t, *J* = 6.0 Hz, 1H), 8.29 (d, *J* = 9.1 Hz, 1H), 8.10 – 7.92 (m, 4H), 7.45 – 7.35 (m, 4H), 5.15 (d, *J* = 3.6 Hz, 1H), 4.78 (d, *J* = 9.1 Hz, 1H), 4.48 – 4.40 (m, 2H), 4.39 – 4.35 (m, 1H), 4.24 (dd, *J* = 15.8, 5.5 Hz, 1H), 3.88 (s, 3H), 3.73 (d, *J* = 3.0 Hz, 2H), 2.45 (s, 3H), 2.11 – 2.00 (m, 1H), 1.98 – 1.86 (m, 1H), 1.04 (s, 9H). ESI-MS: *m/z* = 615.20 ([M+Na]$^+$).

**Synthesis of 4-(((*S*)-1-((2*S*,4*R*)-4-hydroxy-2-((4-(4-methylthiazol-5-yl)benzyl)carbamoyl)pyrrolidin-1-yl)-3,3-dimethyl-1-oxobutan-2-yl)carbamoyl)benzoic acid (I38)**

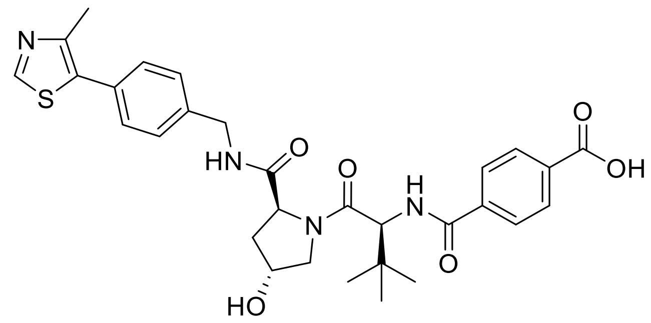


**I37** (1.12 g, 1.83 mmol) and LiOH·H2O (308 mg, 7.34 mmol) were dissolved in MeOH (10 mL) and $H_2O$ (5 mL) and stirred at rt for 16 h. Afterwards, the pH was brought to 1 through the addition of aqueous HCl (10%) and most of the organic solvent was removed under reduced pressure. The residue was partitioned between water and ethyl acetate and the aqueous phase was extracted with ethyl acetate (3x). The combined organic phase was washed with brine, dried over $MgSO_4$, filtered, and volatiles were removed under reduced pressure to provide the title compound (1.03 g, 97%). $^1$H NMR (400 MHz, DMSO-$d_6$) δ 13.00 (s, 1H), 8.98 (s, 1H), 8.59 (t, *J* = 6.1 Hz, 1H), 8.24 (d, *J* = 9.1 Hz, 1H), 8.03 – 7.93 (m, 4H), 7.45 – 7.31 (m, 4H), 4.78 (d, *J* = 9.1 Hz, 1H), 4.51 – 4.35 (m, 3H), 4.24 (dd, *J* = 15.8, 5.5 Hz, 1H), 3.74 (d, *J* = 3.0 Hz, 2H), 2.45 (s, 3H), 2.09 – 2.02 (m, 1H), 1.97 – 1.91 (m, 1H), 1.04 (s, 9H). ESI-MS: *m/z* = 579.30 ([M+H]$^+$).

**Synthesis of *N*[1]-(3-azidopropyl)-*N*[4]-((*S*)-1-((2*S*,4*R*)-4-hydroxy-2-((4-(4-methylthiazol-5-yl)benzyl)carbamoyl)pyrrolidin-1-yl)-3,3-dimethyl-1-oxobutan-2-yl)terephthalamide (X40)**

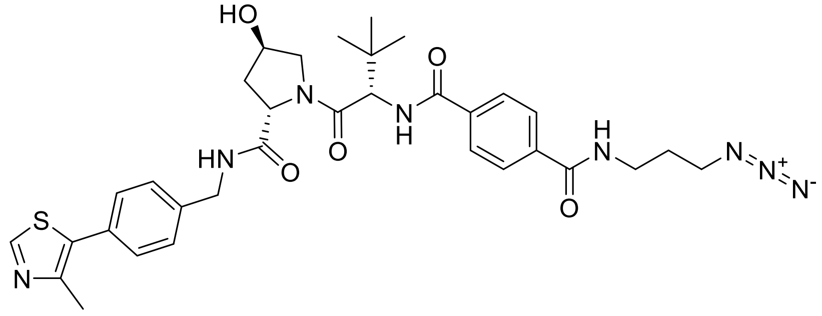

The title compound was prepared according to general procedure A, using **I38** (180 mg, 0.311 mmol), 3-azidopropan-1-amine (31 mg, 0.31 mmol), HATU (154 mg, 0.404 mmol) and DIPEA (163 µL, 0.933 mmol). The title compound was obtained as a colourless solid (191 mg, 93%). $^{1}$H NMR (400 MHz, DMSO-$d_6$) δ 8.98 (s, 1H), 8.63 (dt, *J* = 24.2, 5.9 Hz, 2H), 8.17 (d, *J* = 9.0 Hz, 1H), 7.93 (q, *J* = 8.4 Hz, 4H), 7.46 – 7.32 (m, 4H), 5.18 (d, *J* = 3.6 Hz, 1H), 4.80 (d, *J* = 9.1 Hz, 1H), 4.53 – 4.36 (m, 3H), 4.25 (dd, *J* = 15.7, 5.5 Hz, 1H), 3.75 (s, 2H), 3.42 (t, *J* = 6.8 Hz, 2H), 3.35 (q, *J* = 6.7 Hz, 2H), 2.45 (s, 3H), 2.13 – 2.00 (m, 1H), 1.99 – 1.89 (m, 1H), 1.80 (quin, *J* = 6.8 Hz, 2H), 1.05 (s, 9H). $^{13}$C NMR (101 MHz, DMSO-$d_6$) δ 171.96, 169.42, 166.05, 165.71, 151.48, 147.78, 139.51, 136.83, 136.34, 131.21, 129.72, 128.73, 127.72, 127.51, 127.08, 68.98, 58.89, 57.49, 56.52, 48.60, 41.74, 38.26, 37.99, 36.78, 35.64, 28.38, 26.57, 15.97. ESI-MS: *m/z* = 661.35 ([M+H]$^+$).

**Synthesis of *N*[1]-(4-azidobutyl)-*N*[4]-((*S*)-1-((2*S*,4*R*)-4-hydroxy-2-((4-(4-methylthiazol-5-yl)benzyl)carbamoyl)pyrrolidin-1-yl)-3,3-dimethyl-1-oxobutan-2-yl)terephthalamide (X41)**

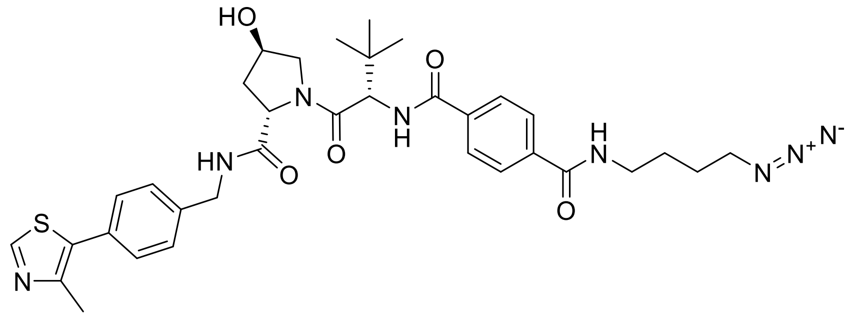

The title compound was prepared according to general procedure A, using **I38** (180 mg, 0.311 mmol), 4-azidobutan-1-amine (36 mg, 0.31 mmol), HATU (154 mg, 0.404 mmol) and DIPEA (163 µL, 0.933 mmol). The title compound was obtained as a colourless solid (190 mg, 90%). $^{1}$H NMR (400 MHz, DMSO-*d*$_6$) δ 8.98 (s, 1H), 8.60 (dt, *J* = 13.8, 5.9 Hz, 2H), 8.15 (d, *J* = 9.0 Hz, 1H), 7.98 – 7.85 (m, 4H), 7.46 – 7.37 (m, 4H), 5.15 (d, *J* = 3.6 Hz, 1H), 4.79 (d, *J* = 9.1 Hz, 1H), 4.50 – 4.37 (m, 3H), 4.25 (dd, *J* = 15.8, 5.6 Hz, 1H), 3.74 (d, *J* = 3.1 Hz, 2H), 3.38 – 3.35 (m, 2H), 3.31 – 3.26 (m, 2H), 2.45 (s, 3H), 2.12 – 2.02 (m, 1H), 1.98 – 1.86 (m, 1H), 1.59 (quin, *J* = 3.4 Hz, 4H), 1.04 (s, 9H). $^{13}$C NMR (101 MHz, DMSO-*d*$_6$) δ 171.88, 169.34, 165.98, 165.49, 151.42, 147.73, 139.47, 136.89, 136.24, 131.15, 129.67, 128.68, 127.65, 127.46, 126.97, 68.91, 58.82, 57.42, 56.44, 50.37, 41.68, 38.67, 37.94, 35.57, 26.52, 26.30, 25.82, 15.92. ESI-MS: *m/z* = 675.40 ([M+H]$^+$).

**Synthesis of *N*$^1$-(6-azidohexyl)-*N*$^4$-((*S*)-1-((2*S*,4*R*)-4-hydroxy-2-((4-(4-methylthiazol-5-yl)benzyl)carbamoyl)pyrrolidin-1-yl)-3,3-dimethyl-1-oxobutan-2-yl)terephthalamide (X42)**

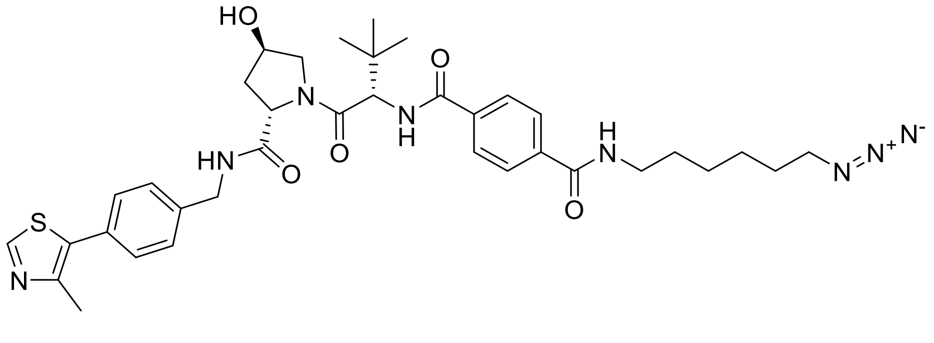

The title compound was prepared according to general procedure A, using **I38** (180 mg, 0.311 mmol), 6-azidohexan-1-amine (44 mg, 0.31 mmol), HATU (154 mg, 0.404 mmol) and DIPEA (163 µL, 0.933 mmol). The title compound was obtained as a colourless solid (179 mg, 58%). $^{1}$H NMR (400 MHz, DMSO-*d*$_6$) δ 8.98 (s, 1H), 8.57 (q, *J* = 6.3 Hz, 2H), 8.15 (d, *J* = 9.0 Hz, 1H), 7.92 (q, *J* = 8.4 Hz, 4H), 7.53 – 7.34 (m, 4H), 5.15 (d, *J* = 3.6 Hz, 1H), 4.79 (d, *J* = 9.1 Hz, 1H), 4.53 – 4.35 (m, 3H), 4.25 (dd, *J* = 15.8, 5.5 Hz, 1H), 3.74 (s, 2H), 3.31 – 3.23 (m, 4H), 2.45 (s, 3H), 2.14 – 2.02 (m, 1H), 1.99 – 1.89 (m, 1H), 1.54 (quin, *J* = 6.2 Hz, 4H), 1.37 – 1.29 (m, 4H), 1.05 (s, 9H). $^{13}$C NMR (101 MHz, DMSO-*d*$_6$) δ 171.88, 169.35, 165.98, 165.39, 151.41, 147.73, 139.46, 136.99, 136.18, 131.15, 129.68, 128.68, 127.62,

127.46, 126.97, 68.91, 58.82, 57.41, 56.44, 50.58, 41.69, 37.94, 35.58, 28.89, 28.17, 26.52, 26.00, 25.88, 15.92. ESI-MS: *m/z* = 703.40 ([M+H]$^+$).

**Synthesis of *N*[1]-(2-(2-(2-azidoethoxy)ethoxy)ethyl)-*N*[4]-((*S*)-1-((2*S*,4*R*)-4-hydroxy-2-((4-(4-methylthiazol-5-yl)benzyl)carbamoyl)pyrrolidin-1-yl)-3,3-dimethyl-1-oxobutan-2-yl)terephthalamide (X43)**

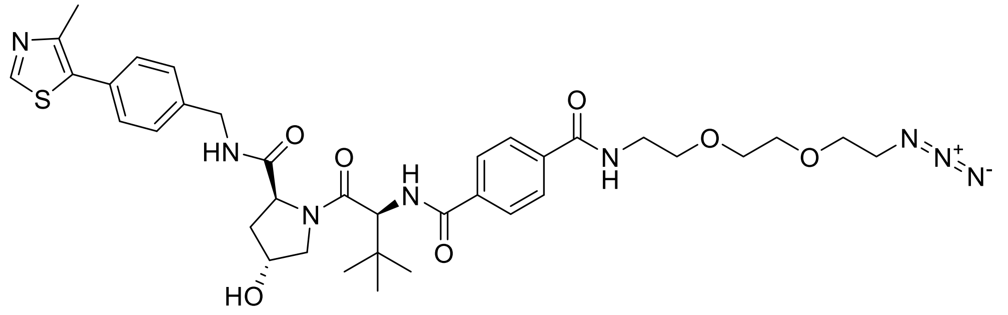

The title compound was prepared according to general procedure B, using **I38** (300 mg, 0.518 mmol) and 1-(2-aminoethoxy)-2-(2-azidoethoxy)ethane (99 mg, 0.57 mmol), 1-methyl-1*H*-imidazole (145 µL, 1.81 mmol) and TCFH (175 mg, 0.622 mmol). The title compound was obtained as a colourless solid (137 mg, 36%). $^1$H NMR (500 MHz, $CDCl_3$) δ 8.68 (s, 1H), 7.86 – 7.75 (m, 4H), 7.42 – 7.32 (m, 4H), 7.23 (t, *J* = 6.1 Hz, 1H), 6.86 – 6.80 (m, 2H), 4.79 – 4.70 (m, 2H), 4.63 – 4.53 (m, 2H), 4.35 (dd, *J* = 14.9, 5.3 Hz, 1H), 4.16 (d, *J* = 11.2 Hz, 1H), 3.70 – 3.62 (m, 12H), 3.36 (t, *J* = 4.9 Hz, 2H), 2.66 – 2.57 (m, 1H), 2.52 (s, 3H), 2.17 – 2.10 (m, 1H), 1.01 (s, 9H). $^{13}$C NMR (126 MHz, $CDCl_3$) δ 171.85, 170.59, 167.01, 166.66, 150.47, 148.69, 138.10, 137.81, 136.11, 131.68, 131.26, 129.74, 128.33, 127.53, 127.46, 70.67, 70.41, 70.38, 70.21, 69.90, 58.61, 58.12, 56.93, 50.74, 43.50, 40.01, 35.90, 35.57, 26.66, 16.22. ESI-MS: *m/z* = 735.4 ([M+H]$^+$).

**Synthesis of *N*[1]-(2-(2-(2-(2-azidoethoxy)ethoxy)ethoxy)ethyl)-*N*[4]-((*S*)-1-((2*S*,4*R*)-4-hydroxy-2-((4-(4-methylthiazol-5-yl)benzyl)carbamoyl)pyrrolidin-1-yl)-3,3-dimethyl-1-oxobutan-2-yl)terephthalamide (X44)**

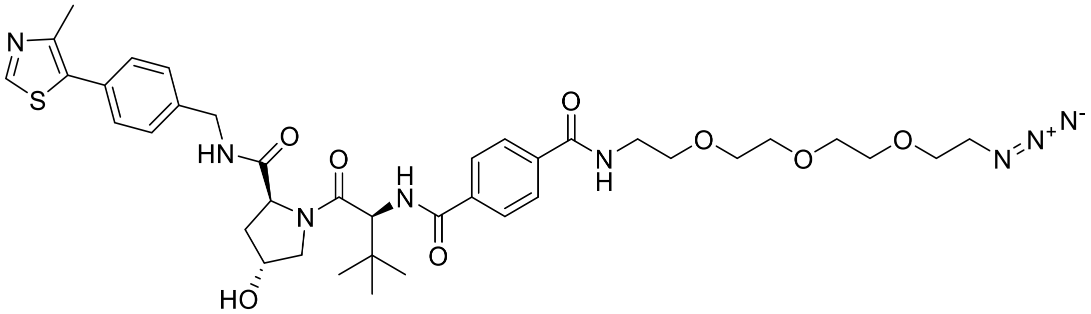

The title compound was prepared according to general procedure B, using **I38** (200 mg, 0.346 mmol) and 1-[2-(2-aminoethoxy)ethoxy]-2-(2-azidoethoxy)ethane (83 mg, 0.38 mmol), 1-methyl-1*H*-imidazole (96 µL, 1.2 mmol) and TCFH (116 mg, 0.415 mmol). The title compound was obtained as a colourless solid (199 mg, 74%). $^{1}$H NMR (500 MHz, $CDCl_3$) δ 8.68 (s, 1H), 7.91 – 7.64 (m, 4H), 7.42 – 7.33 (m, 4H), 7.24 (t, *J* = 5.9 Hz, 1H), 7.04 (t, *J* = 4.5 Hz, 1H), 6.90 (d, *J* = 8.8 Hz, 1H), 4.77 – 4.69 (m, 2H), 4.57 (dd, *J* = 14.8, 6.4 Hz, 2H), 4.36 (dd, *J* = 14.9, 5.3 Hz, 1H), 4.15 (dt, *J* = 11.5, 1.8 Hz, 1H), 3.73 – 3.57 (m, 16H), 3.33 (t, *J* = 5.0 Hz, 2H), 2.62 – 2.53 (m, 1H), 2.52 (s, 3H), 2.17 – 2.09 (m, 1H), 1.00 (s, 9H). $^{13}$C NMR (126 MHz, $CDCl_3$) δ 171.81, 170.68, 166.94, 166.67, 150.47, 148.68, 138.12, 137.80, 136.02, 131.68, 131.24, 129.73, 128.30, 127.55, 127.42, 70.79, 70.77, 70.66, 70.39, 70.34, 70.15, 69.83, 58.67, 58.09, 57.00, 50.77, 43.47, 40.05, 36.03, 35.68, 26.67, 16.22. ESI-MS: *m/z* = 779.4 ([M+H]$^+$).

**Synthesis of *tert*-butyl 4-(((*S*)-1-((2*S*,4*R*)-4-hydroxy-2-((4-(4-methylthiazol-5-yl)benzyl)carbamoyl)pyrrolidin-1-yl)-3,3-dimethyl-1-oxobutan-2-yl)amino)-4-oxobutanoate (I39)**

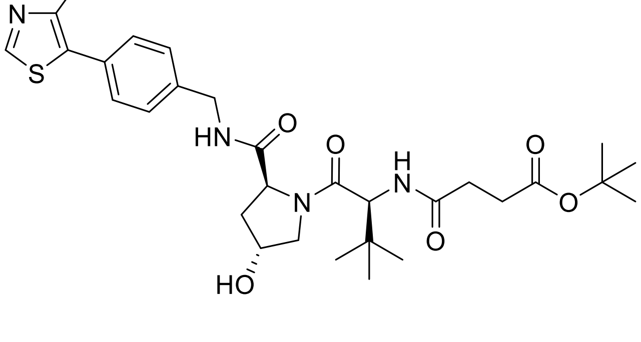

The title compound was prepared according to general procedure A, using (2*S*,4*R*)-1-((*S*)-2-amino-3,3-dimethylbutanoyl)-4-hydroxy-*N*-(4-(4-methylthiazol-5-

yl)benzyl)pyrrolidine-2-carboxamide (1.00 g, 2.32 mmol), 4-(*tert*-butoxy)-4-oxobutanoic acid (405 mg, 2.32 mmol), HATU (1.15 g, 3.02 mmol) and DIPEA (1.21 mL, 6.97 mmol). The title compound was obtained as a colourless solid (934 mg, 69%). $^1$H NMR (400 MHz, DMSO-$d_6$) δ 8.98 (s, 1H), 8.56 (t, $J$ = 6.0 Hz, 1H), 7.93 (d, $J$ = 9.4 Hz, 1H), 7.46 – 7.33 (m, 4H), 5.11 (d, $J$ = 3.5 Hz, 1H), 4.54 (d, $J$ = 9.3 Hz, 1H), 4.47 – 4.39 (m, 2H), 4.35 (s, 1H), 4.21 (dd, $J$ = 15.9, 5.4 Hz, 1H), 3.70 – 3.54 (m, 2H), 2.44 (s, 3H), 2.41 – 2.28 (m, 4H), 2.06 – 1.98 (m, 1H), 1.95 – 1.85 (m, 1H), 1.37 (s, 9H), 0.93 (s, 9H). ESI-MS: $m/z$ = 609.30 ([M+Na]$^+$).

**Synthesis of 4-(((*S*)-1-((2*S*,4*R*)-4-hydroxy-2-((4-(4-methylthiazol-5-yl)benzyl)carbamoyl)-pyrrolidin-1-yl)-3,3-dimethyl-1-oxobutan-2-yl)amino)-4-oxobutanoic acid (I40)**

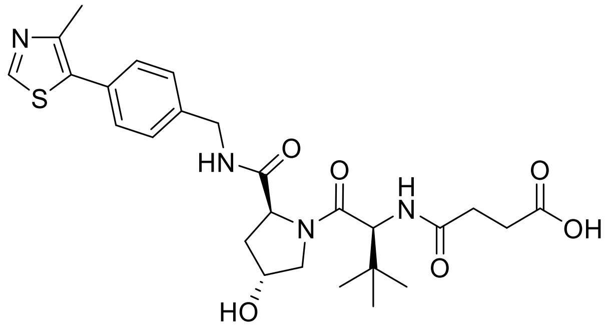


The title compound was prepared according to general procedure D, using **I39** (467 mg, 0.796 mmol). The crude product was used without further purification. ESI-MS: $m/z$ = 531.25 ([M+H]$^+$).

**Synthesis of *N*$^1$-(3-azidopropyl)-*N*$^4$-((*S*)-1-((2*S*,4*R*)-4-hydroxy-2-((4-(4-methylthiazol-5-yl)benzyl)carbamoyl)pyrrolidin-1-yl)-3,3-dimethyl-1-oxobutan-2-yl)succinamide (X45)**

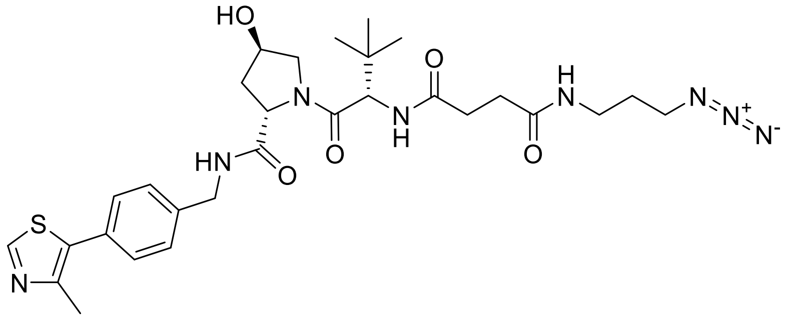

The title compound was prepared according to general procedure A, using **I40** (96 mg, 0.18 mmol), 3-azidopropan-1-amine (18 mg, 0.18 mmol), HATU (90 mg, 0.24 mmol) and DIPEA (95 µL, 0.54 mmol). The title compound was obtained as a colourless solid (99 mg, 89%).

$^{1}H$ NMR (400 MHz, DMSO-$d_6$) δ 8.98 (s, 1H), 8.56 (t, *J* = 6.0 Hz, 1H), 7.97 – 7.82 (m, 2H), 7.44 – 7.36 (m, 4H), 5.12 (d, *J* = 3.5 Hz, 1H), 4.52 (d, *J* = 9.3 Hz, 1H), 4.47 – 4.38 (m, 2H), 4.35 (s, 1H), 4.22 (dd, *J* = 15.8, 5.5 Hz, 1H), 3.71 – 3.55 (m, 2H), 3.36 – 3.34 (m, 3H), 3.08 (qd, *J* = 6.7, 2.6 Hz, 2H), 2.44 (s, 3H), 2.40 – 2.21 (m, 3H), 2.07 – 2.00 (m, 1H), 1.94 – 1.86 (m, 1H), 1.63 (quin, *J* = 6.8 Hz, 2H), 0.93 (s, 9H). $^{13}C$ NMR (101 MHz, DMSO-$d_6$) $^{13}C$ NMR (101 MHz, DMSO) δ 171.95, 171.46, 169.59, 151.46, 147.73, 139.51, 131.17, 129.66, 128.66, 127.44, 68.89, 58.71, 56.40, 41.65, 37.94, 35.81, 35.34, 30.98, 30.55, 28.44, 26.34. ESI-MS: *m/z* = 635.30 ($[M+Na]^+$).

**Synthesis of *tert*-butyl 5-(((*S*)-1-((2*S*,4*R*)-4-hydroxy-2-((4-(4-methylthiazol-5-yl)benzyl)carbamoyl)pyrrolidin-1-yl)-3,3-dimethyl-1-oxobutan-2-yl)amino)-5-oxopentanoate (I41)**

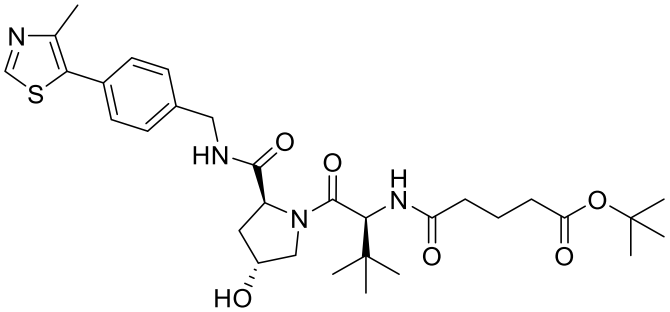


The title compound was prepared according to general procedure A, using (2*S*,4*R*)-1-((*S*)-2-amino-3,3-dimethylbutanoyl)-4-hydroxy-*N*-(4-(4-methylthiazol-5-yl)benzyl)pyrrolidine-2-carboxamide (1.00 g, 2.32 mmol), 5-(*tert*-butoxy)-4-oxopentanoic acid (437 mg, 2.32 mmol), HATU (1.15 g, 3.02 mmol) and DIPEA (1.21 mL, 6.97 mmol). The title compound was obtained as a colourless solid (981 mg, 70%). $^{1}H$ NMR (400 MHz, DMSO-$d_6$) δ 8.98 (s, 1H), 8.56 (t, *J* = 6.1 Hz, 1H), 7.90 (d, *J* = 9.3 Hz, 1H), 7.51 – 7.24 (m, 4H), 5.12 (d, *J* = 3.6 Hz, 1H), 4.54 (d, *J* = 9.3 Hz, 1H), 4.48 – 4.40 (m, 2H), 4.35 (s, 1H), 4.21

(dd, $J$ = 15.8, 5.4 Hz, 1H), 3.70 – 3.53 (m, 2H), 2.44 (s, 3H), 2.32 – 2.21 (m, 1H), 2.21 – 2.11 (m, 3H), 2.02 (d, $J$ = 8.6 Hz, 1H), 1.94 – 1.84 (m, 1H), 1.76 – 1.62 (m, 2H), 1.39 (s, 9H), 0.94 (s, 9H). ESI-MS: $m/z$ = 623.35 ($[M+Na]^+$).

**Synthesis of 5-(((*S*)-1-((2*S*,4*R*)-4-hydroxy-2-((4-(4-methylthiazol-5-yl)benzyl)carbamoyl)pyrrolidin-1-yl)-3,3-dimethyl-1-oxobutan-2-yl)amino)-5-oxopentanoic acid (I42)**

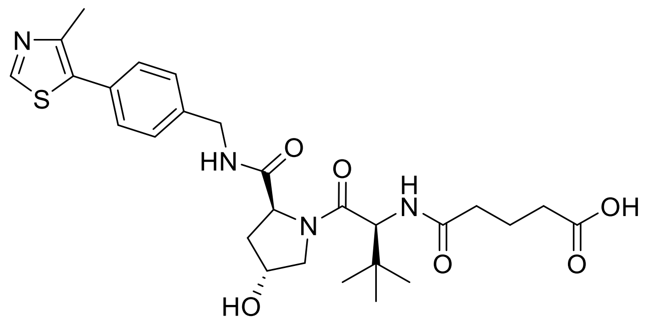

The title compound was prepared according to general procedure D, using **I41** (484 mg, 0.806 mmol). The crude product was used without further purification. ESI-MS: $m/z$ = 545.30 ($[M+H]^+$).

**Synthesis of *N*[1]-(4-azidobutyl)-*N*[5]-((*S*)-1-((2*S*,4*R*)-4-hydroxy-2-((4-(4-methylthiazol-5-yl)benzyl)carbamoyl)pyrrolidin-1-yl)-3,3-dimethyl-1-oxobutan-2-yl)glutaramide (X46)**

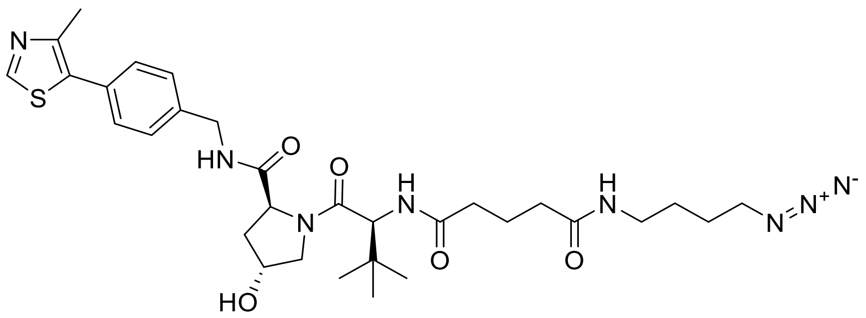

The title compound was prepared according to general procedure A, using **I42** (143 mg, 0.263 mmol), 4-azidobutan-1-amine (30 mg, 0.26 mmol), HATU (130 mg, 0.341 mmol) and DIPEA (137 µL, 0.788 mmol). The title compound was obtained as a colourless solid

(131 mg, 78%). $^1$H NMR (400 MHz, DMSO-$d_6$) δ 8.98 (s, 1H), 8.56 (t, *J* = 6.0 Hz, 1H), 7.87 (d, *J* = 9.3 Hz, 1H), 7.77 (t, *J* = 5.7 Hz, 1H), 7.50 – 7.32 (m, 4H), 5.13 (d, *J* = 3.6 Hz, 1H), 4.53 (d, *J* = 9.3 Hz, 1H), 4.49 – 4.40 (m, 2H), 4.35 (s, 1H), 4.22 (dd, *J* = 15.9, 5.5 Hz, 1H), 3.73 – 3.62 (m, 2H), 3.32 (m, 3H), 3.04 (qd, *J* = 6.8, 2.2 Hz, 2H), 2.44 (s, 3H), 2.29 – 2.10 (m, 2H), 2.07 – 1.99 (m, 3H), 1.95 – 1.85 (m, 1H), 1.75 – 1.63 (m, 2H), 1.56 – 1.38 (m, 3H), 0.94 (s, 9H). $^{13}$C NMR (101 MHz, DMSO-$d_6$) $^{13}$C NMR (101 MHz, DMSO) δ 171.96, 171.75, 171.65, 169.72, 151.46, 147.73, 139.51, 131.17, 129.65, 128.65, 127.43, 68.89, 58.70, 56.38, 50.36, 41.65, 37.94, 37.81, 35.18, 34.95, 34.37, 26.39, 25.76, 21.81, 15.94. ESI-MS: *m/z* = 641.35 ([M+H]$^+$).

**Synthesis of *N*[1]-(6-azidohexyl)-*N*[5]-((*S*)-1-((2*S*,4*R*)-4-hydroxy-2-((4-(4-methylthiazol-5-yl)benzyl)carbamoyl)pyrrolidin-1-yl)-3,3-dimethyl-1-oxobutan-2-yl)glutaramide (X47)**

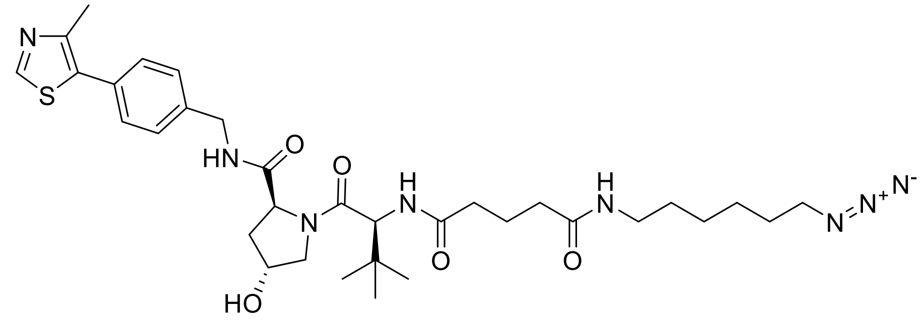


The title compound was prepared according to general procedure A, using **I42** (143 mg, 0.263 mmol), 6-azidohexan-1-amine (37 mg, 0.26 mmol), HATU (130 mg, 0.341 mmol) and DIPEA (137 µL, 0.788 mmol). The title compound was obtained as a colourless solid (131 mg, 78%). $^1$H NMR (400 MHz, DMSO-$d_6$) δ 8.98 (s, 1H), 8.56 (t, *J* = 6.1 Hz, 1H), 7.87 (d, *J* = 9.3 Hz, 1H), 7.72 (t, *J* = 5.6 Hz, 1H), 7.47 – 7.34 (m, 4H), 5.13 (d, *J* = 3.5 Hz, 1H), 4.53 (d, *J* = 9.3 Hz, 1H), 4.47 – 4.39 (m, 2H), 4.35 (s, 1H), 4.22 (dd, *J* = 15.9, 5.5 Hz, 1H), 3.72 – 3.60 (m, 2H), 3.30 (t, *J* = 6.9 Hz, 2H), 3.05 – 2.96 (m, 2H), 2.44 (s, 3H), 2.28 – 2.10 (m, 2H), 2.04 (t, *J* = 7.7 Hz, 3H), 1.97 – 1.86 (m, 1H), 1.74 – 1.63 (m, 2H), 1.51 (quin, *J* = 6.9 Hz, 2H), 1.38 (quin, *J* = 7.3 Hz, 2H), 1.33 – 1.23 (m, 4H), 0.94 (s, 9H). $^{13}$C NMR (101 MHz, DMSO-$d_6$) δ 171.96, 171.77, 171.54, 169.73, 151.45, 147.72, 139.51, 131.17, 129.65, 128.64, 127.43, 68.89, 58.70, 56.39, 56.35, 50.57, 41.65, 38.30, 37.95, 35.17, 34.95, 34.38, 29.01, 28.18, 26.39, 25.93, 25.86, 21.84, 15.94. ESI-MS: *m/z* = 669.45 ([M+H]$^+$).

**Synthesis of *N*[1]-(2-(2-(2-(2-azidoethoxy)ethoxy)ethoxy)ethyl)-*N*[5]-((*S*)-1-((2*S*,4*R*)-4-hydroxy-2-((4-(4-methylthiazol-5-yl)benzyl)carbamoyl)pyrrolidin-1-yl)-3,3-dimethyl-1-oxobutan-2-yl)glutaramide (X48)**

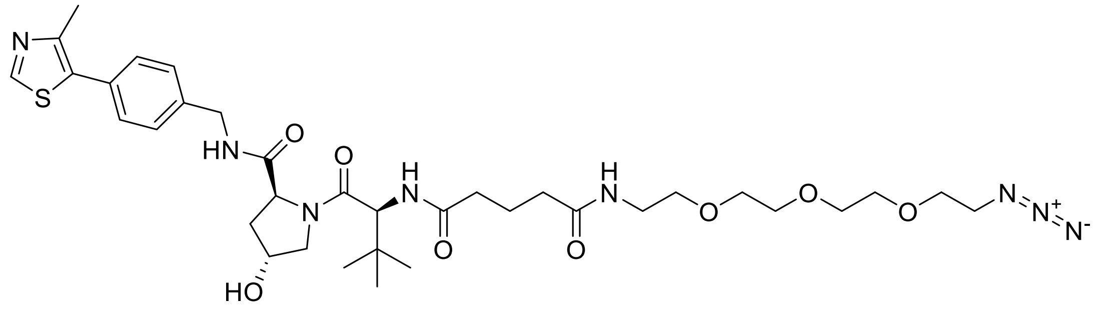

The title compound was prepared according to general procedure B, using **I42** (100 mg, 0.184 mmol) and 1-[2-(2-aminoethoxy)ethoxy]-2-(2-azidoethoxy)ethane (60 mg, 0.27 mmol), 1-methyl-1*H*-imidazole (51 µL, 0.64 mmol) and TCFH (61 mg, 0.22 mmol). The title compound was obtained as a colourless solid (128 mg, 94%). $^{1}$H NMR (500 MHz, $CDCl_3$) δ 8.68 (s, 1H), 7.41 – 7.31 (m, 5H), 6.82 (t, *J* = 5.6 Hz, 1H), 6.76 (d, *J* = 8.3 Hz, 1H), 4.74 (t, *J* = 8.2 Hz, 1H), 4.59 (dd, *J* = 15.0, 6.7 Hz, 1H), 4.50 (t, *J* = 3.1 Hz, 1H), 4.47 (d, *J* = 8.3 Hz, 1H), 4.33 (dd, *J* = 14.9, 5.1 Hz, 1H), 4.19 (d, *J* = 11.5 Hz, 1H), 3.67 – 3.61 (m, 12H), 3.59 – 3.50 (m, 4H), 3.48 – 3.42 (m, 1H), 3.38 (dd, *J* = 5.5, 4.5 Hz, 3H), 2.60 – 2.53 (m, 1H), 2.31 – 2.12 (m, 5H), 2.10 – 2.02 (m, 1H), 1.97 – 1.90 (m, 2H), 0.94 (s, 9H). $^{13}$C NMR (126 MHz, $CDCl_3$) δ 174.11, 172.98, 172.90, 170.66, 150.45, 148.65, 138.23, 131.70, 131.18, 129.68, 128.29, 70.78, 70.58, 70.20, 69.65, 58.53, 58.31, 56.90, 50.80, 43.40, 39.43, 36.04, 34.74, 34.50, 34.44, 29.86, 26.59, 22.32, 16.20. ESI-MS: *m/z* = 745.4 ([M+H]$^{+}$).

**Synthesis of 2-((*S*)-4-(4-chlorophenyl)-2,3,9-trimethyl-6*H*-thieno[3,2-*f*][1,2,4]triazolo[4,3-*a*][1,4]diazepin-6-yl)-*N*-(3-(1-(6-((2-(2,6-dioxopiperidin-3-yl)-1,3-dioxoisoindolin-5-yl)amino)hexyl)-1*H*-1,2,3-triazol-4-yl)propyl)acetamide (P1)**

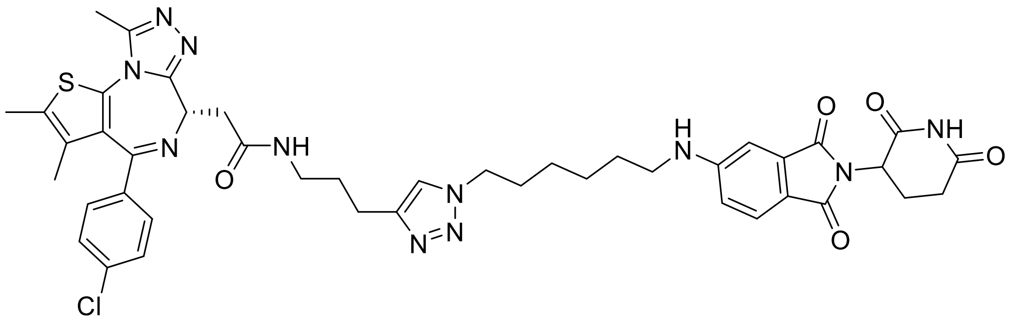

The title compound was prepared according to general procedure E, using **A3** (5.9 mg, 13 μmol), **X15** (5.0 mg, 13 μmol), $CuSO_4 \cdot 5H_2O$ (0.94 mg, 3.8 μmol) and sodium ascorbate (0.75 mg, 3.8 μmol). The title compound was obtained as a yellow solid (9.1 mg, 84%). $^1H$ NMR (400 MHz, DMSO-$d_6$) δ 11.05 (s, 1H), 8.27 (t, *J* = 5.7 Hz, 1H), 7.86 (s, 1H), 7.55 (d, *J* = 8.3 Hz, 1H), 7.50 – 7.33 (m, 4H), 7.07 (t, *J* = 5.0 Hz, 1H), 6.93 (s, 1H), 6.82 (dd, *J* = 8.5, 2.1 Hz, 1H), 5.02 (dd, *J* = 12.9, 5.4 Hz, 1H), 4.51 (dd, *J* = 8.4, 5.8 Hz, 1H), 4.30 (t, *J* = 7.0 Hz, 2H), 3.31 – 3.14 (m, 4H), 3.15 – 2.96 (m, 3H), 2.96 – 2.78 (m, 1H), 2.66 (t, *J* = 7.6 Hz, 2H), 2.59 (s, 3H), 2.56 – 2.53 (m, 1H), 2.39 (s, 3H), 2.03 – 1.95 (m, 1H), 1.86 – 1.70 (m, 4H), 1.59 (s, 3H), 1.56 – 1.48 (m, 2H), 1.44 – 1.33 (m, 2H), 1.31 – 1.20 (m, 2H). $^{13}C$ NMR (101 MHz, DMSO-$d_6$) δ 172.83, 170.18, 169.53, 167.70, 167.16, 163.06, 154.44, 146.40, 136.76, 135.19, 134.20, 132.28, 130.69, 130.12, 129.81, 129.56, 128.44, 125.10, 121.75, 115.81, 53.92, 49.11, 48.61, 42.35, 38.03, 37.67, 30.98, 29.70, 29.08, 28.01, 25.88, 25.62, 22.59, 22.23, 14.02, 12.66, 11.30. ESI-MS: *m/z* = 864.25 ([M+H]$^+$).

**Synthesis of 2-((*S*)-4-(4-chlorophenyl)-2,3,9-trimethyl-6*H*-thieno[3,2-*f*][1,2,4]triazolo[4,3-*a*][1,4]diazepin-6-yl)-*N*-((1-(6-((2-(2,6-dioxopiperidin-3-yl)-1,3-dioxoisoindolin-5-yl)amino)hexyl)-1*H*-1,2,3-triazol-4-yl)methyl)acetamide (P2)**

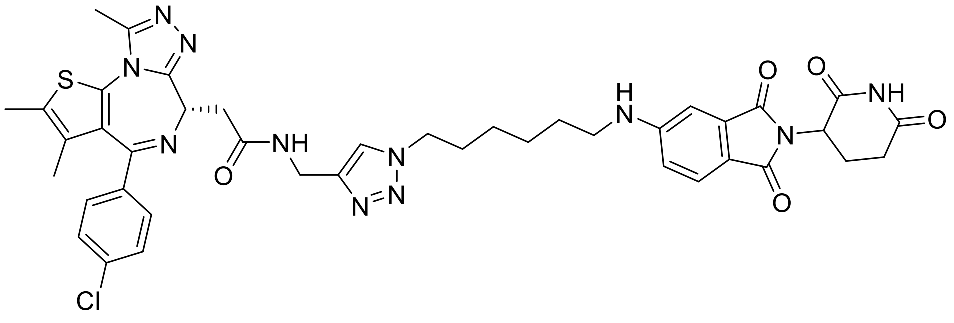

The title compound was prepared according to general procedure E, using **A1** (5.5 mg, 13 μmol), **X15** (5.0 mg, 13 μmol), $CuSO_4{\cdot}5H_2O$ (0.94 mg, 3.8 μmol) and sodium ascorbate (0.75 mg, 3.8 μmol). The title compound was obtained as a yellow solid (10.4 mg, 99%). $^1$H NMR (400 MHz, DMSO-$d_6$) δ 1.05 (s, 1H), 8.73 (t, *J* = 5.7 Hz, 1H), 7.94 (s, 1H), 7.55 (d, *J* = 8.4 Hz, 1H), 7.46 (d, *J* = 8.5 Hz, 2H), 7.37 (d, *J* = 8.6 Hz, 2H), 7.07 (t, *J* = 5.3 Hz, 1H), 6.93 (d, *J* = 2.1 Hz, 1H), 6.82 (dd, *J* = 8.3, 1.9 Hz, 1H), 5.02 (dd, *J* = 12.9, 5.4 Hz, 1H), 4.52 (t, *J* = 7.1 Hz, 1H), 4.35 (d, *J* = 5.6 Hz, 2H), 4.31 (t, *J* = 7.1 Hz, 2H), 3.27 (t, *J* = 7.3 Hz, 2H), 3.12 (q, *J* = 6.2 Hz, 2H), 2.94 – 2.76 (m, 1H), 2.59 (s, 3H), 2.54 (s, 2H), 2.39 (s, 3H), 2.02 – 1.94 (m, 1H), 1.79 (quin, *J* = 7.4 Hz, 2H), 1.61 (s, 3H), 1.53 (quin, *J* = 6.9 Hz, 2H), 1.46 – 1.32 (m, 2H), 1.31 – 1.20 (m, 2H). $^{13}$C NMR (101 MHz, DMSO-$d_6$) δ $^{13}$C NMR (101 MHz, DMSO) δ 172.83, 170.19, 169.62, 167.71, 167.16, 163.06, 155.08, 154.44, 149.87, 145.00, 136.72, 135.22, 134.21, 132.29, 130.68, 130.21, 129.84, 129.52, 128.42, 125.11, 122.71, 115.82, 53.90, 49.24, 48.61, 42.37, 40.43, 37.54, 34.29, 30.98, 29.68, 28.01, 25.89, 25.62, 22.24, 14.06, 12.66, 11.30. ESI-MS: *m/z* = 836.25 ([M+H]$^+$).

**Synthesis of 2-((*S*)-4-(4-chlorophenyl)-2,3,9-trimethyl-6*H*-thieno[3,2-*f*][1,2,4]triazolo[4,3-*a*][1,4]diazepin-6-yl)-*N*-(3-(1-(6-((2-(2,6-dioxopiperidin-3-yl)-1,3-dioxoisoindolin-4-yl)amino)hexyl)-1*H*-1,2,3-triazol-4-yl)propyl)acetamide (P3)**

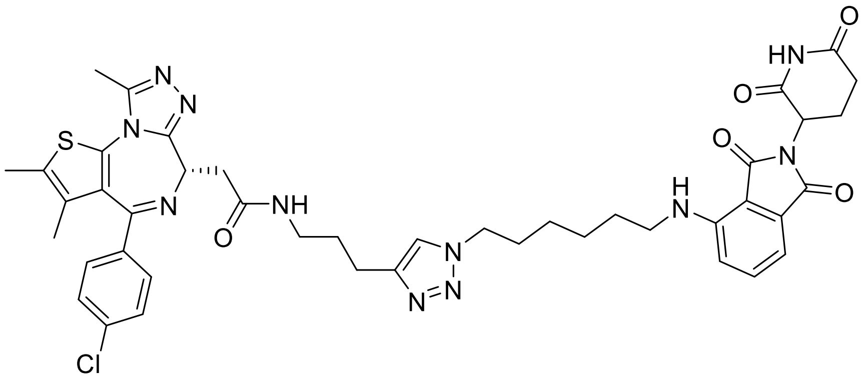

The title compound was prepared according to general procedure E, using **A3** (5.9 mg, 13 μmol), **X6** (5.0 mg, 13 μmol), $CuSO_4{\cdot}5H_2O$ (0.94 mg, 3.8 μmol) and sodium ascorbate (0.75 mg, 3.8 μmol). The title compound was obtained as a yellow solid (6.2 mg, 57%). $^1$H NMR (400 MHz, DMSO-$d_6$) δ 11.08 (s, 1H), 8.26 (t, *J* = 5.7 Hz, 1H), 7.85 (s, 1H), 7.56 (dd, *J* = 8.6, 7.1 Hz, 1H), 7.50 – 7.36 (m, 4H), 7.06 (d, *J* = 8.6 Hz, 1H), 7.00 (d, *J* = 7.0 Hz, 1H), 6.51

(t, *J* = 5.9 Hz, 1H), 5.04 (dd, *J* = 12.8, 5.4 Hz, 1H), 4.51 (dd, *J* = 8.5, 5.8 Hz, 1H), 4.29 (t, *J* = 7.0 Hz, 2H), 3.29 – 3.14 (m, 5H), 3.14 – 3.06 (m, 1H), 2.93 – 2.77 (m, 1H), 2.66 (t, *J* = 7.7 Hz, 2H), 2.59 (s, 3H), 2.55 – 2.51 (m, 2H), 2.39 (s, 3H), 2.05 – 1.98 (m, 1H), 1.84 – 1.74 (m, 4H), 1.59 (s, 3H), 1.53 (quin, *J* = 7.3 Hz, 2H), 1.35 (quin, *J* = 7.3 Hz, 2H), 1.25 (quin, *J* = 7.2 Hz, 2H). $^{13}$C NMR (101 MHz, DMSO-$d_6$) δ 172.81, 170.10, 169.53, 168.94, 167.31, 163.05, 155.13, 149.82, 146.39, 136.76, 136.27, 135.18, 132.27, 130.69, 130.12, 129.81, 129.56, 128.43, 121.73, 117.16, 110.38, 109.02, 53.92, 49.10, 48.54, 41.72, 38.04, 37.68, 30.97, 29.65, 29.09, 28.46, 25.67, 25.58, 22.59, 22.15, 14.01, 12.65, 11.29. ESI-MS: *m/z* = 864.20 ([M+H]$^+$).

**Synthesis of (2*S*,4*R*)-1-((*S*)-2-(4-(4-(2-(2-((*S*)-4-(4-chlorophenyl)-2,3,9-trimethyl-6*H*-thieno[3,2-*f*][1,2,4]triazolo[4,3-*a*][1,4]diazepin-6-yl)acetamido)ethyl)-1*H*-1,2,3-triazol-1-yl)benzamido)-3,3-dimethylbutanoyl)-4-hydroxy-*N*-(4-(4-methylthiazol-5-yl)benzyl)pyrrolidine-2-carboxamide (P4)**

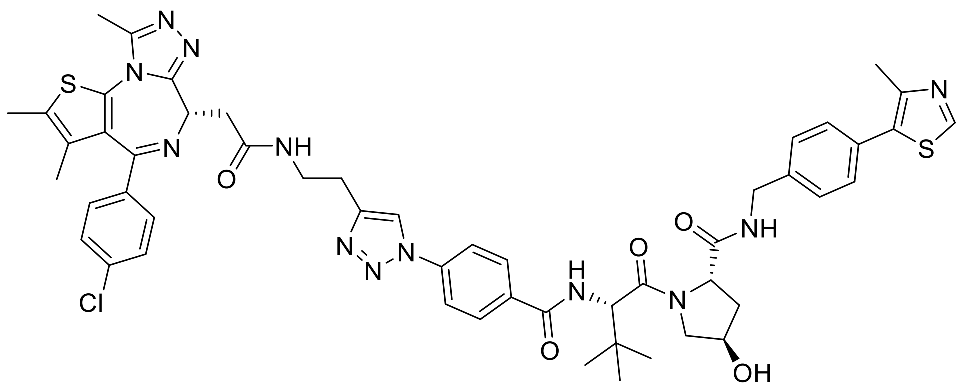

The title compound was prepared according to general procedure E, using **A2** (3.9 mg, 8.7 µmol), **X37** (5.0 mg, 8.7 µmol), $CuSO_4 \cdot 5H_2O$ (0.65 mg, 2.6 µmol) and sodium ascorbate (0.52 mg, 2.6 µmol). The title compound was obtained as a colourless solid (4.5 mg, 50%). $^1$H NMR (400 MHz, DMSO-$d_6$) δ 8.98 (s, 1H), 8.79 (s, 1H), 8.58 (t, *J* = 6.1 Hz, 1H), 8.39 (t, *J* = 5.7 Hz, 1H), 8.21 (d, *J* = 9.0 Hz, 1H), 8.09 (d, *J* = 8.8 Hz, 2H), 7.99 (d, *J* = 8.8 Hz, 2H), 7.47 – 7.35 (m, 7H), 5.16 (d, *J* = 3.6 Hz, 1H), 4.80 (d, *J* = 9.1 Hz, 1H), 4.52 (t, *J* = 7.2 Hz, 1H), 4.46 (q, *J* = 7.2, 6.2 Hz, 1H), 4.43 – 4.37 (m, 1H), 4.24 (dd, *J* = 15.8, 5.6 Hz, 1H), 3.74 (d, *J* = 3.0 Hz, 2H), 3.52 – 3.45 (m, 2H), 3.44 – 3.40 (m, 2H), 3.28 – 3.23 (m, 2H), 2.91 (t, *J* = 6.9 Hz, 2H), 2.58 (s, 3H), 2.45 (s, 3H), 2.40 (s, 3H), 2.11 – 2.00 (m, 1H), 1.97 – 1.90

(m, 1H), 1.60 (s, 3H), 1.05 (s, 9H). $^{13}C$ NMR (101 MHz, DMSO-$d_6$) δ 171.92, 169.72, 169.37, 165.53, 163.12, 155.11, 151.48, 149.91, 147.75, 145.94, 139.49, 138.63, 136.76, 135.21, 133.57, 132.24, 131.18, 130.72, 130.12, 129.88, 129.68, 129.58, 129.55, 128.70, 128.46, 127.47, 120.99, 119.09, 68.93, 58.84, 57.49, 56.46, 53.86, 41.69, 39.52, 38.23, 37.96, 37.71, 35.59, 26.55, 25.51, 15.95, 14.02, 12.68, 11.31. ESI-MS: *m/z* = 514.30 ($[M+2H]^{+2}$).

**Synthesis of 2-((*S*)-4-(4-chlorophenyl)-2,3,9-trimethyl-6*H*-thieno[3,2-*f*][1,2,4]triazolo[4,3-*a*][1,4]diazepin-6-yl)-*N*-((1-(6-((2-(2,6-dioxopiperidin-3-yl)-1,3-dioxoisoindolin-4-yl)amino)hexyl)-1*H*-1,2,3-triazol-4-yl)methyl)acetamide (P5)**

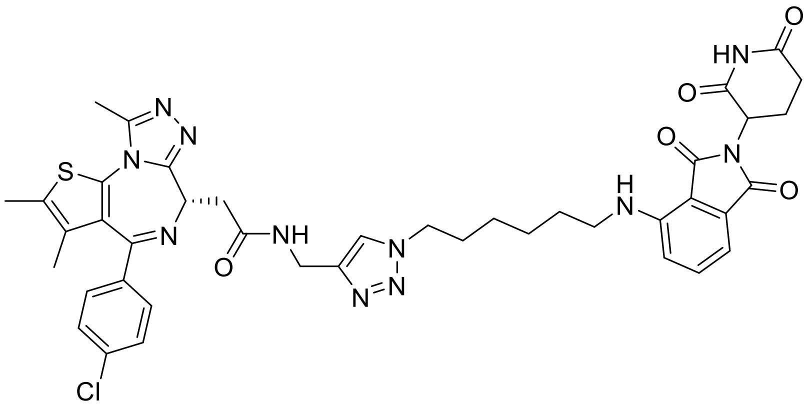


The title compound was prepared according to general procedure E, using **A1** (5.5 mg, 13 µmol), **X6** (5.0 mg, 13 µmol), $CuSO_4·5H_2O$ (0.94 mg, 3.8 µmol) and sodium ascorbate (0.75 mg, 3.8 µmol). The title compound was obtained as a yellow solid (10.1 mg, 96%). $^1H$ NMR (400 MHz, DMSO-$d_6$) δ 11.08 (s, 1H), 8.72 (t, *J* = 5.8 Hz, 1H), 7.94 (s, 1H), 7.56 (t, *J* = 7.8 Hz, 1H), 7.45 (d, *J* = 8.2 Hz, 2H), 7.37 (d, *J* = 8.2 Hz, 2H), 7.06 (d, *J* = 8.6 Hz, 1H), 7.00 (d, *J* = 7.0 Hz, 1H), 6.51 (t, *J* = 6.0 Hz, 1H), 5.04 (dd, *J* = 12.8, 5.4 Hz, 1H), 4.52 (dd, *J* = 8.1, 6.3 Hz, 1H), 4.35 (d, *J* = 5.7 Hz, 2H), 4.30 (t, *J* = 7.2 Hz, 2H), 3.29 – 3.22 (m, 4H), 2.94 – 2.81 (m, 1H), 2.59 (s, 3H), 2.55 – 2.52 (m, 2H), 2.39 (s, 3H), 2.06 – 1.98 (m, 1H), 1.78 (quin, *J* = 6.9 Hz, 2H), 1.61 (s, 3H), 1.54 (quin, *J* = 7.2 Hz, 2H), 1.37 – 1.33 (m, 2H), 1.31 – 1.23 (m, 2H). $^{13}C$ NMR (101 MHz, DMSO-$d_6$) δ 172.80, 170.09, 169.59, 168.94, 167.30, 163.03, 155.07, 149.86, 146.40, 144.97, 136.70, 136.27, 135.21, 132.19, 130.67, 130.20, 129.83, 129.51, 128.41, 122.70, 117.16, 110.37, 109.02, 53.90, 49.22, 48.53, 41.73, 37.54, 34.28, 30.97, 29.64, 28.46, 25.68, 25.59, 22.15, 14.05, 12.65, 11.29. ESI-MS: *m/z* = 836.20 ($[M+H]^+$).

**Synthesis of 4-(4-(2-(4-(2-(2-((*S*)-4-(4-chlorophenyl)-2,3,9-trimethyl-6*H*-thieno[3,2-*f*][1,2,4]triazolo[4,3-*a*][1,4]diazepin-6-yl)acetamido)ethyl)-1*H*-1,2,3-triazol-1-yl)acetyl)piperazin-1-yl)-*N*-(2,6-dioxopiperidin-3-yl)-2-methoxybenzamide (P6)**

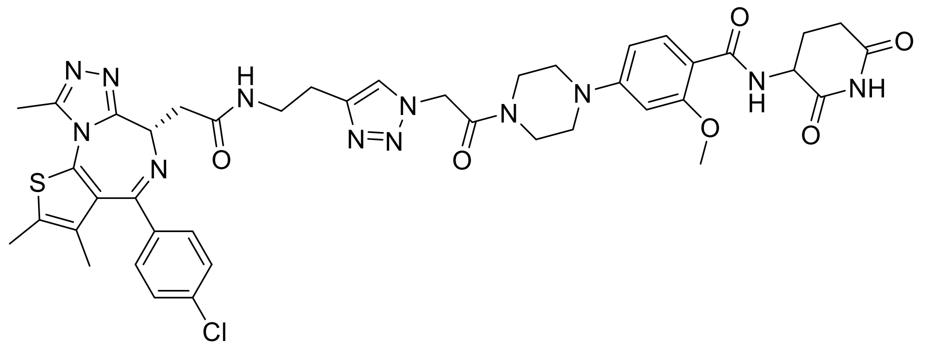

The title compound was prepared according to general procedure E, using **A2** (5.3 mg, 12 μmol), **X35** (5.0 mg, 12 μmol), $CuSO_4 \cdot 5H_2O$ (0.87 mg, 3.5 μmol) and sodium ascorbate (0.69 mg, 3.5 μmol). The title compound was obtained as a colourless solid (5.7 mg, 56%). $^1H$ NMR (400 MHz, DMSO-$d_6$) δ 10.87 (s, 1H), 8.45 (d, *J* = 7.0 Hz, 1H), 8.36 (t, *J* = 5.5 Hz, 1H), 7.86 (s, 1H), 7.81 (d, *J* = 8.8 Hz, 1H), 7.49 (d, *J* = 8.3 Hz, 2H), 7.42 (d, *J* = 8.2 Hz, 2H), 6.63 (d, *J* = 9.0 Hz, 1H), 6.59 (s, 1H), 5.51 (s, 2H), 4.71 (dt, *J* = 12.5, 6.2 Hz, 1H), 4.52 (t, *J* = 7.1 Hz, 1H), 3.94 (s, 3H), 3.69 (t, *J* = 5.2 Hz, 2H), 3.62 (t, *J* = 5.5 Hz, 2H), 3.45 (s, 2H), 3.41 – 3.36 (m, 3H), 3.27 – 3.21 (m, 2H), 2.84 (t, *J* = 7.3 Hz, 2H), 2.76 (ddd, *J* = 18.3, 13.5, 5.6 Hz, 1H), 2.60 (s, 3H), 2.56 – 2.51 (m, 2H), 2.41 (s, 3H), 2.17 – 2.01 (m, 2H), 1.63 (s, 3H). $^{13}C$ NMR (101 MHz, DMSO-$d_6$) δ 172.96, 172.66, 169.55, 164.58, 164.27, 163.07, 158.90, 155.13, 154.04, 149.88, 144.08, 136.77, 135.21, 132.41, 132.24, 130.71, 130.16, 129.86, 129.59, 128.49, 124.06, 111.00, 106.73, 97.82, 55.93, 53.82, 50.56, 50.09, 46.96, 46.70, 43.75, 41.12, 38.40, 37.61, 31.24, 31.04, 28.35, 25.55, 24.37, 22.09, 14.05, 12.68, 11.31. ESI-MS: *m/z* = 881.25 ($[M+H]^+$).

**Synthesis of (2*S*,4*R*)-1-((*S*)-2-(4-(4-((4-(2-((*S*)-4-(4-chlorophenyl)-2,3,9-trimethyl-6*H*-thieno[3,2-*f*][1,2,4]triazolo[4,3-*a*][1,4]diazepin-6-yl)acetyl)piperazin-1-yl)methyl)-1*H*-1,2,3-triazol-1-yl)benzamido)-3,3-dimethylbutanoyl)-4-hydroxy-*N*-(4-(4-methylthiazol-5-yl)benzyl)pyrrolidine-2-carboxamide (P7)**

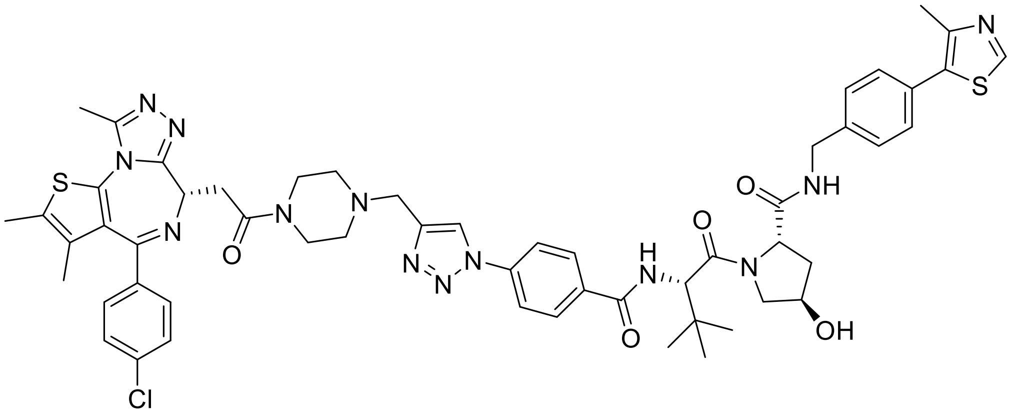

The title compound was prepared according to general procedure E, using **A4** (4.4 mg, 8.7 µmol), **X37** (5.0 mg, 8.7 µmol), $CuSO_4 \cdot 5H_2O$ (0.65 mg, 2.6 µmol) and sodium ascorbate (0.52 mg, 2.6 µmol). The title compound was obtained as a colourless solid (5.0 mg, 53%). $^1H$ NMR (400 MHz, DMSO-$d_6$) δ 8.98 (s, 1H), 8.87 (s, 1H), 8.58 (t, *J* = 6.1 Hz, 1H), 8.25 (d, *J* = 9.0 Hz, 1H), 8.11 (d, *J* = 8.7 Hz, 2H), 8.03 (d, *J* = 8.7 Hz, 2H), 7.48 (d, *J* = 8.7 Hz, 2H), 7.45 – 7.38 (m, 6H), 5.16 (d, *J* = 3.6 Hz, 1H), 4.81 (d, *J* = 9.1 Hz, 1H), 4.56 (t, *J* = 6.7 Hz, 1H), 4.46 (q, *J* = 7.5, 6.9 Hz, 1H), 4.40 (dd, *J* = 14.4, 5.0 Hz, 2H), 4.24 (dd, *J* = 15.9, 5.6 Hz, 1H), 3.75 (d, *J* = 3.0 Hz, 3H), 3.68 (s, 2H), 3.60 (dd, *J* = 16.4, 7.2 Hz, 1H), 3.55 – 3.44 (m, 1H), 3.40 (dd, *J* = 16.3, 6.3 Hz, 1H), 2.59 (s, 3H), 2.55 – 2.51 (m, 2H), 2.48 – 2.45 (m, 4H), 2.45 (s, 3H), 2.41 (s, 3H), 2.13 – 2.00 (m, 1H), 1.98 – 1.89 (m, 1H), 1.62 (s, 3H), 1.06 (s, 9H). $^{13}C$ NMR (101 MHz, DMSO-$d_6$) δ 171.91, 169.36, 168.15, 165.54, 162.87, 155.24, 151.46, 149.76, 147.74, 139.49, 138.52, 136.76, 135.19, 132.19, 131.16, 130.67, 130.14, 129.88, 129.54, 128.69, 128.47, 127.47, 119.28, 68.92, 58.83, 57.51, 56.45, 54.15, 41.68, 37.96, 35.57, 34.71, 26.56, 15.94, 14.00, 12.67, 11.26. ESI-MS: *m/z* = 541.80 ($[M+2H]^{+2}$).

**Synthesis of (*S*)-2-(4-(4-chlorophenyl)-2,3,9-trimethyl-6*H*-thieno[3,2-*f*][1,2,4]triazolo[4,3-*a*][1,4]diazepin-6-yl)-*N*-(3-(1-(4-(4-(2-(3-(2,4-dioxotetrahydropyrimidin-1(2*H*)-yl)-2-methylphenoxy)acetyl)piperazine-1-carbonyl)phenyl)-1*H*-1,2,3-triazol-4-yl)propyl)acetamide (P8)**

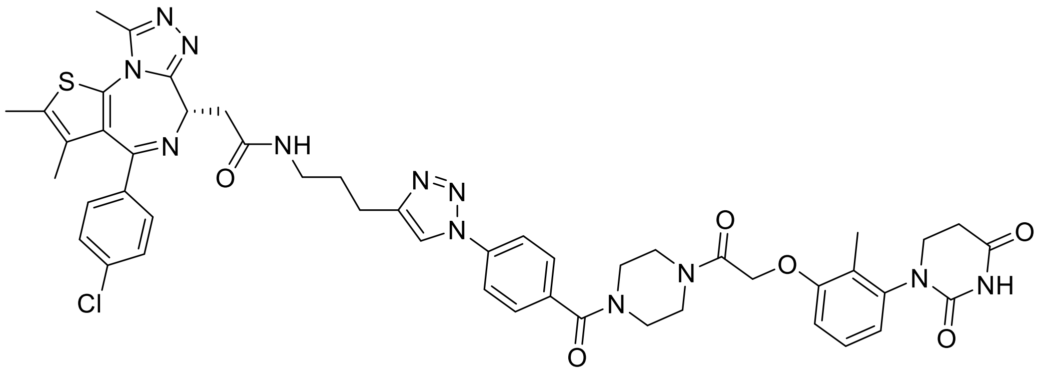

The title compound was prepared according to general procedure E, using **A3** (4.7 mg, 10 µmol), **X24** (5.0 mg, 10 µmol), $CuSO_4 \cdot 5H_2O$ (0.76 mg, 3.1 µmol) and sodium ascorbate (0.61 mg, 3.1 µmol). The title compound was obtained as a colourless solid (9.3 mg, 95%). $^1$H NMR (400 MHz, DMSO-$d_6$) δ 10.32 (s, 1H), 8.65 (s, 1H), 8.32 (t, *J* = 5.7 Hz, 1H), 7.97 (d, *J* = 8.7 Hz, 2H), 7.66 (d, *J* = 8.6 Hz, 2H), 7.48 – 7.37 (m, 4H), 7.17 (t, *J* = 8.1 Hz, 1H), 6.97 – 6.78 (m, 2H), 4.91 (s, 2H), 4.52 (dd, *J* = 8.3, 6.0 Hz, 1H), 3.82 – 3.72 (m, 1H), 3.69 – 3.38 (m, 8H), 3.32 – 3.17 (m, 5H), 2.83 – 2.74 (m, 3H), 2.67 (dt, *J* = 16.6, 5.5 Hz, 1H), 2.59 (s, 3H), 2.40 (s, 3H), 2.05 (s, 3H), 1.88 (quin, *J* = 7.1 Hz, 2H), 1.60 (s, 3H). $^{13}$C NMR (101 MHz, DMSO-$d_6$) δ $^{13}$C NMR (126 MHz, DMSO) δ 170.76, 169.60, 168.28, 166.16, 163.09, 156.56, 155.15, 151.76, 149.86, 147.96, 141.71, 137.40, 135.37, 135.20, 132.28, 130.72, 130.14, 129.85, 129.59, 128.84, 128.44, 126.52, 124.20, 120.32, 119.67, 110.71, 66.50, 53.93, 44.69, 40.43, 38.02, 37.70, 31.08, 28.85, 22.57, 14.04, 12.68, 11.30, 10.78. ESI-MS: *m/z* = 957.15 ([M+H]$^+$).

**Cell Viability Assay**

The effect of the compounds on cell viability was determined using the CellTiter-Glo® 2.0 Cell Viability Assay (Promega: G9241) following manufacturer's protocol. 10 µL of HEK293T cells were seeded at a cell density of $2 \cdot 10^5$ cells/mL into individual wells of a white 384-well plate (Greiner: 781207) and the cells were allowed to equilibrate for 1 h at 37 °C and 5% $CO_2$. After equilibration, the compounds were titrated at various concentrations using an Echo acoustic dispenser (Labcyte) and the cells were incubated for 24 h at 37 °C and 5% $CO_2$. Equal volume of CellTiter-Glo® 2.0 reagent (10 µL) was added to each well and cells were incubated for 10 min at rt. Filtered luminescence was measured on a PHERAstar plate reader (BMG Labtech) and data was evaluated using

GraphPad Prism 9 software employing a normalized curve it with the following equation: $Y = Bottom + (Top-Bottom)/(1 + 10^{(LogEC50-X)*HillSlope})$.

**HiBiT endpoint detection for BRD4 degradation**

Endogenously BRD4 HiBiT-tagged HEK293T (HEK293T$^{BRD4-HiBiT}$) cells were obtained as a kind gift from Promega Corp. To measure degradation, 10 µl of a total concentration of $2.5x10^5$ cells/ml in DMEM medium were seeded into white small volume 384 well plates (Greiner, 784075) and allowed to settle overnight. Subsequently, the PROTACs were added to the seeded cells, using an Echo acoustic dispenser (Labcyte) and the plate was incubated for the indicated time at 37°C and 5 % $CO_2$. After incubation, HiBiT Lytic detection reagent was prepared by dilution of LgBiT protein (1:100) and lytic substrate (1:50) in Lytic detection buffer (Promega, N3040). For detection, 10 µl of the prepared mix was added to the treated cells and incubated for 10 minutes at rt. Readout was carried out in a PHERAStar FSX plate reader (BMG Labtech) using the LUM plus optical module. Degradation data were then plotted with GraphPad Prism 9 software using a normalized 3-parameter curve fit with the following equation: Y = 100/(1 + 10^(X-LogIC50))

**HiBiT endpoint detection for WDR5 and AURKA degradation**

The mutated nanoluciferase (K55R, K77R, K80R, K91R, K125R, K126R and K138R), referred to as Nluc (Kless), was cloned by PCR amplification of the vector pRRL-Puro-C-term-Luc-FKBP12 (Kless) using the forward primer TACGCGTCATATGACTAGTGGGA and the reverse primer CGGATCCTCACGCCAGAATGCGTTCGCA. The amplified product was inserted into the pRRL-PGK-Puro entry vector using MluI/BamHI restriction sites to yield pRRL-PGK-Puro-Nluc (Kless). AURORA-A-Nluc (Kless) was cloned by amplification of the vector pRRL-PGK-Hygro-HiBiT-AURORA-A, containing full-length AURORA-A, via PCR using the forward and reverse primers CCACCGGTATGGACCGATCT and GTACGCGTAGACTGTTTGCTAGCTGATTCTTTGTTTTG, respectively. pRRL-PGK-Puro-AURORA-A-Nluc (Kless) was generated through insertion of the PCR product into the pRRL-PGK-Puro-Nluc (Kless) entry vector using AgeI/MluI restriction sites. WDR5-Nluc (Kless) was cloned by PCR amplification of the vector pRRL-PGK-Hygro-HiBiT-WDR5 containing full-length WDR5 using the forward primer CCACCGGTATGGCGACGG and the reverse primer GTACGCGTGCAGTCACTCTTCCACAGTTTAATTGTT. The PCR product was inserted into the pRRL-PGK-Puro-Nluc (Kless) entry vector using AgeI/MluI restriction sites to obtain pRRL-PGK-Puro-WDR5 Nluc (Kless). The PCR reactions were performed using the Phusion High-Fidelity DNA Polymerase (Thermo Fisher Scientific).

Stable MV4-11$^{AURORA\text{-}A\text{-}NLuc(Kless)}$ and MV4-11$^{WDR5\text{-}NLuc(Kless)}$ cells were generated using lentiviral infection. The lentivirus was produced by transfecting HEK293T cells with the plasmids psPAX2, pMD2.G, and pRRL-PGK-Puro-AURORA-A-Nluc (Kless) or the pRRL-PGK-Puro-WDR5-Nluc (Kless) using polyethyleneimine (PEI, Sigma). The virus-containing supernatant was filtered and used to infect MV4-11 cells, which were subsequently selected with puromycin (InvivoGen) at a final concentration of 2 µg/mL after 48 h of infection.

MV4-11$^{AURORA\text{-}A\text{-}HiBiT}$ or MV4-11$^{WDR5\text{-}HiBiT}$ cells were seeded in a black 96-well cell culture microplate (Greiner) and treated for 6 h with the test compounds at concentrations of 50 nM, 200 nM or 1 µM. JB301 and AD122 were used at concentrations of 150 nM and 1 µM, respectively. The control cells were treated with dimethylsulfoxide (DMSO; Roth). The Nano-Glo® HiBiT Lytic Detection System (Promega) was used for the assay. The luminescence was measured using the Infinite M Plex, multimode microplate reader (TECAN).

**Nanoluciferase live cell measurement for WDR5 and AURKA degradation**

MV4-11$^{AURORA\text{-}A\text{-}Nluc(Kless)}$ or MV4-11$^{WDR5\text{-}Nluc(Kless)}$ cells were seeded in a black 96-well cell culture microplate (Greiner) at a density of 200 000 cells/mL in Opti-MEM™ reduced serum medium, without phenol red (Thermo Fisher Scientific), supplemented with 10 % FBS (Capricorn Scientific), 15 mM HEPES pH 7.2. The Nano-Glo® Endurazine™ live cell substrate (Promega) was diluted and added to the cells according to the manufacturer´s instructions. Prior to the addition of the test compounds, the cells were incubated for 3 h at 37 °C in 5 % $CO_2$. Finally, the cells were treated with 1 µM of the PROTACs and the kinetic measurement was done at 15 min intervals during 12 h at 37 °C. JB301 was used at a concentration of 150 nM and AD122 at 1 µM. DMSO served as a vehicle control. The luminescence was measured using the Infinite M Plex, multimode microplate reader (TECAN).

**HiBiT endpoint detection for sEH degradation**

To generate the HeLa-sEH-HiBiT cell line, HeLa cells were stably transfected with the construct hsEH_aa1-aa555_Linker-HiBiT_pSB-hPGK using the Sleeping Beauty transposon system (10.1002/biot.201400821). This construct, assembled via Gibson cloning, encodes the human soluble epoxide hydrolase (sEH; amino acids 1–555) fused at its C-terminus to the HiBiT peptide, driven by the human PGK promoter. Initially, an intermediate construct, hsEH_aa1-aa555_Linker-HiBiT_pSBtet, was created by inserting the hsEH_aa1-aa555_Linker-HiBiT sequence into the pSBtet-bla vector. This version enables stable expression of the fusion protein under the control of a doxycycline-inducible tetOn promoter. Subsequently, the tetOn promoter was replaced with the constitutive hPGK promoter to produce the final construct, hsEH_aa1-aa555_Linker-HiBiT_pSB-hPGK. Primers for DNA amplification were obtained from Eurofins. All PCR reactions were carried out using Q5® High-Fidelity DNA Polymerase following the manufacturer's instructions. Each PCR product underwent digestion with DpnI at 37 °C for 1 h, followed by enzyme inactivation at 80 °C for 20 min, and subsequent purification using the GeneJET PCR Purification Kit according to the supplier's protocol. The hsEH-Linker-HiBiT insert was generated through a two-step process. In PCR 1, a dsDNA fragment encoding the Linker-HiBiT sequence was amplified from a plasmid containing the codon-optimized CDS for the linker (SSGNSGGSSG) and HiBiT (VSGWRLFKKIS) sequences for *Homo sapiens*. A forward primer (5'-CCACCGGTGGTCTCAAAGATGAGCAGCGGCAACAGC-3') and a reverse primer (5'-TCGATGGAAGCTTGGCCTGACAGGCCTCAGCTGATCTTCTTGAACAGCCG-3') were used to add a 21 bp overlap with the C-terminus of the hsEH CDS at the 5' end, and a 29 bp region at the 3' end containing a TGA stop codon and overlap with the pSBtet-Bla vector backbone near the SfiI site downstream of the MCS. The reaction conditions were: initial denaturation at 98 °C for 1 min; 25 cycles of 98 °C for 20 s, 66 °C for 20 s, and 72 °C for 30 s; with a final extension at 72 °C for 5 min. To generate the final hsEH_aa1-aa555_Linker-HiBiT insert, PCR 2 was performed as a fusion PCR using both the PCR 1 product and the published sEH construct from Hahn et al. (10.1002/cmdc.201100433) as templates. The same reverse primer as in PCR 1 and a forward primer (5'-TACCCTCGAAAGGCCTCTGAGGCCACCATGACGCTGCGCGC-3') were used. This forward primer introduced a 27 bp overlap with the 5' SfiI site region of pSBtet-Bla to facilitate Gibson Assembly. Reaction conditions were: 98 °C for 1 min; 25 cycles of 98 °C for 40 s, 65 °C for 20 s, and 72 °C for 1 min 30 s; with a final extension at 72 °C for 5 min. For PCR 3, the entire pSBtet-Bla vector backbone was amplified using the forward primer (5'-TGAGGCCTGTCAGGCCAAGCTTCCATCGA-3') and reverse primer (5'-CATGGTGGCCTCAGAGGCCTTTCGAGGGTA-3'). Cycling conditions were: 98 °C for 1 min; 25 cycles of 98 °C for 20 s, 66 °C for 20 s, and 72 °C for 5 min; with a final extension at 72 °C for 8 min. The final hsEH_aa1-aa555_Linker-HiBiT_pSBtet construct was assembled using the NEBuilder® HiFi DNA Assembly Cloning Kit as per the manufacturer's instructions. The 5 µl assembly reaction contained approximately 40 ng of insert DNA (PCR 2) and 52 ng of vector backbone (PCR 3). For the final construct hsEH_aa1-aa555_Linker-HiBiT_pSB-hPGK, the hPGK promoter fragment was generated in PCR 4 using pLKO.1-puro-shNM as template with the forward primer (5'-GGTCCGCTATCTAGACGAGTAGCAGAGATCCACTTTGGCC-3') and reverse primer (5'-GGCAAAAGAGTTGGAATTGGCCCTGGGGAGAGAGGTCGG-3'). Reaction conditions: 98 °C for 30 s; 25 cycles of 98 °C for 10 s, 62 °C for 30 s, and 72 °C for 30 s; final extension at 72 °C for 4 min. In PCR 5, the intermediate construct (hsEH_aa1-aa555_Linker-HiBiT_pSBtet) was amplified with forward primer (5'-GCCAATTCCAACTCTTTTGCCTTATACC-3') and reverse primer (5'-ACTCGTCTAGATAGCGGACC-3') to generate a linearized plasmid excluding the tet-On promoter region. Conditions were: 98 °C for 2 min; 30 cycles of 98 °C for 40 s, 62 °C for 20 s, and 72 °C for 5 min; with a final extension at 72 °C for 8 min. The final construct was assembled using the NEBuilder® HiFi DNA Assembly Cloning Kit as recommended, with 5 µl assembly reactions containing ~13 ng of the hPGK promoter fragment (PCR 4) and ~66 ng of the linearized vector (PCR 5). HeLa cells were seeded two days prior to transfection in 6-well plates at a density of $4 \times 10^5$ cells per well in 3 mL of DMEM (1X) medium with phenol, supplemented with 10% Corning® Fetal Bovine Serum, 100 U/mL penicillin and 100 µg/mL streptomycin, and 1 mM sodium pyruvate. This medium is hereafter referred to as DMEMsup. On the day of transfection, each well was washed with 2 mL PBS and then incubated with 1 mL Opti-MEM™ medium. Transfection was performed using the Lipofectamine™ 3000 protocol with minor adjustments. 500 µL of Opti-MEM™ was mixed with 4.2 µg of hsEH_aa1-aa555_Linker-HiBiT_pSB-hPGK plasmid, 0.2 µg of pSB100x transposase plasmid, and 8.8 µL of P3000™ reagent. Separately, 500 µL of Opti-MEM™ was combined with 4 µL of Lipofectamine™ 3000. The two mixtures were then combined and incubated at rt for 10 min before being added to a single well of the 6-well plate. Cells were incubated with the transfection mixture at 37 °C and 5% $CO_2$ for 4 h, after which the medium was replaced with 2 mL of DMEMsup. 24 h post-transfection, cells were transferred to a 75 $cm^2$ tissue culture flask and cultured in selection medium, consisting of 50 mL DMEMsup supplemented with 25 µL of a 10 mg/mL blasticidin stock solution (final concentration: 5 µg/mL). After 4 days under selection, cells were transferred to a 175 $cm^2$ flask and maintained in selection medium for an additional 10–14 days to establish stably transfected cell lines.

The degradation of sEH induced by PROTACs was evaluated using the previously established sEH-HiBiT lytic assay, following the published protocol (acs.jmedchem.5c00552). Briefly, HeLa cells stably expressing the sEH-HiBiT fusion protein (hereafter referred to as $HeLa^{sEH\text{-}HiBiT}$) were cultured in growth medium DMEMsup. Cells were maintained at 37 °C in a 5% $CO_2$ atmosphere. For assay setup, cells were harvested in growth medium, adjusted to a density of $4{\cdot}10^5$ cells/mL, and seeded into 384-well tissue culture plates at 50 µL per well (equivalent to 2000 cells per well) using a Multidrop Combi dispenser (Thermo Fisher Scientific). Plates were sealed with AeraSeal™ semipermeable film and incubated for 24 h at 37 °C, 5% $CO_2$. For compound screening, crude PROTACs were tested at final concentrations of 200 nM and 1 µM. Stock solutions were prepared in DMSO at 40 µM and 200 µM in a 96-deep-well plate, then diluted with growth medium to 2.2 µM and 11 µM (final DMSO concentration 5.5%). From these dilutions, 5 µL was added to each well in triplicate, yielding a final assay volume of 55 µL and a final DMSO concentration of 0.5%. Plates were centrifuged for 1 min at 300 rpm, resealed, and incubated for 6 or 18 h at 37 °C and 5% $CO_2$. Following treatment, cells were washed four times with DPBS using a HydroSpeed™ plate washer (Tecan), leaving 10 µL of residual volume in each well. Cell lysis was carried out by adding 1 µL of Mammalian Lysis Buffer, followed by centrifugation for 1 minute at 300 rpm and a 10-minute incubation at rt. In the meantime, the Nano-Glo substrate mix was freshly prepared using 6760 µL of Nano-Glo HiBiT Extracellular Buffer, 135 µL of Nano-Glo HiBiT Extracellular Substrate, and 68 µL of LgBiT Protein (all from the Nano-Glo® HiBiT Extracellular Detection System Kit, Promega, calculated for 616 wells). After lysis, 10 µL of the substrate mix was added to each well, followed by centrifugation (1 min at 300 rpm) and a 10-minute incubation at rt. Luminescence was then measured using a Spark Multimode Microplate Reader (Tecan). The average luminescence of each triplicate was normalized to the DMSO control and plotted against the respective azide using Prism 7.0 (GraphPad Software).
