## Supplemental Information for "Click. Screen. Degrade. A Miniaturized D2B Workflow for rapid PROTAC Discovery"

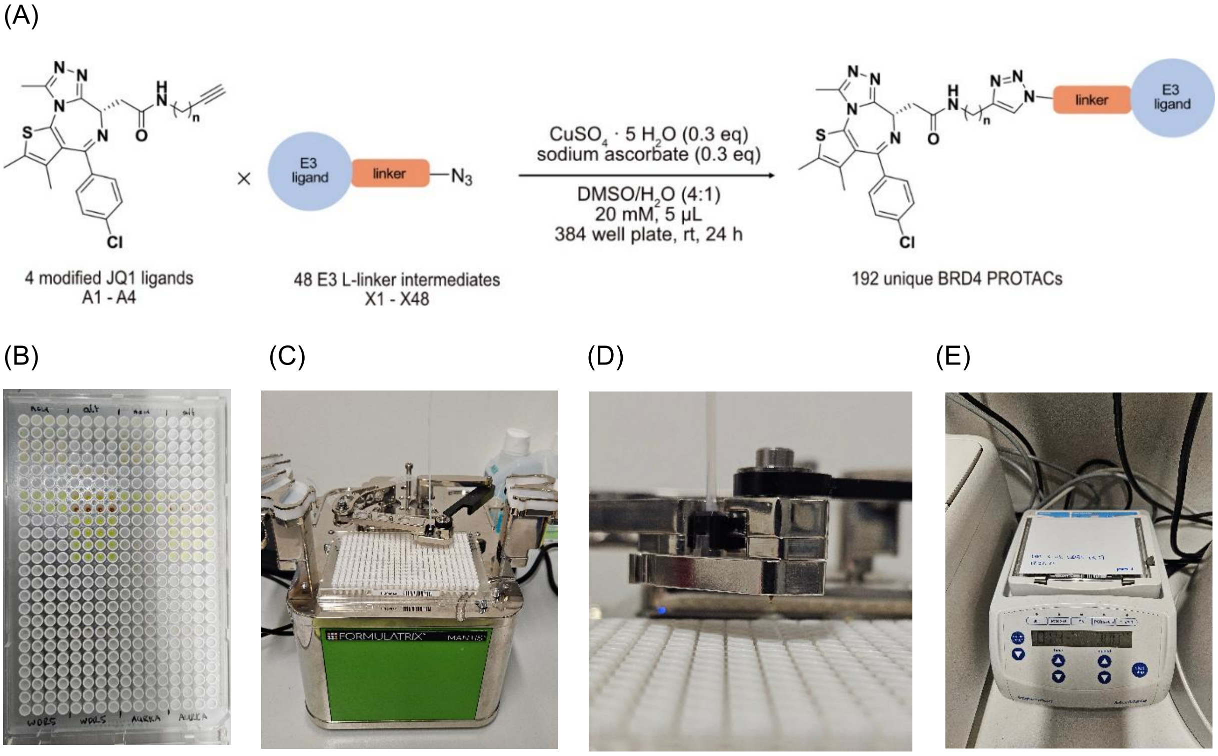


**Figure S1. (Related to Figure 4A). Reaction scheme, conditions and setup for the synthesis of 192 unique BRD4 PROTACs.**

(A) Optimized reaction conditions for the CuAAC chemistry using modified BRD4-targeting ligands A1-A4 and azides X1-48 to obtain 192 unique BRD4 PROTACs. The equivalent of alkyne and azide was set to 1 eq., the equivalent of $CuSO_4{\cdot}5H_2O$ and sodium ascorbate to 0.3 eq., with the reaction temperature at rt and the reaction time at 24 h (plate shaker: 300 rpm). (B) 384 well plate with pipetted azide stock solutions. (C) MANTIS® Liquid Dispenser in action (zoomed out). (D) MANTIS® Liquid Dispenser in action (zoomed in). (E) Plate shaker with attached sealed 384 well plate at 300 rpm.

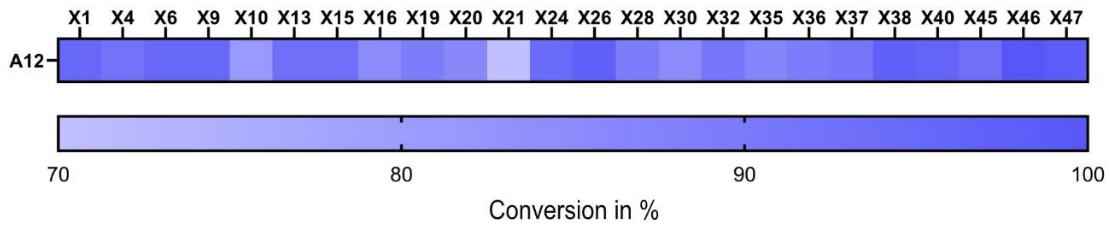


**Figure S2. (Related to Figure 5C). Color-coded heat-map indicating overall conversion rates of CuAAC chemistry of WDR5-targeting ligand A12 and various selected azides.**
Conversion of the CuAAC chemistry for the combination of WDR5-targeting ligand A12 and various selected azides based on HPLC UV data. The average conversion rate amounts to 90.3%.

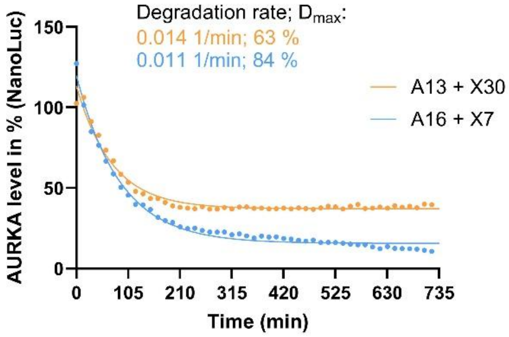


| One phase decay | A16 + X7 | A13 + X30 |
|---|---|---|
| Y0 | 119,9 | 113,3 |
| Plateau | 15,63 | 37,18 |
| K | 0,01136 | 0,01390 |
| Half Life | 61,03 | 49,87 |
| Tau | 88,05 | 71,94 |
| Span | 104,3 | 76,16 |

**Figure S3. (Related to Figure 6E) NanoLuc time dependent live cell measurement monitoring the degradation of AURKA.**
Treatment of MV4-11$^{WDR5\text{-}Nluc(Kless)}$ cells with crude reaction mixtures of PROTACs A16 + X7 and A13 + X30 at a PROTAC concentration of 1 µM. The kinetic measurement was done at 15 min intervals during 12 h at 37 °C. The initial phase of each concentration-dependent degradation curve was fitted using a one-component exponential decay model in GraphPad Prism. From this analysis, the best fit parameters K (degradation rate) and plateau (minimum remaining fraction) were determined.

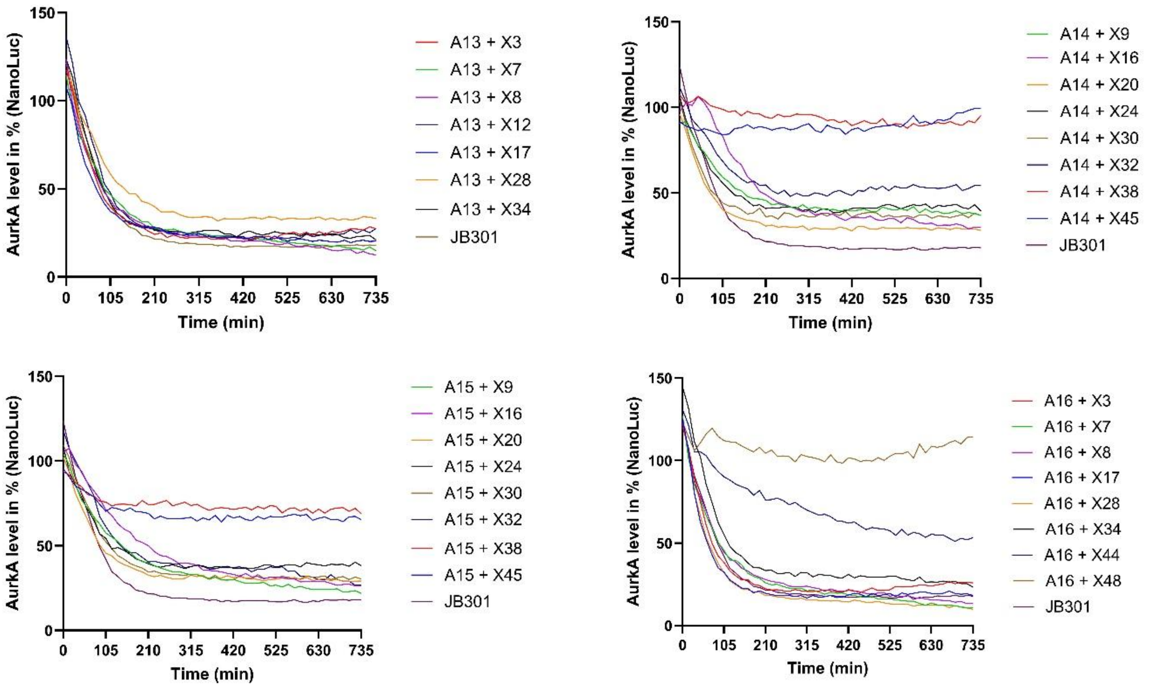


**Figure S4. (Related to Figure 6E) NanoLuc time dependent live cell measurement monitoring the degradation of AURKA.**

Treatment of MV4-11$^{WDR5\text{-}Nluc(Kless)}$ cells with crude reaction mixtures of different PROTACs at a PROTAC concentration of 1 µM. The kinetic measurement was done at 15 min intervals during 12 h at 37 °C. JB301 was used a positive control at a concentration of 150 nM.

| POI | Mass (Da) | protein family | function | subcellular localization |
|---|---|---|---|---|
| BRD4 | 152,219 | bromodomain and extra terminal domain family | Chromatin reader protein; key role in transmission of epigenetic memory | Nucleoplasm |
| sEH | 62,616 | Epoxide hydrolase | Mainly: metabolism of lipid mediators | Cytosol, peroxisome |
| WDR5 | 36,588 | WD-repeat proteins | Contributes to histone modification | Nucleoplasm |
| AURKA | 45,823 | Serine/threonine-protein kinase | Regulation of cell cycle progression | Mainly: centrosome and basal body |

**Table S1. Various properties of target proteins BRD4, sEH, WDR5 and AURKA selected for this study.**

**Frequently Asked Questions S1.**

**Q: Are the used azides bench stable?**
A: Yes, the synthesized azides are stable at rt for at least 30 days but are preferably stored at -20 °C to prevent any decomposition.

**Q: How are the stock solutions prepared and are they stable?**
A: 50 mM stock solutions of alkynes A1-16 and azides X1-48 in DMSO were prepared once and used throughout this study for the optimization of the click reactions as well as the assembly of the PROTACs. All stock solutions were stored at -80 °C and so far (after 8 months), no decomposition of the compounds was observed.

**Q: Are there any major by-products and side-products observed in each click reaction?**
A: No, the CuAAC is usually a very clean reaction, with most of the conversion rates being above 90% for multiple alkynes, based on different POI ligands with varying chemical properties. In some cases though, lower conversion rates were observed with usually one of the reactants being leftover, which is traceable to the imbalance between the amount of alkyne and azide in the respective well. For example when pipetting (either manually or with the MANTIS® Liquid Dispenser), errors do occur, e.g. having a drop of stock solution remaining at the tip of the pipette, that lead to the ratio between alkyne and azide not being precisely 1:1. By minimizing these sort of errors, exquisite conversion rates are achievable with this method.

**Q: Does the purity of azide and alkyne affect the click reaction?**
A: Yes, if one or even both of the reactants are impure or have trace solvents, it becomes difficult to set the ratio between azide and alkyne at 1:1. As mentioned earlier, this leads to lower conversion rates which subsequently compromises the results of the desired D2B screening. Ideally, the reactants should be crystallized solids or amorphous powders, which not only eases up the compound handling while preparing the stock solutions but, more importantly, ensures that the concentration of the stock solutions is as accurate as possible.

**Q: What else to pay attention to when carrying out reactions in a plate based format?**
A: To ensure optimal click reactions at low reaction volumes (< 10 µL), plates with V-bottom wells should be used. Interestingly, click reactions that were performed in flat bottom as well as U-bottom wells were not nearly as efficient and reliable as the ones prepared in plates with V-shaped wells.

**Q: Can the reactions be run more diluted while maintaining high conversion rates?**
A: No, as of now at least. Though we have tried to lower the reaction volume even further (< 5 µL), a significant drop-off in the average conversion rates was observed (Figure 4C). Specifically the consistency and reliability of the click reactions were affected since handling such low volumes accurately has proven to be fairly demanding. When pipetting four different stock solutions (alkyne, azide, $CuSO_4 \cdot 5H_2O$ and sodium ascorbate) a small error can significantly compromise the efficiency of the reaction and handling such low volumes only increases their occurrence, leading to less-consistent conversion rates.

**Data S1. NMR Spectra and LC-MS Spectra, related to compound in Figure 4G**

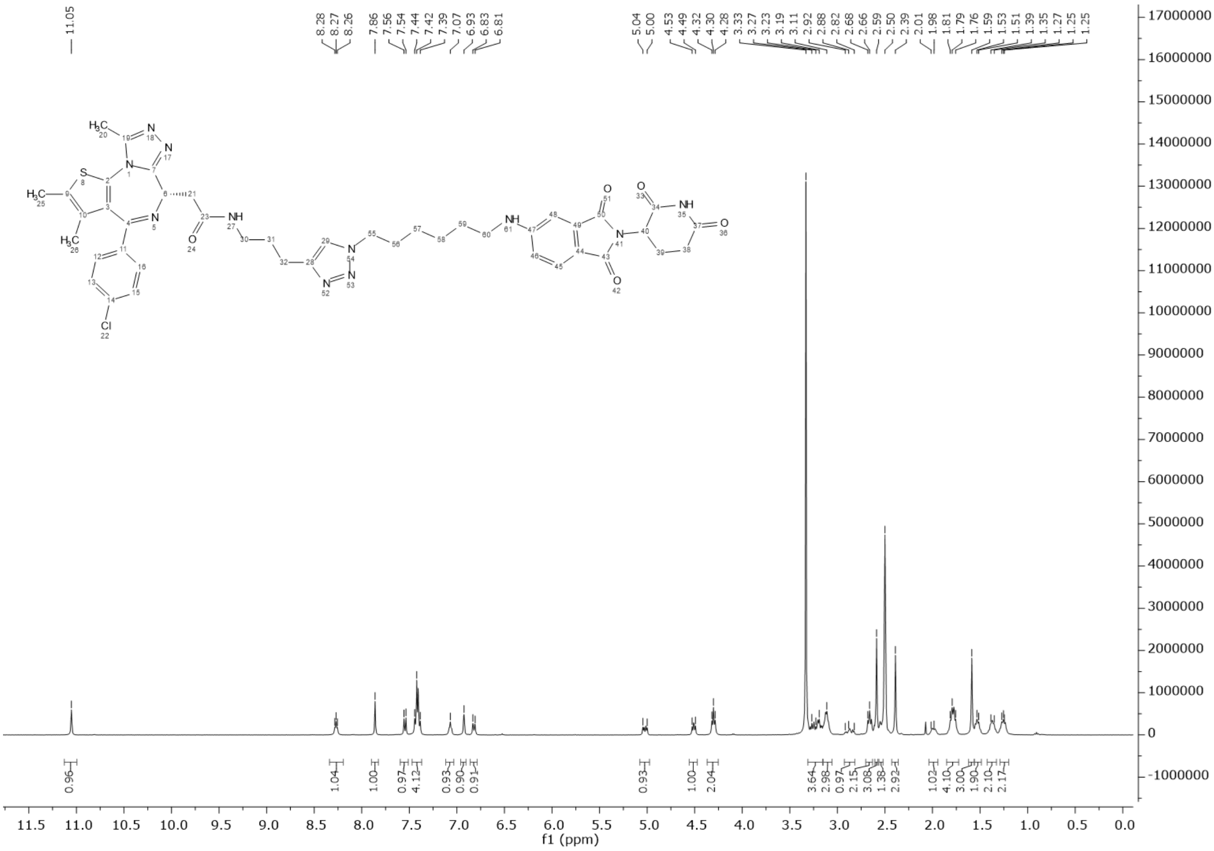

[1]H NMR spectrum of **P1**, related to compound **P1** in Figure 4G

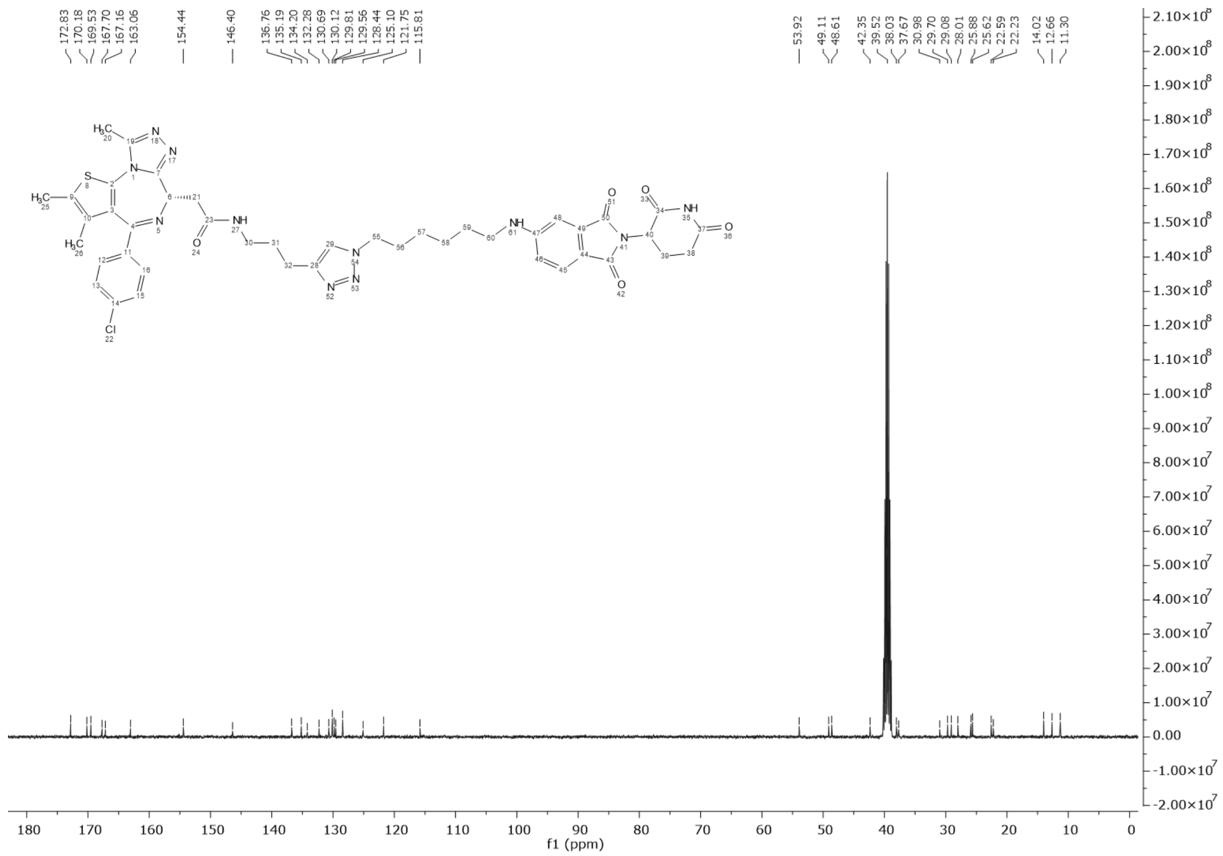

$^{13}$C NMR spectrum of **P1**, related to compound **P1** in Figure 4G

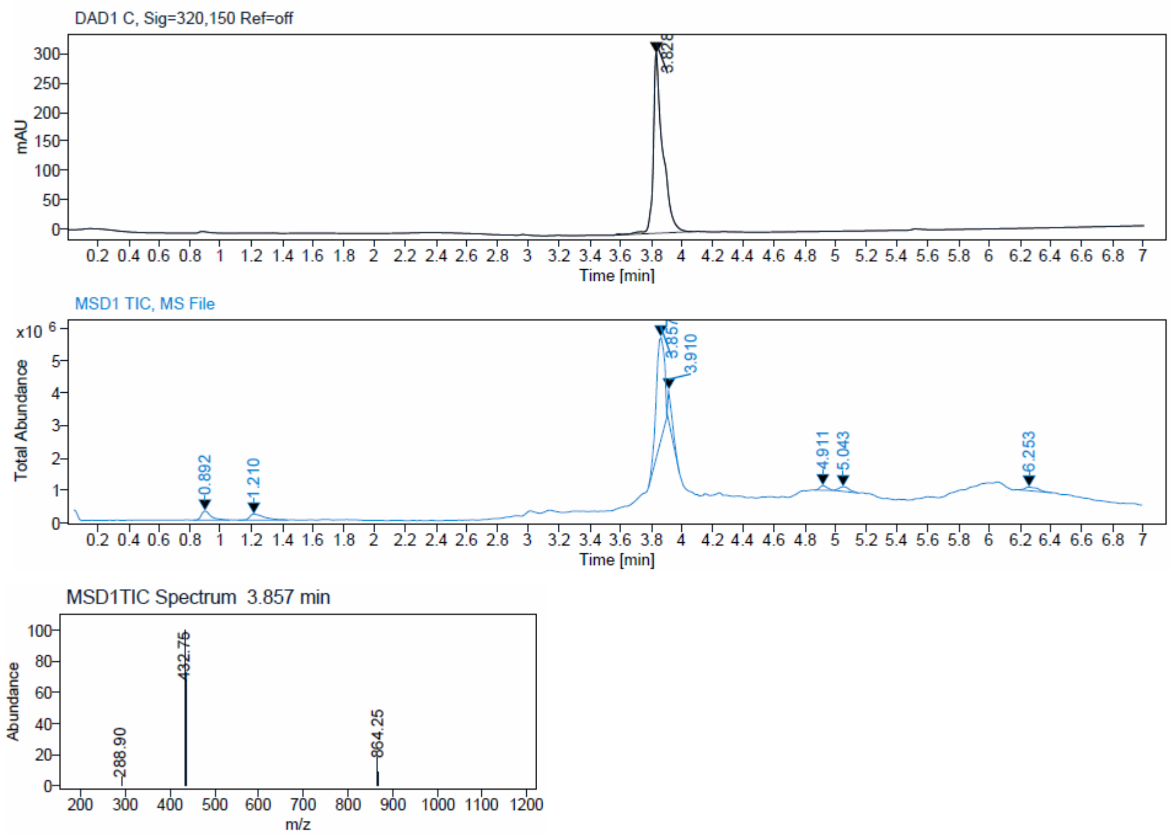


LC-MS spectrum of **P1**, related to compound **P1** in Figure 4G

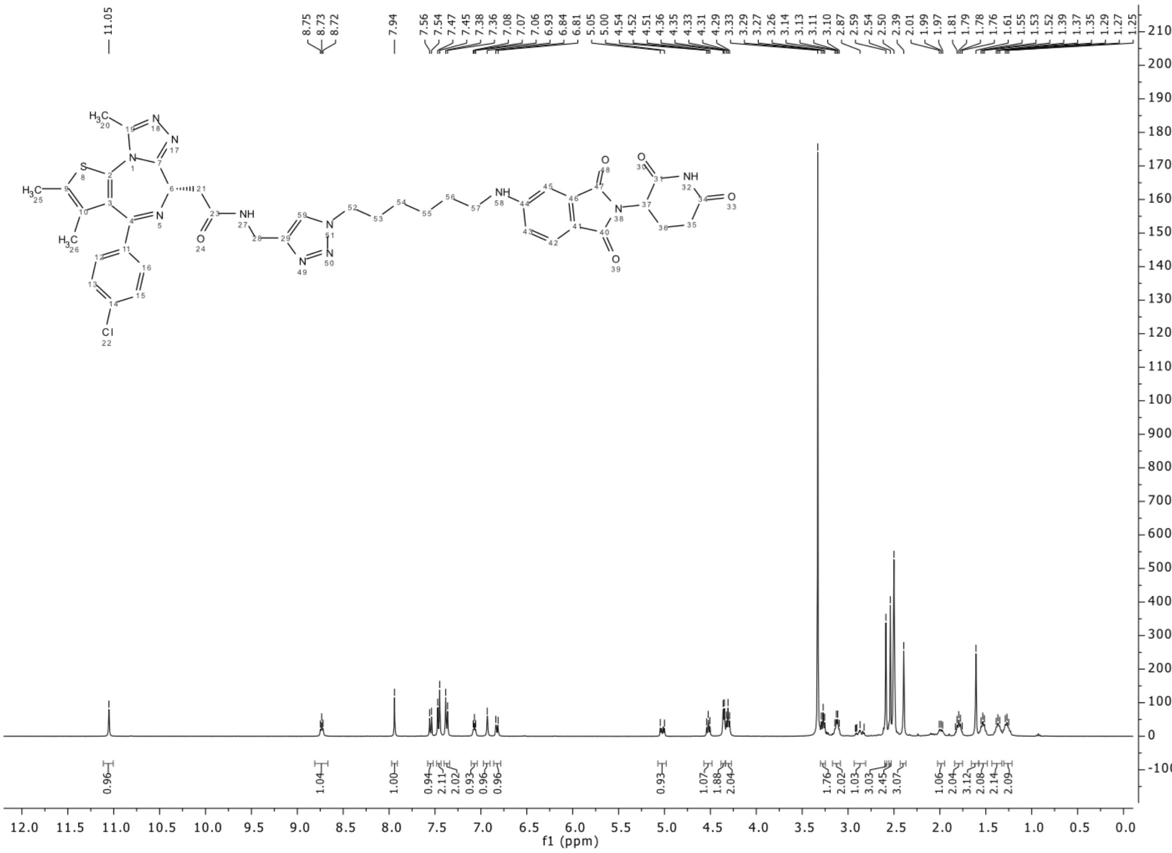

[1]H NMR spectrum of **P2**, related to compound **P2** in Figure 4G

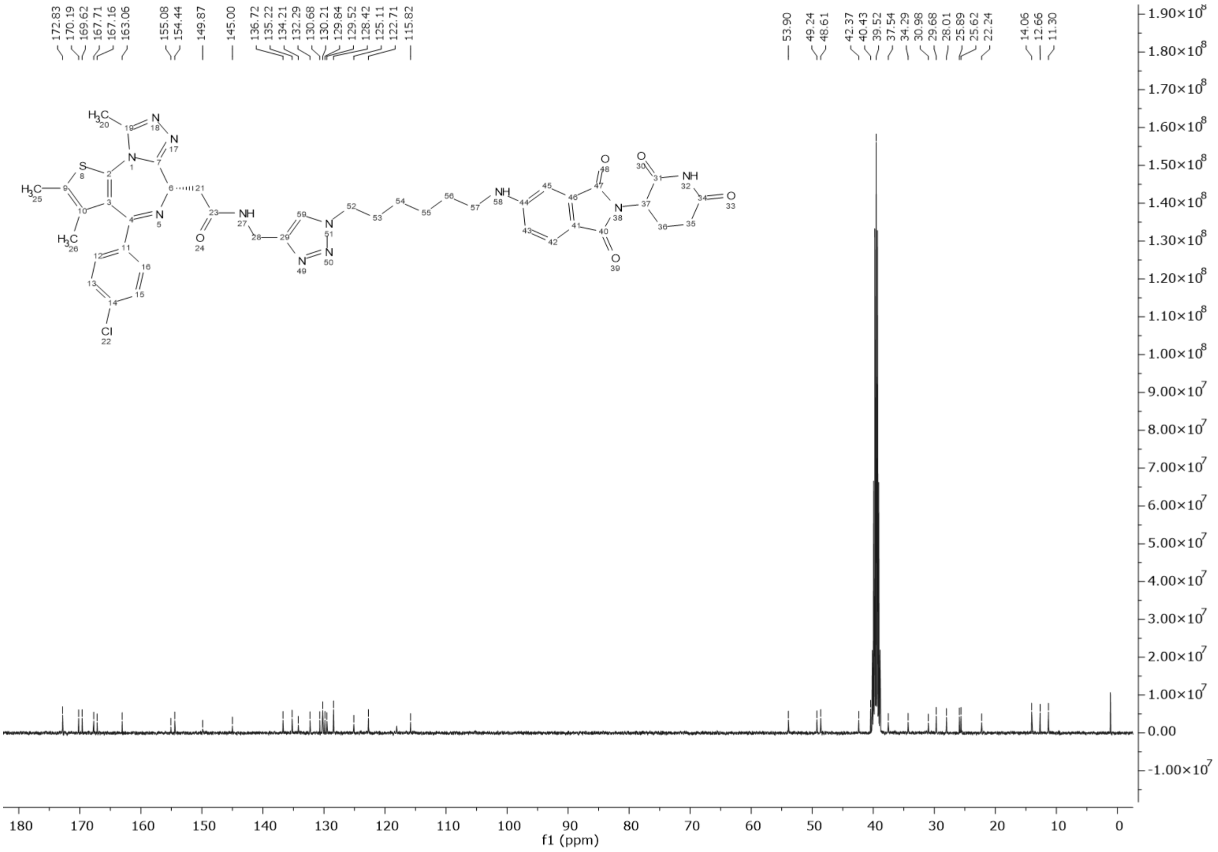

$^{13}$C NMR spectrum of **P2**, related to compound **P2** in Figure 4G

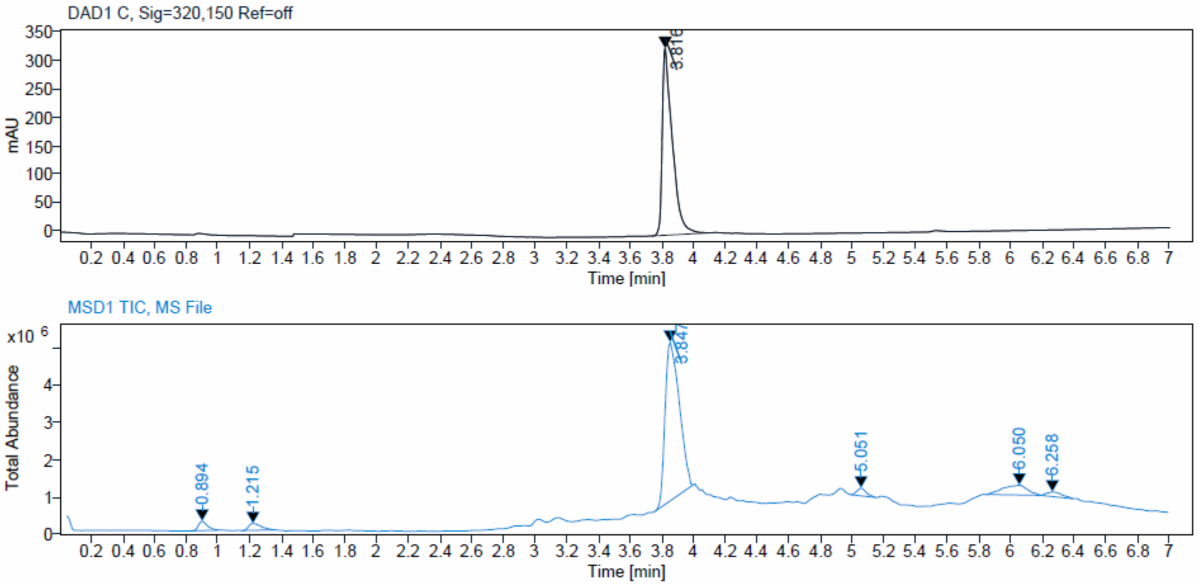

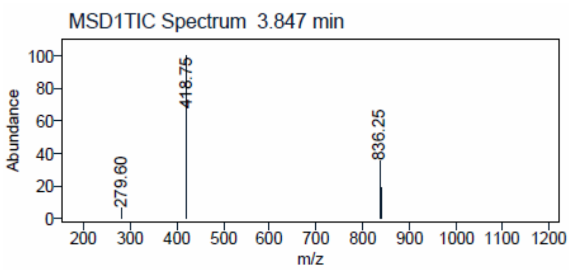


LC-MS spectrum of **P2**, related to compound **P2** in Figure 4G

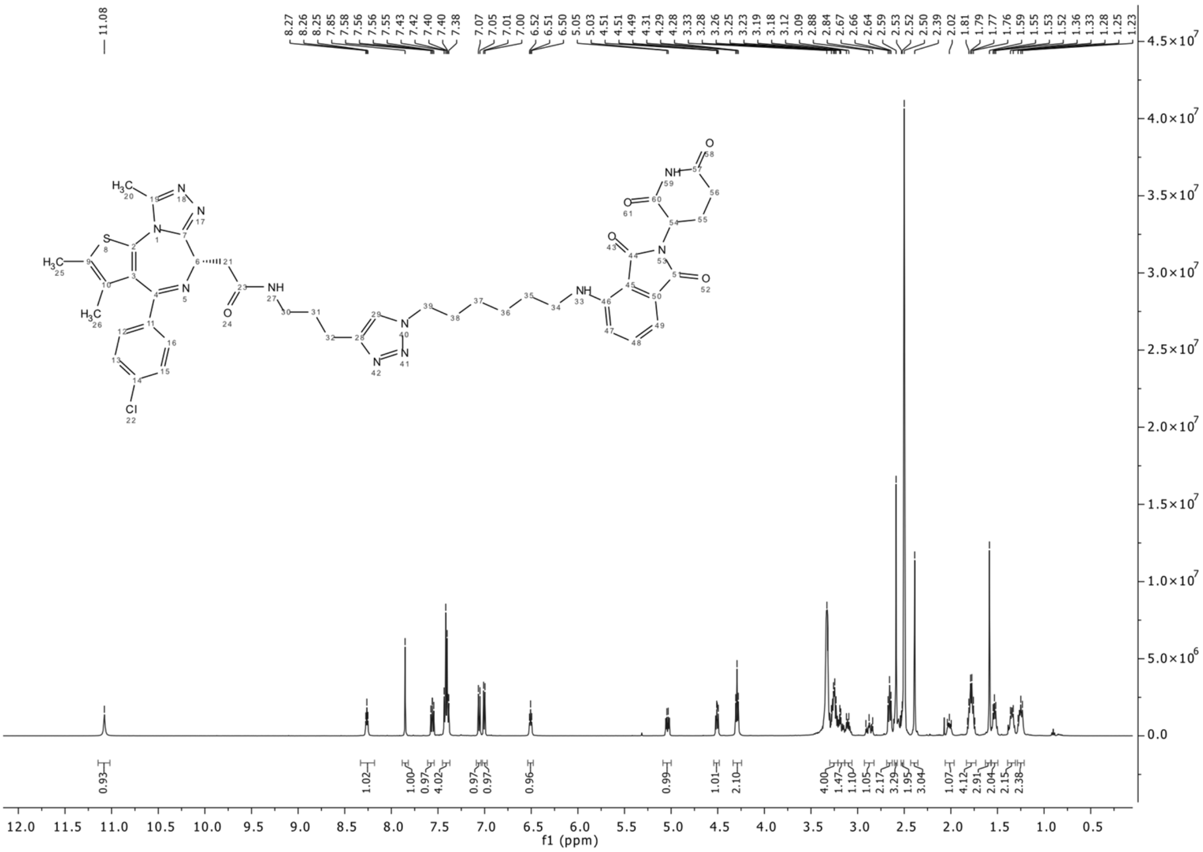


[1]H NMR spectrum of **P3**, related to compound **P3** in Figure 4G

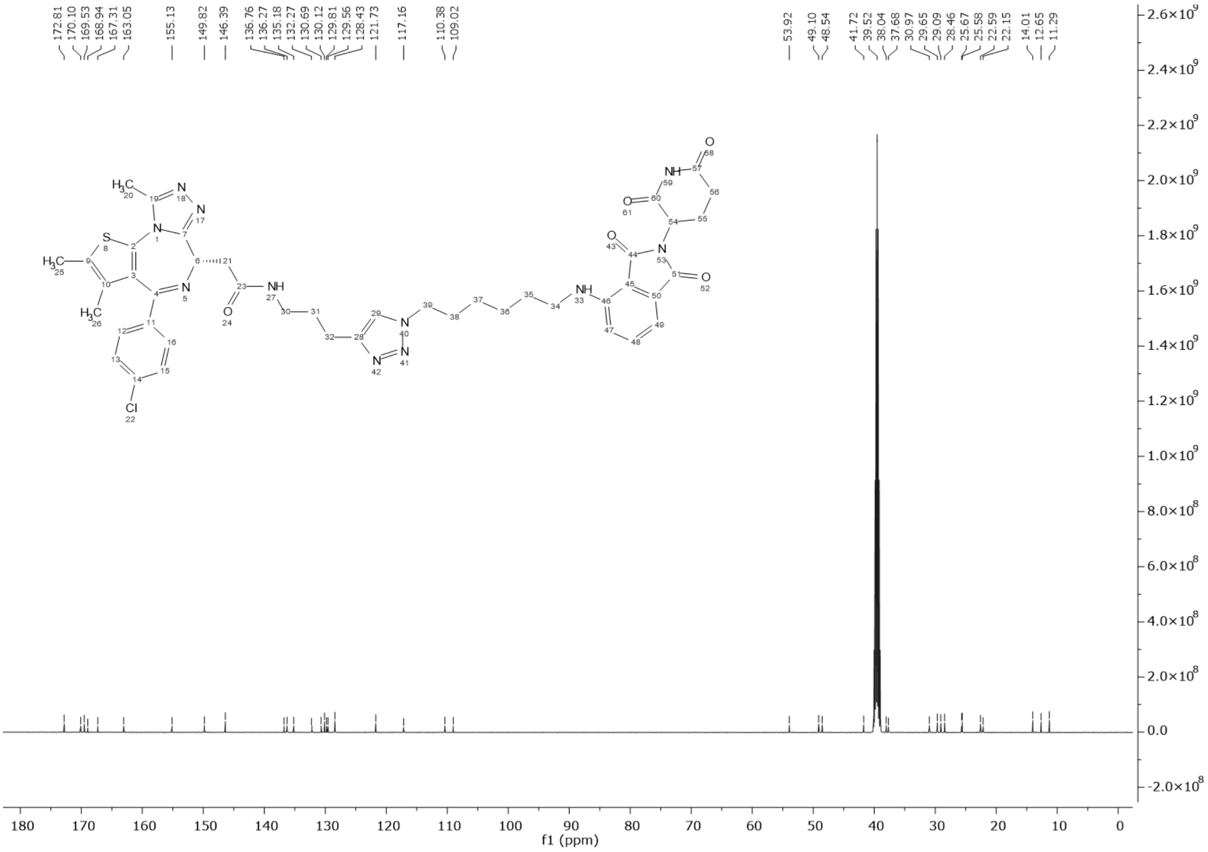


$^{13}$C NMR spectrum of **P3**, related to compound **P3** in Figure 4G

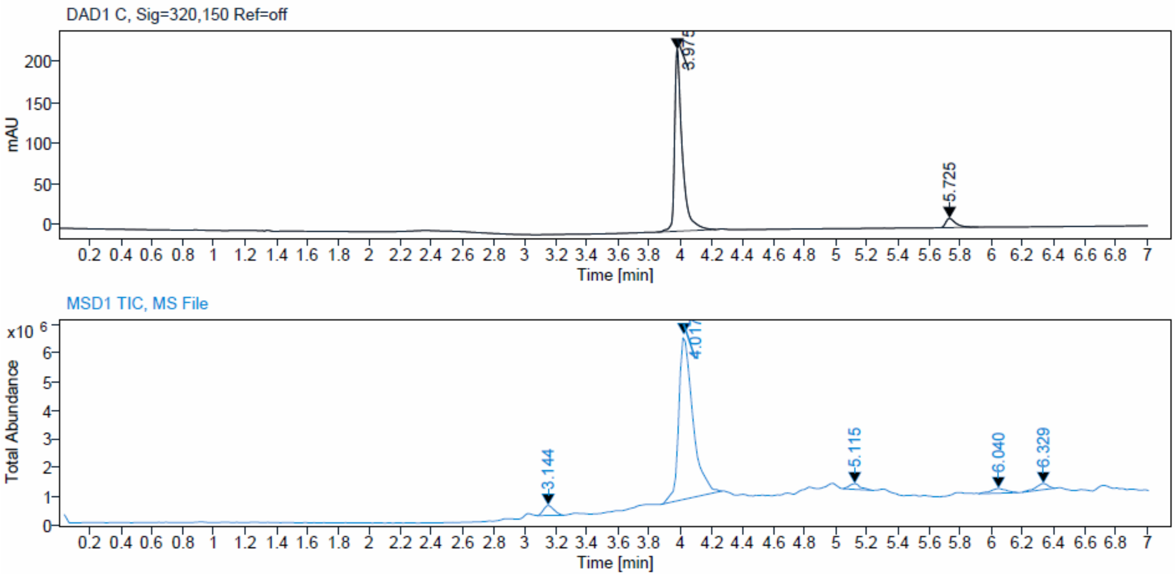

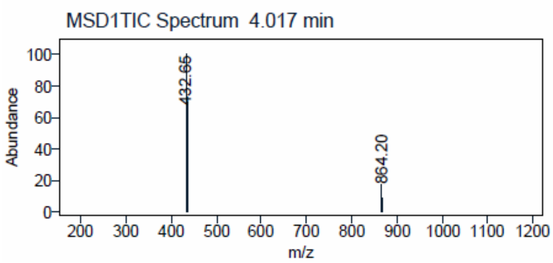


LC-MS spectrum of **P3**, related to compound **P3** in Figure 4G

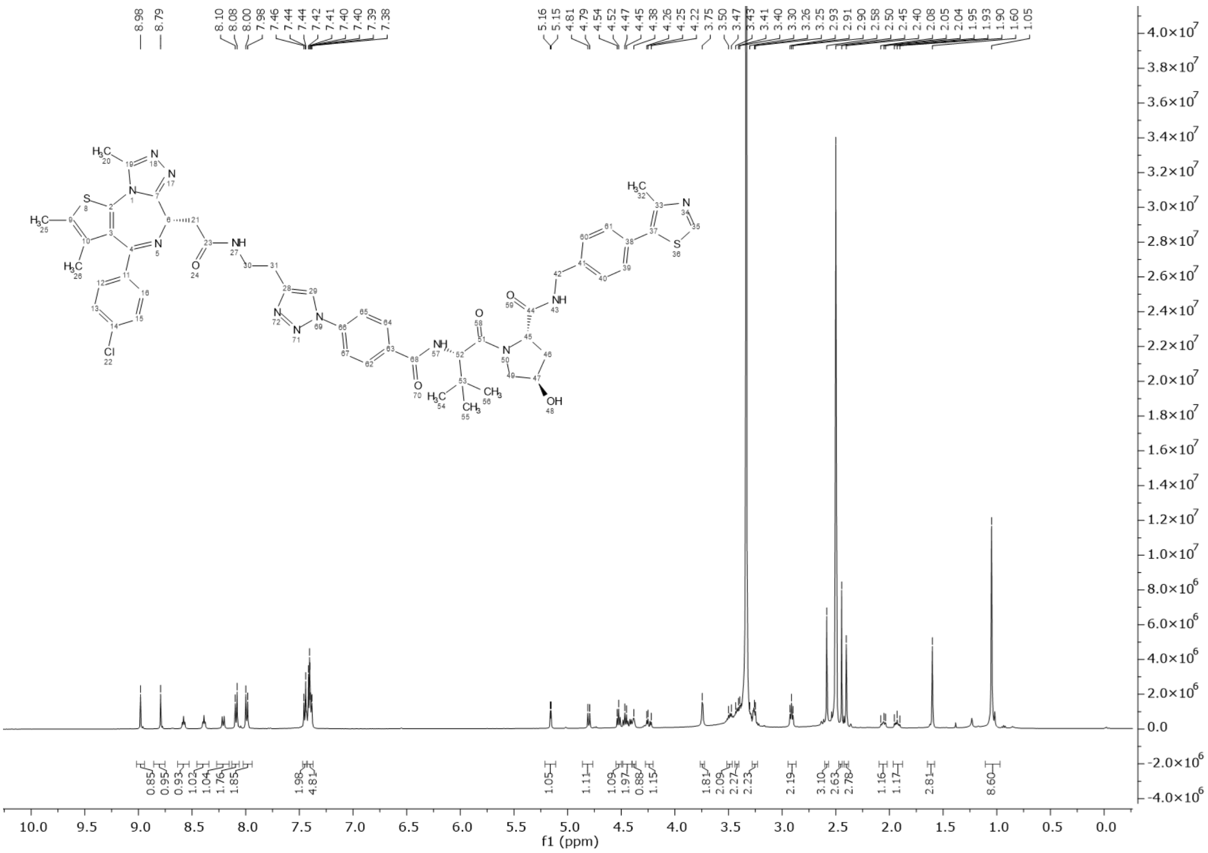

$^{1}$H NMR spectrum of **P4**, related to compound **P4** in Figure 4G

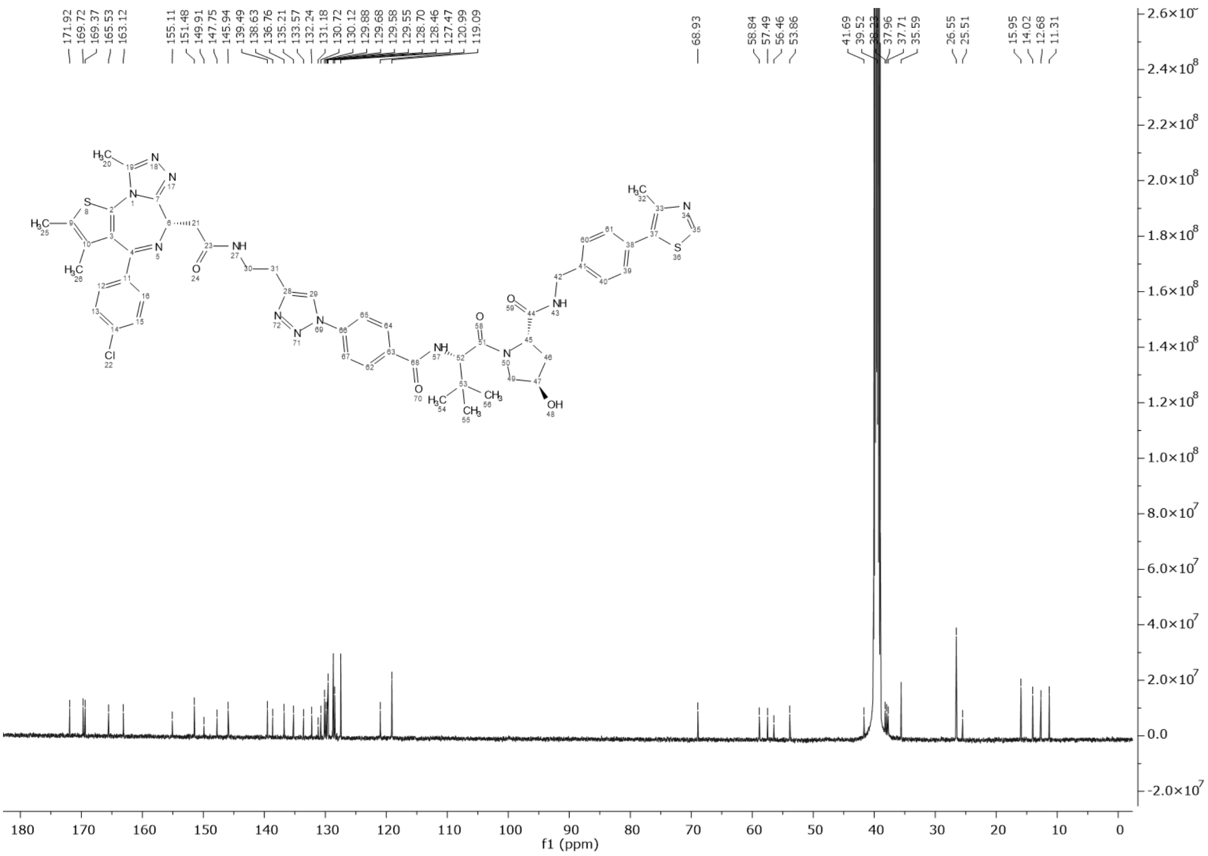

$^{13}$C NMR spectrum of **P4**, related to compound **P4** in Figure 4G

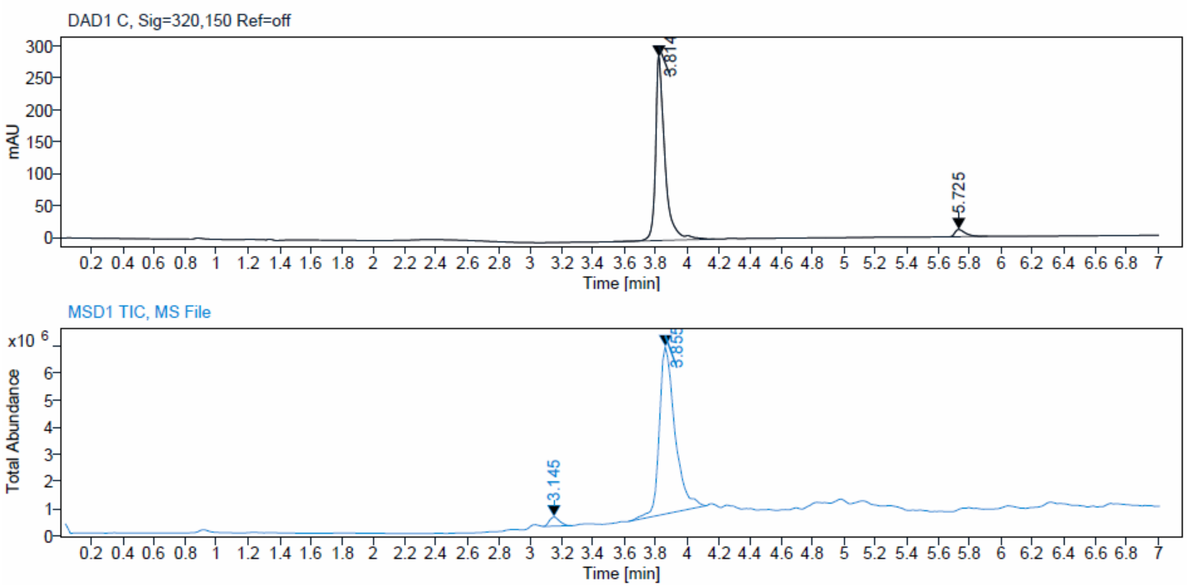

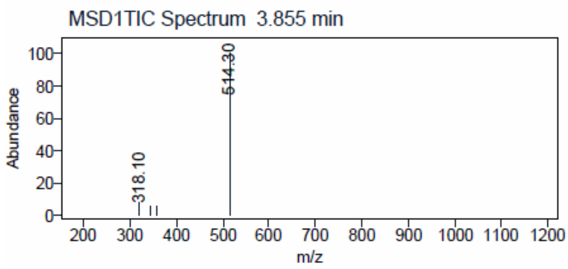


LC-MS spectrum of **P4**, related to compound **P4** in Figure 4G

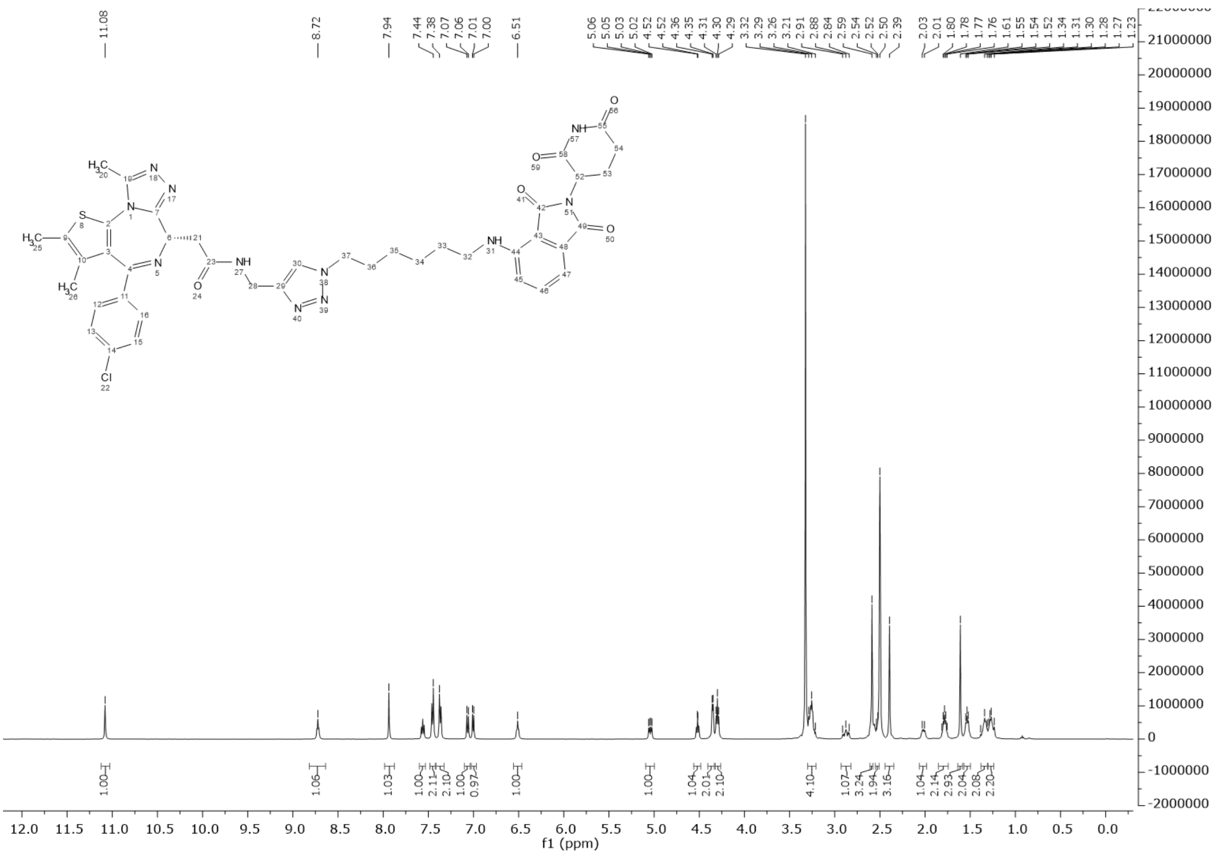


[1]H NMR spectrum of **P5**, related to compound **P5** in Figure 4G

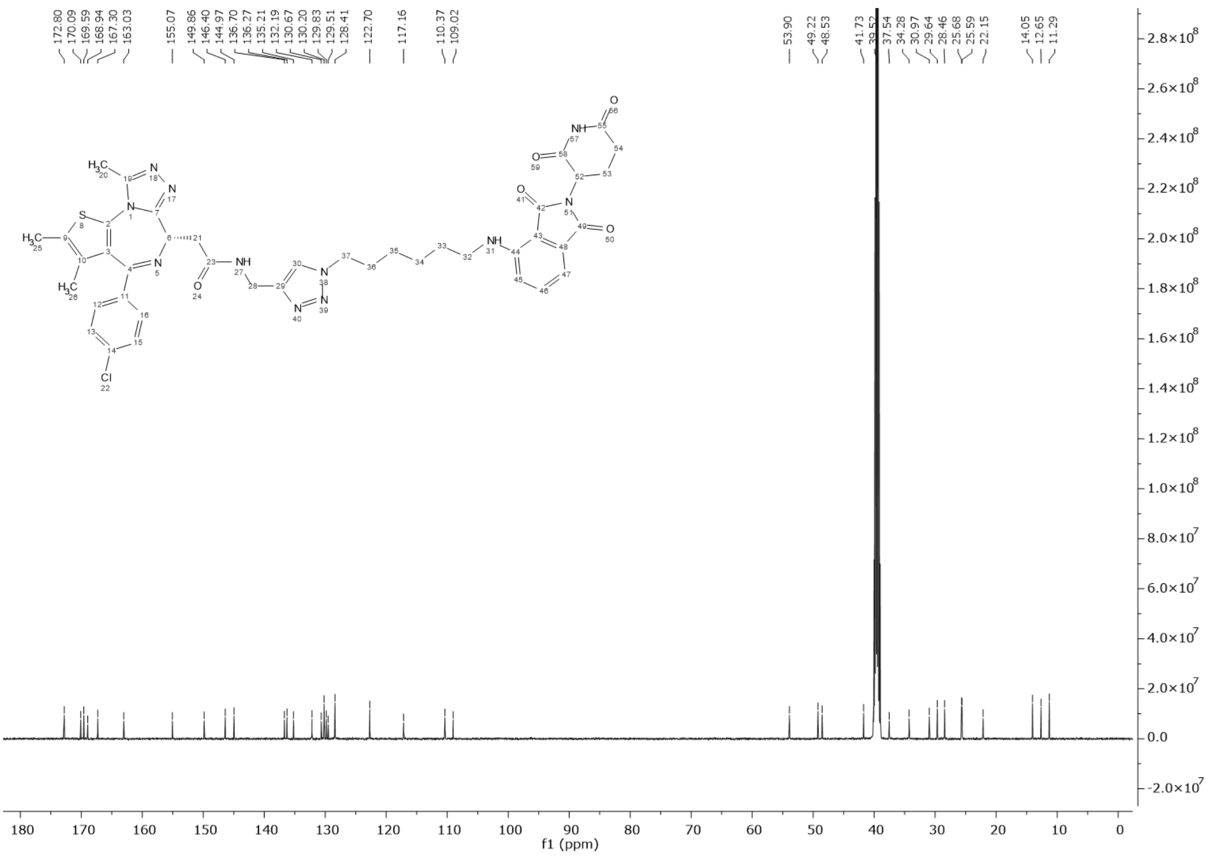


$^{13}C$ NMR spectrum of **P5**, related to compound **P5** in Figure 4G

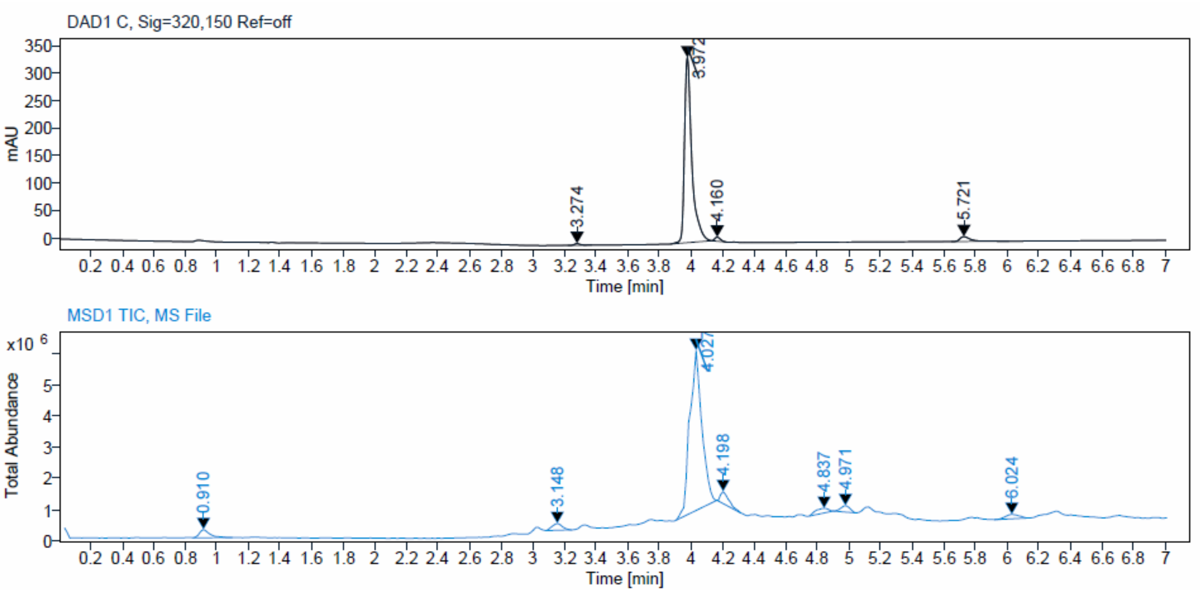

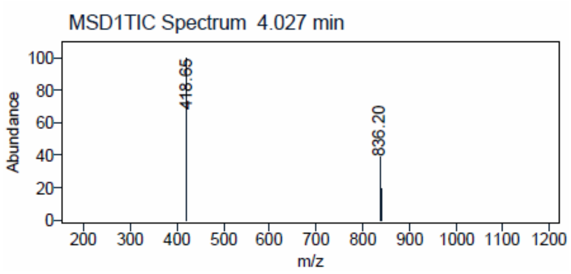


LC-MS spectrum of **P5**, related to compound **P5** in Figure 4G

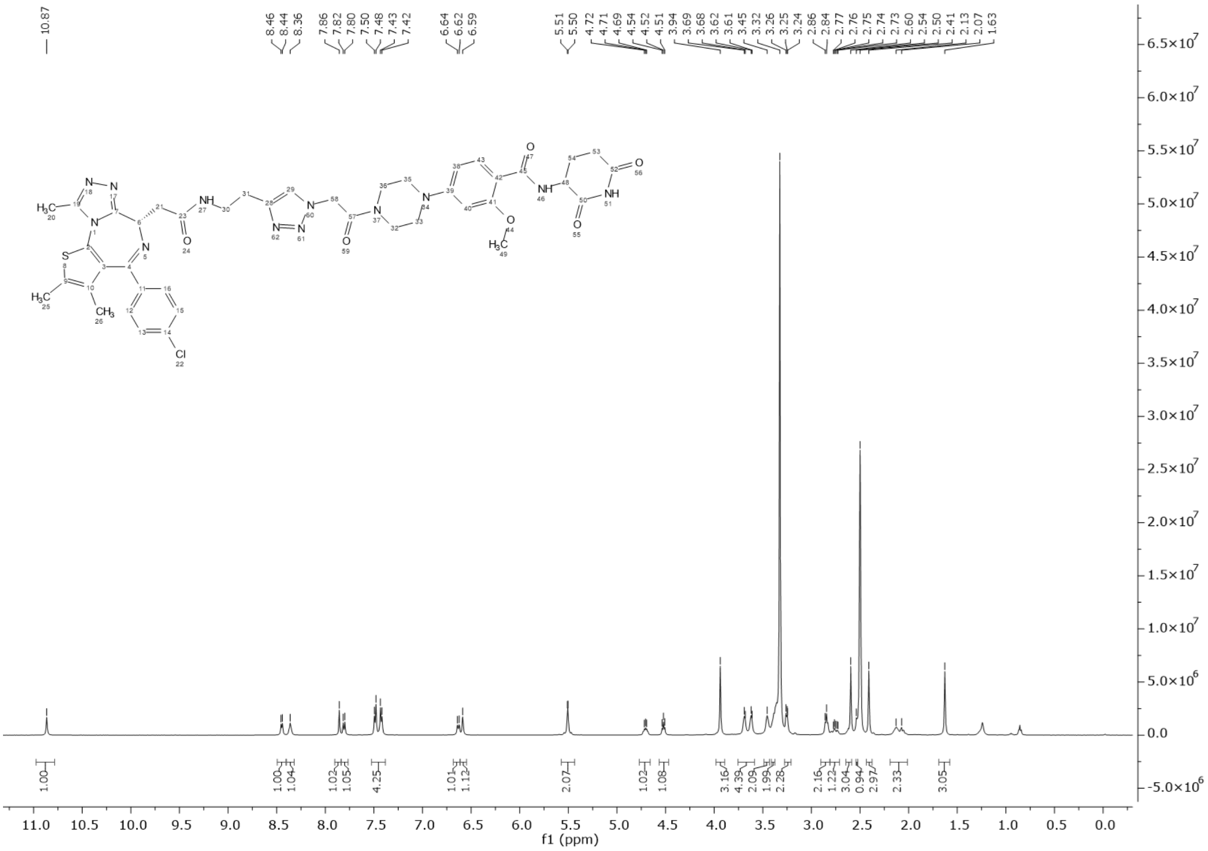


[1]H NMR spectrum of **P6**, related to compound **P6** in Figure 4G

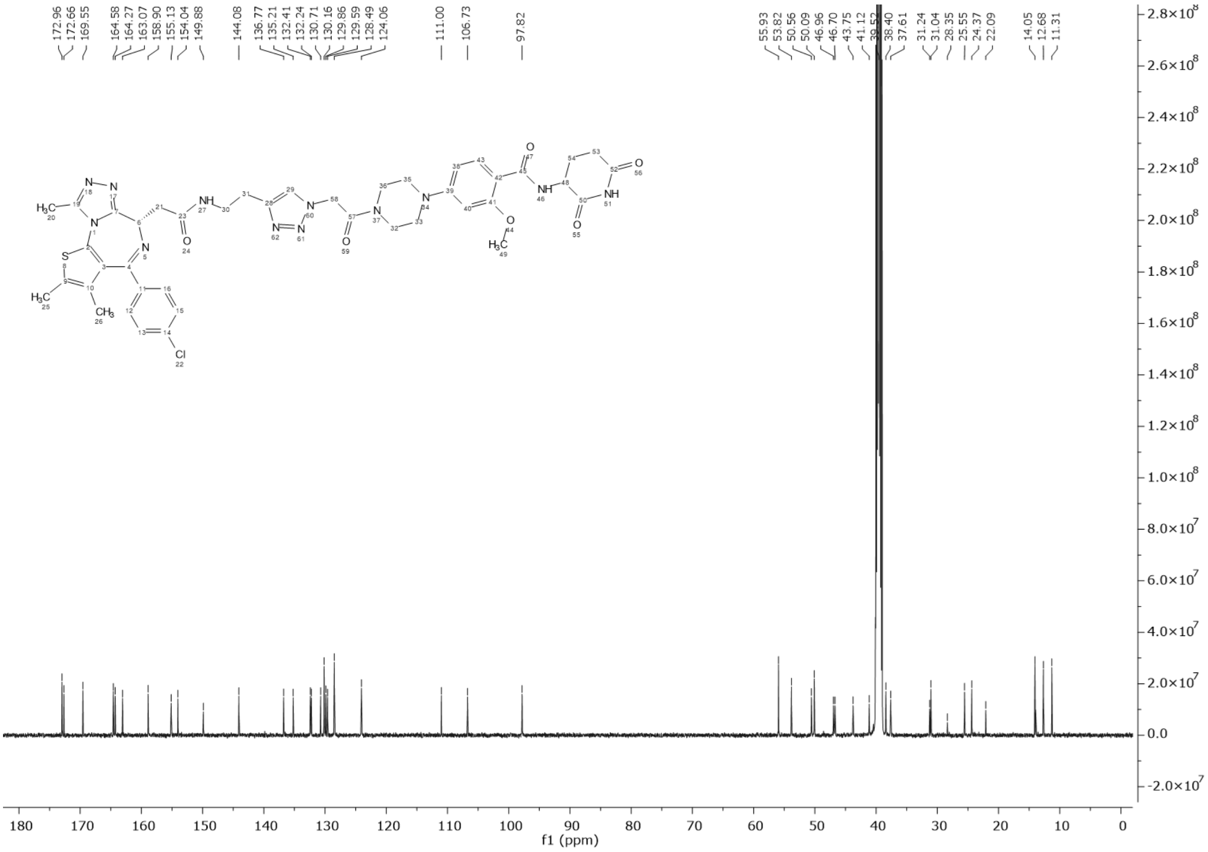

$^{13}$C NMR spectrum of **P6**, related to compound **P6** in Figure 4G

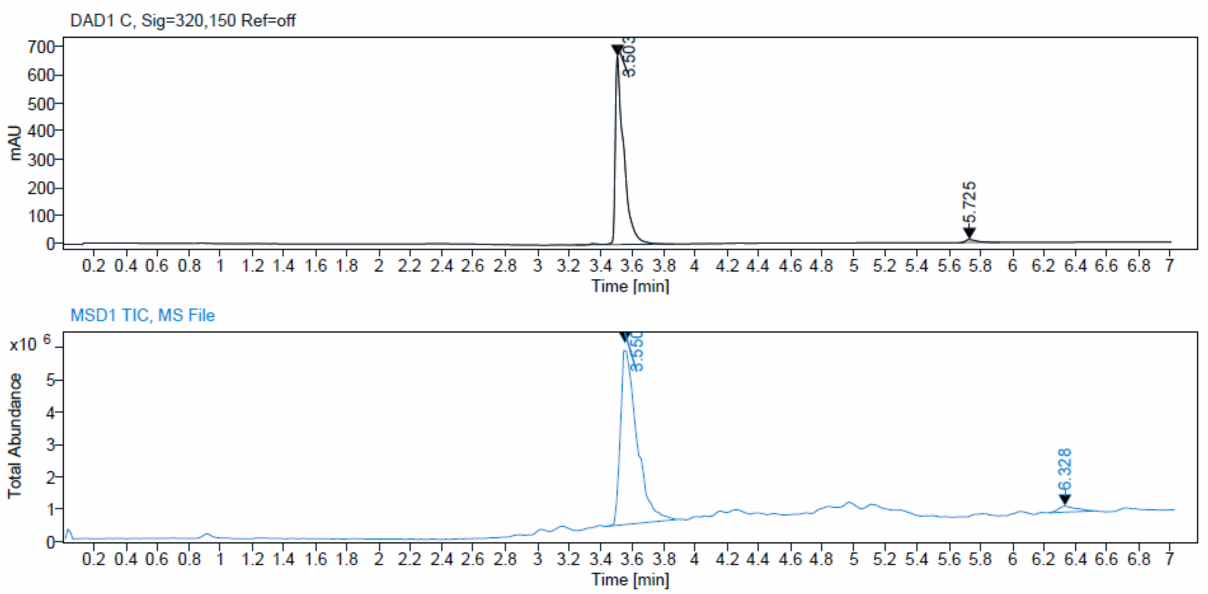

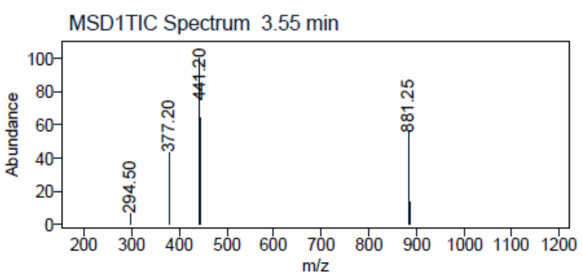


LC-MS spectrum of **P6**, related to compound **P6** in Figure 4G

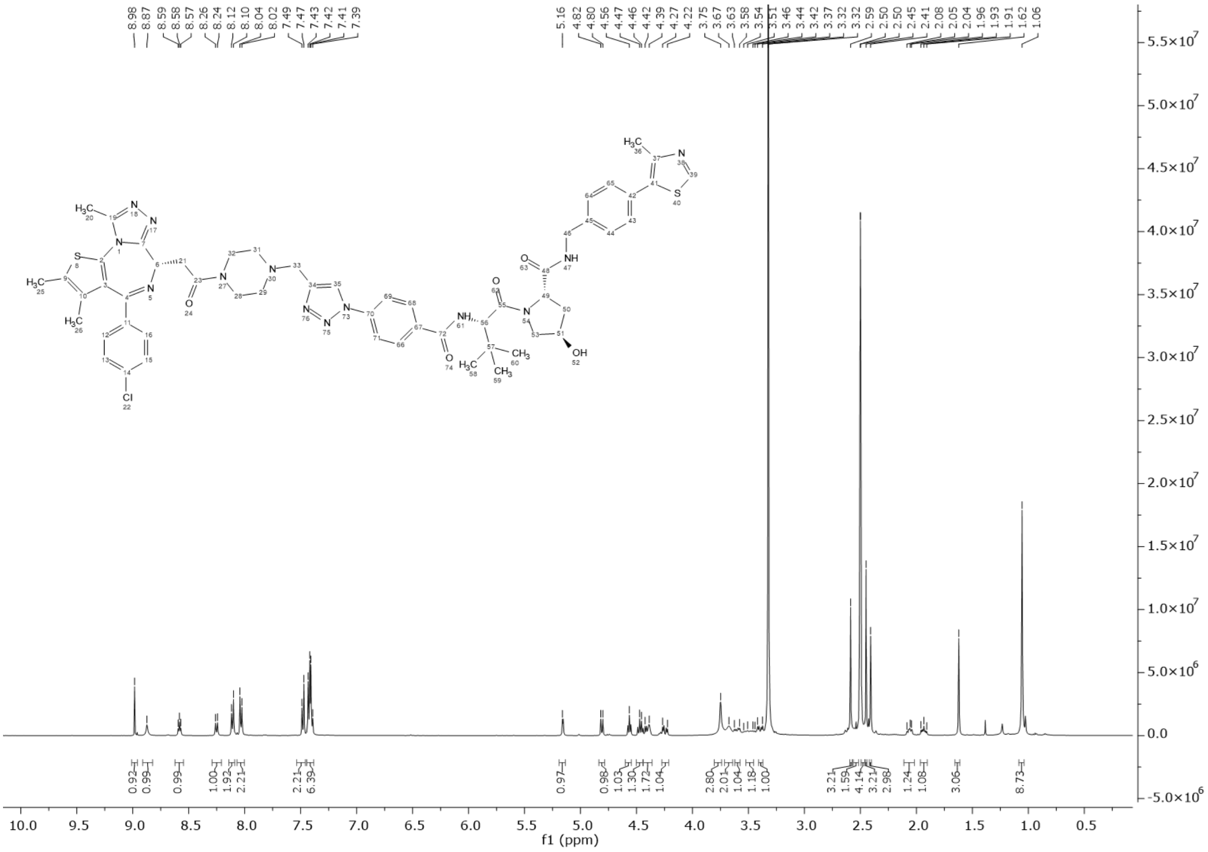


$^1$H NMR spectrum of **P7**, related to compound **P7** in Figure 4G

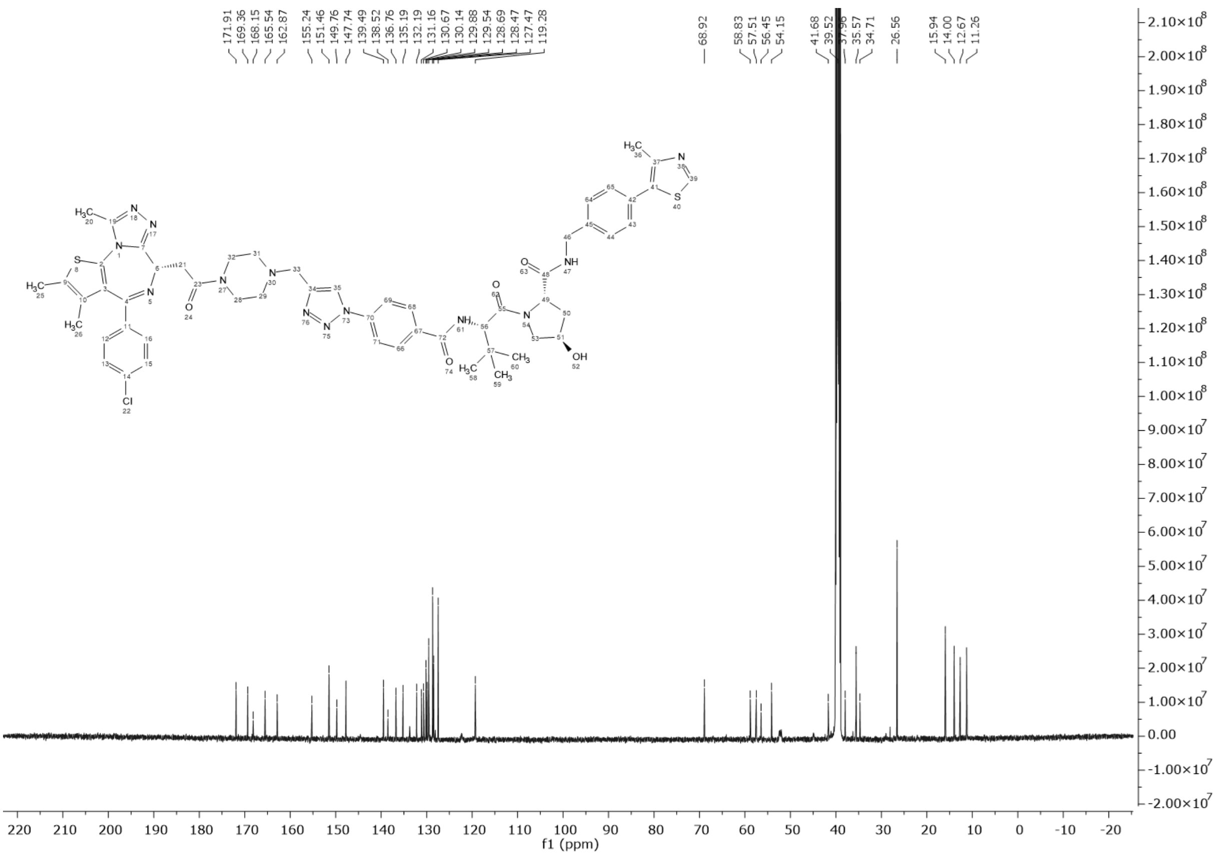

$^{13}$C NMR spectrum of **P7**, related to compound **P7** in Figure 4G

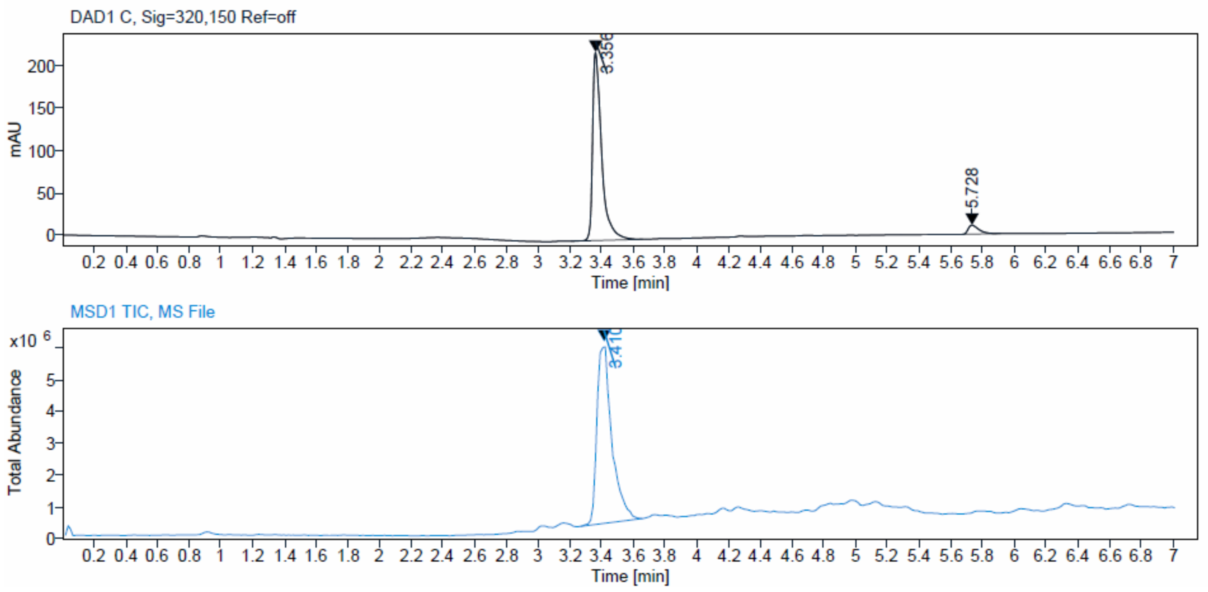

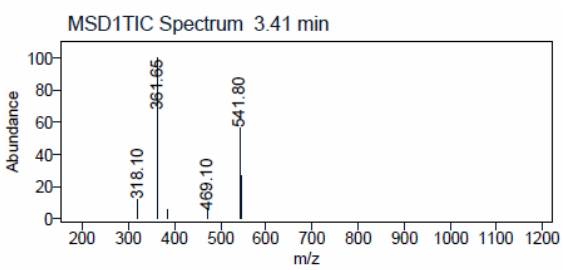


LC-MS spectrum of **P7**, related to compound **P7** in Figure 4G

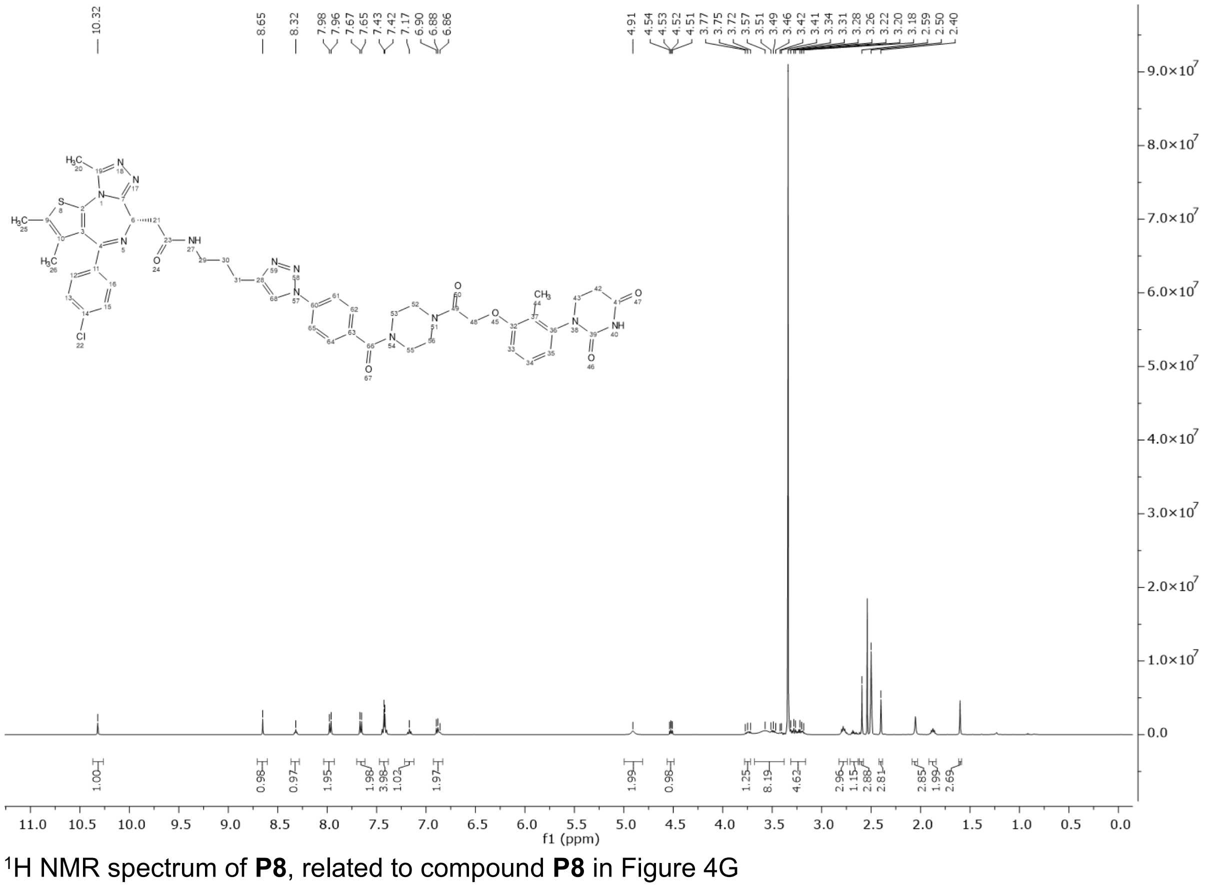


[1]H NMR spectrum of **P8**, related to compound **P8** in Figure 4G

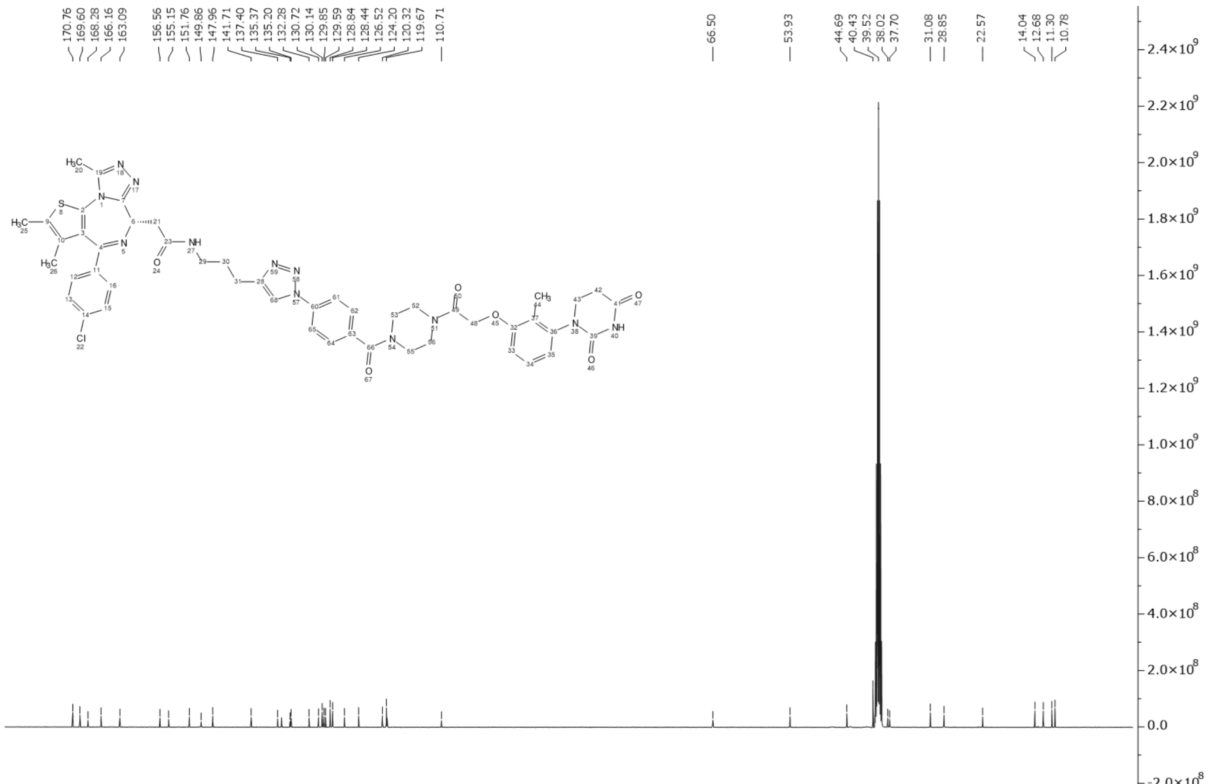

$^{13}C$ NMR spectrum of **P8**, related to compound **P8** in Figure 4G

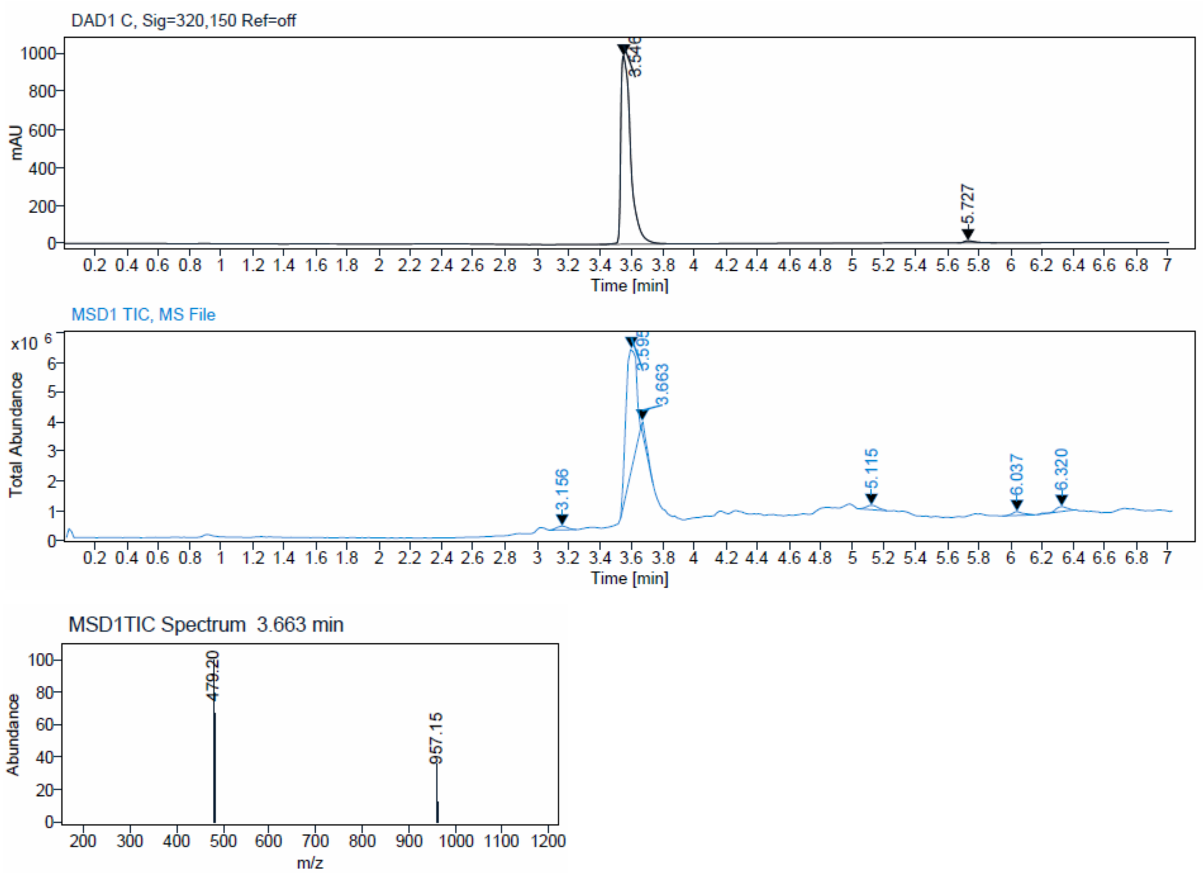


LC-MS spectrum of **P8**, related to compound **P8** in Figure 4G
